# Multiple probabilistic models extract features from protein sequence data and resolve functional diversity of very different protein families

**DOI:** 10.1101/717249

**Authors:** R. Vicedomini, J.P. Bouly, E. Laine, A. Falciatore, A. Carbone

**Affiliations:** Sorbonne Université, CNRS, IBPS, Laboratoire de Biologie Computationnelle et Quantitative - UMR 7238, 4 place Jussieu, 75005 Paris, France; Sorbonne Université, Institut des Sciences du Calcul et des Données; CNRS, Sorbonne Université, Institut de Biologie Physico-Chimique, Laboratory of Chloroplast Biology and Light Sensing in Microalgae - UMR7141, Paris, France; Institut Universitaire de France, Paris 75005, France

**Keywords:** genome, metagenome, functional classification, protein classification, probabilistic model, profile, cryptochrome, photolyase, photoreceptor, WW domain, glycoside hydrolase, Radical SAM, Haloacid Dehalogenase, B12-binding domain containing, methylthiotransferase, SPASM/twitch domain containing

## Abstract

Sequence functional classification has become a critical bottleneck in understanding the myriad of protein sequences that accumulate in our databases. The great diversity of homologous sequences hides, in many cases, a variety of functional activities that cannot be anticipated. Their identification appears critical for a fundamental understanding of living organisms and for biotechnological applications.

ProfileView is a sequence-based computational method, designed to functionally classify sets of homologous sequences. It relies on two main ideas: the use of multiple probabilistic models whose construction explores evolutionary information in available databases, and a new definition of a representation space where to look at sequences from the point of view of probabilistic models combined together. ProfileView classifies families of proteins for which functions should be discovered or characterised within known groups.

We validate ProfileView on seven classes of widespread proteins, involved in the interaction with nucleic acids, amino acids and small molecules, and in a large variety of functions and enzymatic reactions. ProfileView agrees with the large set of functional data collected for these proteins from the literature regarding the organisation into functional subgroups and residues that characterize the functions. Furthermore, ProfileView resolves undefined functional classifications and extracts the molecular determinants underlying protein functional diversity, showing its potential to select sequences towards accurate experimental design and discovery of new biological functions.

ProfileView proves to outperform three functional classification approaches, CUPP, PANTHER, and a recently developed neural network approach based on Restricted Boltzmann Machines. It overcomes time complexity limitations of the latter.

## 1 Introduction

The functional classification of biological sequences has become a fundamental bottleneck to the understanding of the ever-increasing genomic and metagenomic sequence data accumulating in our databases. This quest depends on the correct domain annotation of coding genes (Ponting and Dickens, 2001; Prakash and Taylor, 2012; De Filippo *et al*., 2012), which, in the past, was handled by sequence homology-, and feature-based approaches.

The first and most intuitive approach searches for homologous sequences to already known protein or domain sequences (Hawkins *et al*., 2006; Wass and Sternberg, 2008; Loewenstein *et al*., 2009; Clark and Radivojac, 2011; Törönen *et al*., 2018) and does it either by a direct pairwise sequence alignment or by passing through protein signatures, which are descriptions of protein or domain families derived from multiple sequence alignments. It is based on the “orthology-function conjecture” for which orthologues carry out biologically equivalent functions in different organisms, in contrast to paralogues whose functions typically diverge after duplication (Gabaldón and Koonin, 2013). Due to complex processes of evolution, many homologues diversified their functions and the sequence homology approach should be applied with great awarness: different similarity levels in homology should induce different levels in functional annotation transfer. This represents a serious pitfall for the approach. A second pitfall, is linked to the production of probabilistic models, describing conserved characteristics across sequences. Indeed, these families might be made of a few members very divergent from each others (rare) or of a continuum of thousands of sequences due to a lack of functional/evolutionary pressure, which challenges the family definition and produces super-family/clan totally degenerated models (most frequent) of restrained use.

The second class of methods is based on the selection of an appropriate set of features (like short sequence segments or wavelet decompositions) (Karchin *et al*., 2005; Wen *et al*., 2005; Wan and Jones, 2020; Bonetta and Valentino, 2020). Other computational schemas use protein structure (Pazos and Sternberg, 2004; Pal and Eisenberg, 2005; Lee *et al*., 2007; Dawson *et al*., 2017), phylogenetics and evolutionary relationships (Eisen, 1998; Engelhardt *et al*., 2005, 2011; Gaudet *et al*., 2011; Sahraeian *et al*., 2015; Gumerov and Zhulin, 2020), interaction and association data (Deng *et al*., 2002; Vazquez *et al*., 2003; Letovsky and Kasif, 2003; Nabieva *et al*., 2005; Sharan *et al*., 2007; Cao *et al*., 2014; Pham and Lichtarge, 2020) and a combination of those (Shin *et al*., 2007; Furnham *et al*., 2012; Boari de Lima *et al*., 2016; Cao and Cheng, 2016; Zhang *et al*., 2017; Kulmanov and Hoehndorf, 2020), with the evident dependence on the availability of different data-types and a large and very diversified dataset of sequences.

Novel computational approaches classifying sequences by function and overcoming the limitations intrinsic to existing methods would help screening sequences to design accurate experiments directed to functional testing and to discover new functions. ProfileView was conceived for this purpose.

ProfileView is a computational method able to classify hundreds/thousands of homologous sequences into functional groups. It is strongly based on the understanding of the structure of the sequence data imposed by the evolutionary history of the sequences. The first main step of ProfileView is to encode functional and structural information belonging to the protein family into multiple probabilistic models that capture the diversity of the homologous sequences in the family. Based on the set of different models for the family, the second main step of ProfileView is to define an original sequence space which organises sequences by function. Biologically interpretable information and functional motifs are extracted from the classification process. That is, the family members are organised in a tree structure, where subfamily delineations are possible thanks to the hierarchical organisation. The presence of multiple functions in a family or subfamily makes it desirable to subdivide its members into smaller groups in order to capture the differences in function-related features at a level lower than the subfamily. ProfileView representative models and their specific conserved motifs proved to be good indicators of this functional delineation. ProfileView can be applied on a large scale on very diverse datasets.

In the past, the usage of multiple probabilistic models demonstrated to be powerful in the context of domain annotation (Bernardes *et al*., 2016; Ugarte *et al*., 2018), where they showed to be highly accurate on full genomes and metagenomic/metatranscriptomic datasets, allowing for the discovery of new sequences enriching protein families (Fortunato *et al*., 2016; Amato *et al*., 2017). Here, these models are not used to discover homologous sequences but to capture the variety of functional motifs characterizing a protein family. Their construction demands a relatively small number of sequences (a minimum of 20), and therefore, they can encode even functional motifs that are poorly represented in sequence space, generating a possibly very large motifs diversification.

To highlight its power and generality, we applied ProfileView to seven protein families whose members are characterised by a large functional diversity, multiple members are functionally well-characterised proteins and subfamilies delineations have been validated experimentally together with their functional motifs: the Cryptochrome/Photolyase Family (CPF), the WW domains, the glycoside hydrolase enzymes GH30 family and four protein subgroups belonging to two enzyme superfamilies, the Haloacid Dehydrogenase (HAD/*β*-PGM/Phosphatase-like subgroup) and the Radical SAM (B12-binding domain containing, Methylthiotransferase and SPASM/twitch domain containing). These families and subgroups allowed us to demonstrate the power in feature extraction, the simplicity in the interpretability of the results and the methodological approach, and the computational efficiency of ProfileView compared to a recent artificial neural networks approach to sequence classification (Tubiana *et al*., 2019). Comparisons are also made with the PANTHER classification system (Mi *et al*., 2012, 2013) and the CUPP platform (Barrett and Lange, 2019). For each protein family, ProfileView agrees with all available experimental data. Many homologous protein sequences yet to be classified were classified by ProfileView in this work.

## Results

### Converting sequences in multidimensional vectors with probabilistic models

Our methodological approach to sequence classification, ProfileView, is outlined hereafter and illustrated in **Fig. 1**. ProfileView takes as input a set of homologous sequences and a protein domain, and returns a classification of the sequences in functional subgroups together with functional motifs characterising the subgroups.

**Figure 1:**
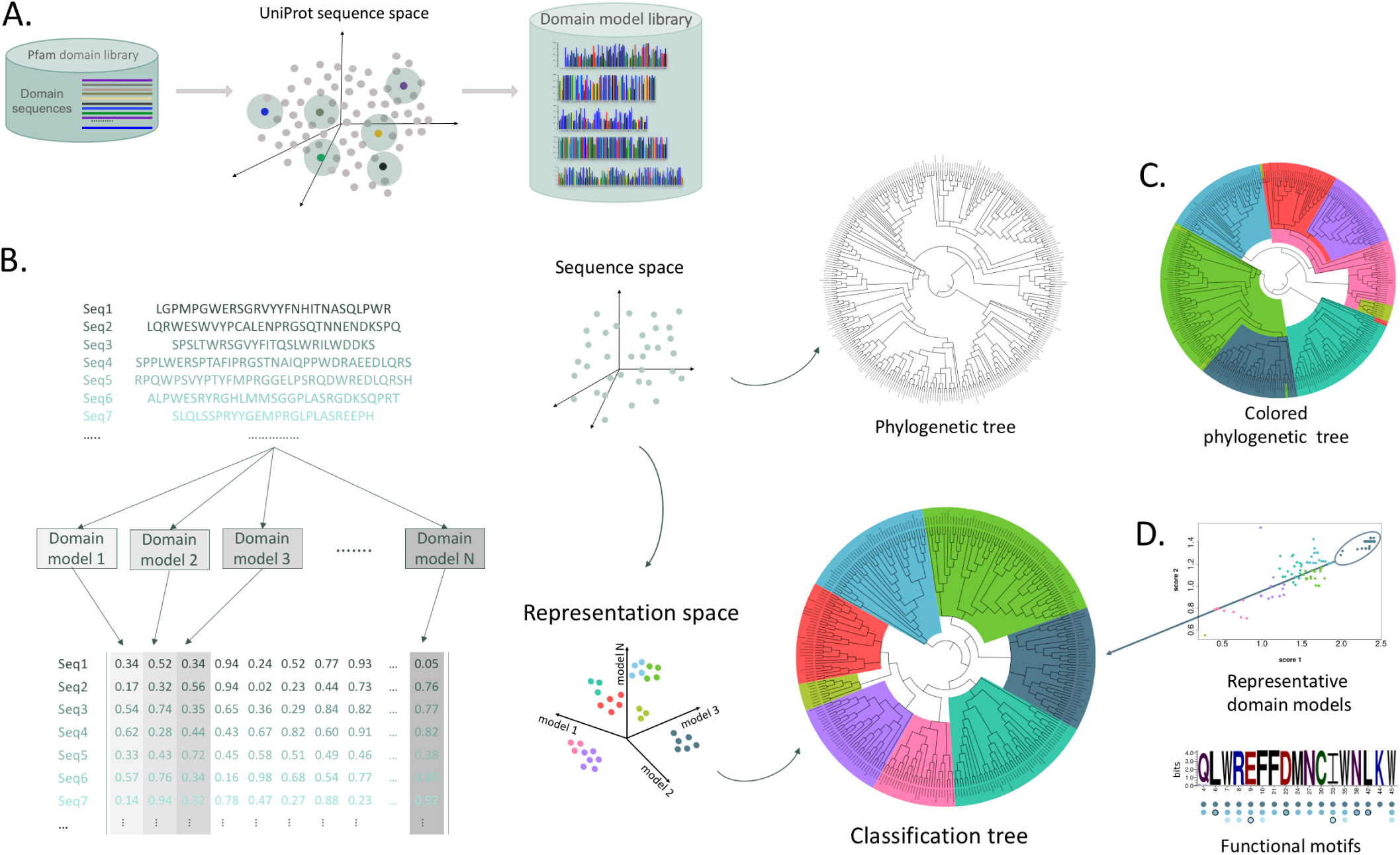
Schema of the ProfileView approach. **A.** Model library construction in ProfileView: all representative sequences from the Pfam domain library are selected for the domain under study. For each representative domain sequence (coloured dots), ProfileView searches for close sequences in UniProt and constructs with HH-Blits several probabilistic models making a library of models for the domain. **B.** Sequences (dots in sequence space, top center) code for proteins with different functions. ProfileView defines a probabilistic mapping from sequences onto the representation space (bottom center) which is indicative of the function of the corresponding protein sequences. The mapping is realised through the contribution of the domain probabilistic models that evaluate the probability of their match against each sequence. Each protein sequence is mapped into a vector of real numbers (coloured row in the matrix, bottom) representing the quality of the match of all models. In sequence space, sequences organise in a phylogenetic tree and, in representation space, they organise in a classification tree based on their distance. ProfileView clusters sequences in the representation space and colors them to indicate a shared function. This coloring is reported in the classification tree. **C.** Phylogenetic tree where sequences are colored as in the classification tree (B). Coloring shows a different organisation within the two trees. **D.** The classification tree allows to identify best representative models for subtrees and their characteristic functional motifs.

The first main idea of ProfileView is to extract conserved patterns from the space of available sequences (**Fig. 1A**; see Methods) through the construction of many probabilistic models for a protein family that should sample the diversity of the available homologous sequences and reflect shared structural and functional characteristics. These models, called Clade-Centered Models or CCM (Bernardes *et al*., 2016; Ugarte *et al*., 2018), are built as conservation profiles. Compared to consensus models (*e.g.*, a pHMM (Eddy, 1998)), they avoid the loss of functional signals when distant sequences are considered. To construct them, we consider the *full* set of sequences *S^i^* associated with a Pfam domain *D^i^* (Finn et al., 2014) and, for each sequence *s_j_* ∈ *S^i^*, we construct a *clade-centered* profile HMM (CCM) by retrieving a set of homologous sequences close to *s_j_* from UniProt (see Methods). Such a model displays features characteristic of *s_j_*and that might differ from other domain sequences *s_k_*∈ *S^i^*. The more *s_j_*and *s_k_*are divergent, the more CCMs are expected to highlight different features. In order to capture feature characteristics of protein interaction sites and/or determinants of functional specificity for protein families likely sharing the same domain architecture, we built highly specific clade-centered models by considering domain sequences in UniProt that display a high sequence identity to *s_j_*. Note that in the past, we constructed CCMs to improve domain annotation (Bernardes *et al*., 2016; Ugarte *et al*., 2018) and, for those models, we employed less restrictive conditions for sequence selection in UniProt.

The second main idea of ProfileView is to use CCMs to embed input sequences into a multidimensional representation space, where each dimension is associated with a CCM (Fig. 1B-D). Namely, for each input sequence to be classified, each model is matched against the sequence, and the value of the match, expressing how close a model is to the sequence, is recorded as a vector entry (**Fig. 1B**, left). This space is called “functional space” because nearby sequences, matching similar profile motifs, are supposed to share the same functional motifs. ProfileView clusters sequences (converted into vectors) within this space by hierarchical clustering and provides a functional classification tree (**Fig. 1B**, bottom right). As illustrated in **Fig. 1C**, the topology of the functional tree is not expected to match the one of the phylogenetic tree. Some of the subtrees of the classification tree will be associated with representative probabilistic models and functional motifs (**Fig. 1D**). Indeed, representative models will be used to subdivide family or subfamily members into smaller groups, in order to capture differences in function-related features of the family, i.e. creating groups that preferably include only one function. All details of the ProfileView pipeline are explained in Method.

### Seven protein families analysed with ProfileView

ProfileView was ran on seven different protein families listed in **Table I** (see **Table S1** and **Table S2** for further characteristics) and was validated on their known functionally characterised sequences. In **Table II**, we provide a quick summary of ProfileView performance by reporting what proportion of sequences is correctly classified by ProfileView for each protein family (see Methods). ProfileView identified a large number of functionally known positions and specific protein residues in interaction with either nucleic acids, amino acids or small molecules. For two families, the Cryptochrome/Photolyase Family (CPF) and the WW domain family, we shall show in detail how ProfileView can provide a functional classification for a large number of functionally uncharacterised sequences, and novel information on conserved amino acids that could be useful to design testing experiments.

**Table I:** Summary of the characteristics of the protein families used for the evaluation: number of sequences, number of sequences after filtering (steps II and III of the pipeline), number of sequences with known function, Pfam domain used for ProfileView classification, MMseq2 clustering condition to identify representative sequences in Pfam (“–” indicates no clustering), number of models constructed for ProfileView analysis. Further features are described in Table S1 and Table S2.

| Superfamily/<br>Family | Subgroup | #seqs | #filt<br>seqs | #func<br>seqs | Pfam domain<br>(accession code) | Clust<br>cond | #models |
| --- | --- | --- | --- | --- | --- | --- | --- |
| Cryptochrome/<br>Photolyase (CPF) | – | 397 | 307 | 72 | FAD (PF03441) | – | 3735 |
| WW domain | – | 349 | 349 | 54 | WW (PF00397) | – | 3733 |
| Glycoside hydrolase<br>family 30 (GH30) | – | 1803 | 1675 | 695 | Glyco-hydro-30 (PF02055)<br>Glyco-hydro-30-2 (PF14587) | – | 1894 |
| Haloacid<br>Dehalogenase | HAD/β-PGM/<br>Phosphatase-like | 391 | 259 | 259 | HAD (PF12710)<br>HAD_2 (PF13419) | $\geq 40\%$ | 4075 |
| Radical SAM | B12-binding<br>domain containing | 273 | 258 | 258 | B12-binding (PF02310)<br>B12-binding_2 (PF02607) | $\geq 60\%$ | 3504 |
| | Methylthiotransferase | 400 | 393 | 393 | Radical.SAM (PF04055) | $\geq 40\%$ | 4501 |
| | SPASM/twitch<br>domain containing | 128 | 29 | 29 | SPASM (PF13186) | $\geq 60\%$ | 2663 |
| | | 128 | 115 | 115 | Radical.SAM (PF04055) | $\geq 40\%$ | 4501 |

**Table II:** Summary of ProfileView performance in classifying functionally characterised sequences. To evaluate what proportion of sequences with a characterised function (TP+FN; see column “# func seqs” in Table I) is correctly classified (TP) by ProfileView, we use the Recall measure (TP/TP+FN; see Methods).

| Protein family | TP | TP+FN | Recall |
| --- | --- | --- | --- |
| Cryptochrome/Photolyase (CPF) | 71 | 72 | 98.6 |
| WW domain | 54 | 54 | 100 |
| Glycoside hydrolase family 30 on EC numbers | 34 | 34 | 100 |
| Glycoside hydrolase family 30 on CAZy families | 694 | 695 | 99.8 |
| HAD/β-PGM/Phosphatase-like | 259 | 259 | 100 |
| B12-binding domain containing | 253 | 258 | 98.1 |
| Methylthiotransferase | 380 | 393 | 96.7 |
| SPASM/twitch domain containing with SPASM domain | 29 | 29 | 100 |
| SPASM/twitch domain containing with Radical SAM | 114 | 115 | 99.1 |

### ProfileView on the CPF family

The Cryptochrome/Photolyase Family (CPF), involved in the interaction with nucleic acids, amino acids and small molecules, is widely distributed in all kingdoms of life (Jaubert *et al*., 2017; Sancar, 2003; Brettel and Byrdin, 2010; Chaves *et al*., 2011). CPF members share the same fold, yet can perform very different functions and have completely different partners: cryptochromes (CRY) are mainly photoreceptors using light to activate specific signalling pathways; some CRY also acts as light-independent transcriptional regulators of the circadian clock; photolyases (PL) are light-activated enzymes repairing UV-damaged DNA (CPD or (6-4) lesions). All CPFs non-covalently bind FAD (Flavin Adenine Dinucleotide) and share a mechanism of FAD photoreduction by intra-protein electron transfer (Björn, 2015). The different CPF functional classes are described in **Supplemental File** (section 1) and listed in the inset legend of **Fig. 2** (bottom right).

**Figure 2:**
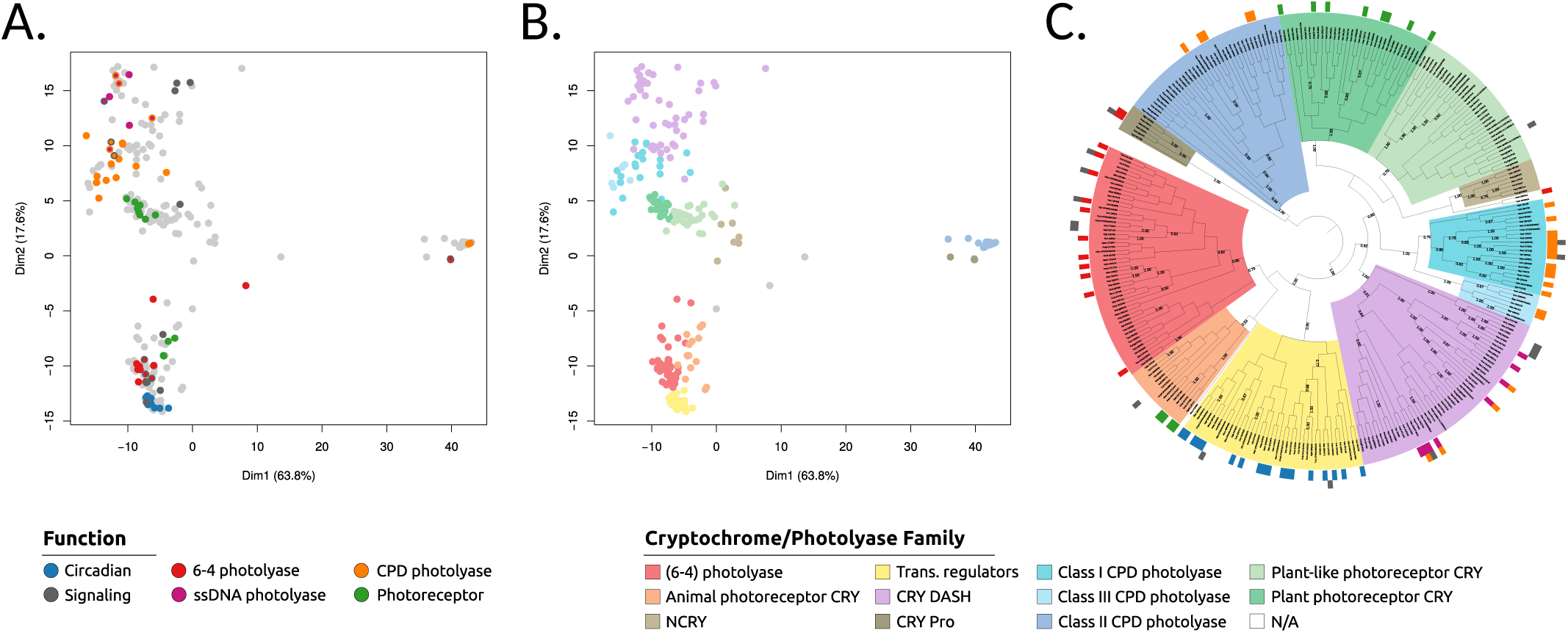
ProfileView representation space and classification tree for the CPF family, and compatibility with experimental work. **A.** 2-dimensional projection of the ProfileView representation space for 307 FAD-binding domain CPF sequences obtained by Principle Component Analysis (PCA). The axes correspond to the first and second PCA components explaining the 63.8% and 17.6% of the dispersion, respectively. Colors correspond to sequences that are either experimentally functionally classified (see the colour legend “Function”) or unclassified (light grey). When a sequence is known to have a double function, it is reported with two colours (the inside colour refers to the known primary function). **B.** As in A, where unclassified points in A are classified by using hierarchical clustering (see the colour legend “Cryptochrome/Photolyase Family”). **C.** The ProfileView classification tree is a finer representation of the hierarchical clustering realised on the high dimensional representation space and illustrated in B. Colors of subtrees are identified by representative models and correspond to known CPF classes (inset legend, right), with the exception of the NCRY subtree. External coloured labels define known functions for the sequences (inset legend, left). Some of the 307 sequences are known to hold multiple functions and are labelled by two colors. The function “signalling” (dark grey) refers to signalling processes of different nature (photoreceptor, transcription, unknown). Numbers on the internal nodes correspond to the percentage of sequences in the corresponding subtree that are separated from the remaining sequences in the tree by the best representative model occurring in the model library (see **Fig. S1** for details).

In our analysis, we make the hypothesis that the FAD (flavin adenine dinucleotide) binding domain, occurring in all CPF sequences, contains all functional information leading to a functional diversification of the family. Indeed, the FAD binding domain is known to non-covalently bind the FAD chromophore which can be in different oxidation and protonation states (Sancar, 2003) specifically associated with different functions. It is also known to interact specifically either with the damaged DNA, with other domains present in CPF proteins (*e.g.*, C-ter extensions in some photoreceptor cryptochromes) or with other protein partners (Czarna *et al*., 2013).

ProfileView is validated on two different types of data: functionally characterised CPF sequences and functionally characterised positions within CPF sequences. These latter are compiled in a manually curated list of positions (**Supplemental File** “CPF mutants used for validation.xlsx”) from the literature. Furthermore, we combined them with structural modelling to analyse CPF subgroups in detail.

#### Validation of ProfileView on the functional diversity of CPF members

The ProfileView representation space shows a consistent functional organisation of CPF sequences (**Fig. 2C** and **Fig. S1**) since sequences known for having the same functional characterisation occur together in large subtrees of the ProfileView classification tree. The perfect split of 71 out of 72 functionally characterised CPF sequences within the 11 subtrees allows us to uniquely associate each subtree with a known functional class (see **Table S3**). This provides the first proof of the method’s classification power.

Most importantly, at the root, the ProfileView tree topology organises large subtrees consistently with known functional classes (**Fig. 3A**). Namely, the ProfileView tree separates light-independent circadian transcriptional regulator CRYs from the light-dependent (6-4) photolyases (PLs) and animal photoreceptor cryptochromes (PR CRY; (**Fig. 3E**, top)). It also clearly separate the DNA repair (6-4) PL from the PR CRY. It reconciles classes I and III cyclobutane pyrimidine dimer (CPD) PLs into a single subtree, while keeping them distinct, and it clearly separates them from plant and plant-like PR CRYs (**Fig. 3F**, top). For the characterised sequences displaying double function (**Fig. S1**), their DNA repair/photolyase activity (either CPD or (6-4)) is consistently determined by ProfileView that groups these sequences in the photolyase subtrees. At the best of our knowledge, these sharp separations, in agreement with known functional characterisations, have never been obtained by sequence analysis before.

**Figure 3:**
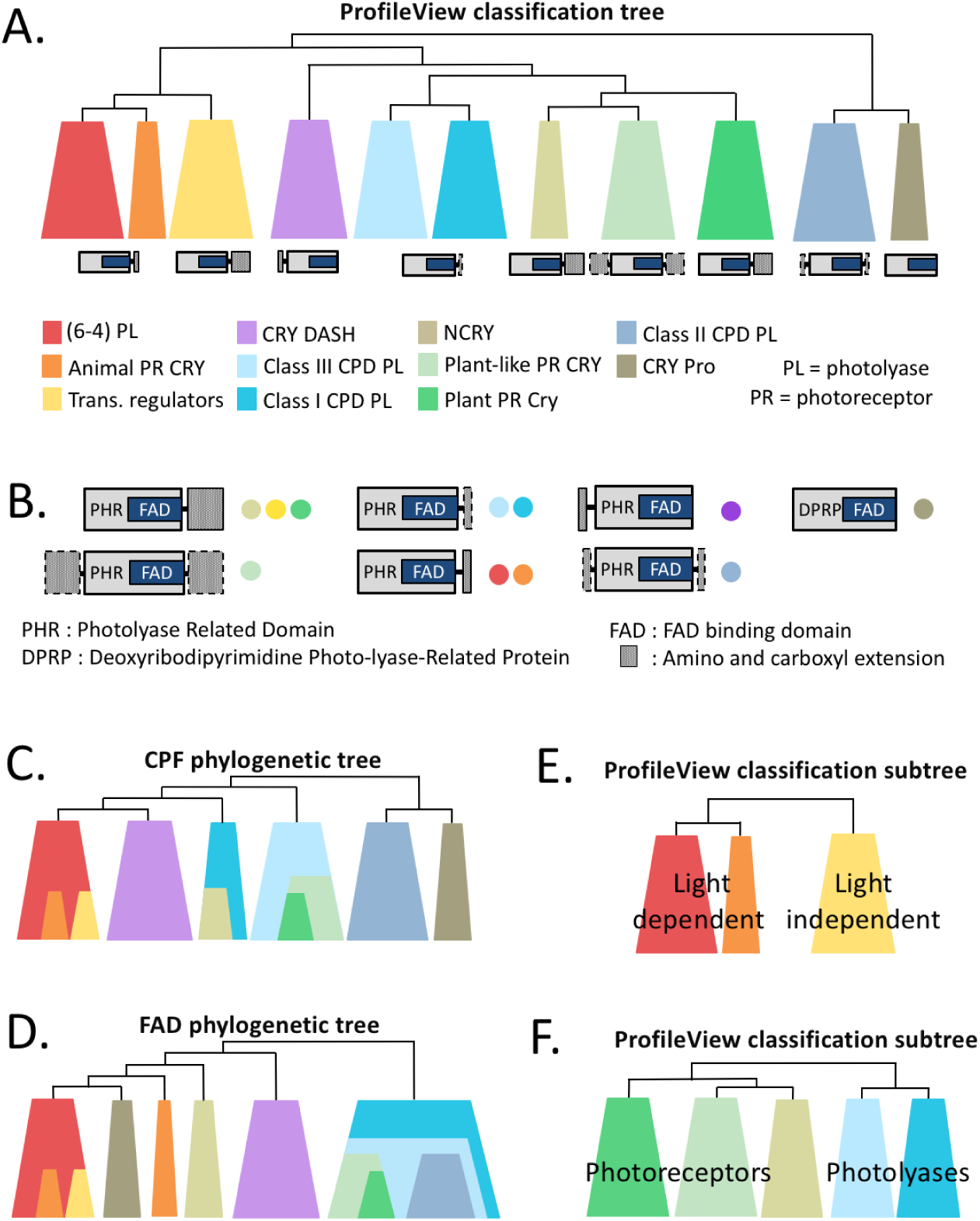
Topological comparison between the ProfileView classification tree and the phylogenetic trees for the CPF family and the FAD binding domain. **A.** Schema illustrating the topological structure of the ProfileView tree in Fig. 2C and **Fig. S1**. Colors correspond to groups of sequences clustering together and comprising sequences with known function (bottom). The domain architectures known to be characteristic of each subtree is reported (see B for more details). **B.** Domain architectures for proteins belonging to different subtrees of A are reported (colours as in A). C- and N-terminal regions are indicated with grey boxes. Dotted border lines indicate terminal regions only present occasionally in an architecture. **C.** Scheme of the main topological structure of the CPF phylogenetic tree constructed from the 307 CPF sequences containing the FAD binding domain. Colors as in A. See the CPF phylogenetic tree in **Fig. S3**. **D.** Scheme of the main topological structure of the FAD phylogenetic tree constructed from the 307 FAD-binding domain sequences. Colors as in A. See the FAD phylogenetic tree in **Fig. S4**. **E, F.** Two zooms on subtrees of the ProfileView classification tree involving classes of CPF sequences described in A. Colors as in A.

Interestingly, the ProfileView tree allowed for the identification of a yet functionally uncharacterized subtree (named NCRY; see **Fig. 2C** and **Fig. 3A**) of proteins showing strong sequence divergence. The same subtree was also identified by sequence similarity network analysis in (Emmerich *et al*., 2020) without inferring any functional classification for it, and by the phylogenetic tree based on the FAD binding domain in CPF sequences (FAD tree, for short; **Fig. 3D**). ProfileView positions NCRY close to the Plant PR CRY and plant-like PR CRY. In contrast, the phylogenetic tree of CPF sequences (CPF tree, for short) includes NCRY within class I CPD PL and the FAD tree places it close to the animal PR CRY and CRY DASH. To our knowledge only one protein from this family has been characterised and it was shown to bind FAD but to lack DNA repair/photolyase activity (Worthington *et al*., 2003) which is in accordance with the position of this family in our functional tree. This finding highlights the potential of ProfileView to reveal novel functional classes within a protein family. (See also **Fig. S2**.)

#### Comparison of the ProfileView tree with the FAD and CPF phylogenetic trees

The comparison of ProfileView classification tree (**Fig. 2C**) with the CPF tree (**Fig. S3**) and the FAD tree (**Fig. S4**) highlights important differences in the topological organisation of major functional classes. A cartoon in **Figs 3ACD** compares the three trees for easy visualization. We notice that the CPF phylogenetic tree (**Fig. 3C**): 1. incorrectly groups sequences exhibiting disparate functions, for instance plant PR CRY and plant-like PR CRY are clustered within class III CPD PL; 2. hides the NCRY subtree within class I CPD PLs; 3. mixes light-dependent and light-independent proteins in a subtree where animal PR CRY and circadian transcriptional regulators are clustered within (6-4) PL sequences. Furthermore, the compatibility of domain architectures associated with different functional classes of CPF sequences (**Fig. 3B**) is coherent with the ProfileView tree topology (**Fig. 3A** bottom) and much less so with the CPF phylogenetic tree. Compare, for instance, the architectures for the classes plant-like PR CRY, plant PR CRY and NCRY, or those for classes I and III CPD PLs. All members of these classes have a PHR domain in which a specific CPF FAD binding domain is found, but C- and N-ter extensions of variable sequence or length. The architectures for plant-like PR CRY, plant PR CRY and NCRY possess N- or C-ter extensions whereas classes I and III CPD PLs only possess the PHR domain. Classes which are topologically close in the ProfileView tree preserve sequence/length characteristics of C- and N-ter regions and agree with what is expected in contrast to the subtrees of the CPF phylogenetic tree.

Similar observations can be highlighted by comparing the ProfileView tree with the FAD phylogenetic tree (**Fig. 3D**).

Summarizing, the reconstruction of ProfileView tree topology highlights three important results: 1. The resolution in two functional groups of light-independent proteins (transcriptional repressor CRY) and proteins which bind the FAD chromophore and need light for their function (PL and PR CRY; **Fig. 3E**); 2. the resolution of classes I and III CPD PL into two distinct sibling subtrees (**Fig. 3F**); 3. the prediction of possible novel functions, by the identification of novel groups as NCRY (**Fig. 3F**).

#### Representative models, motifs and the validation of ProfileView on functionally characterized positions

ProfileView associates representative models and functional motifs to the subtrees of its classification tree. They are used to highlight subfamily delineations and molecular determinants underlying functions and interactions, respectively.

A representative model for a subtree of the ProfileView tree is a probabilistic model that, ideally, “separates” the sequences in a subtree from all other sequences in the ProfileView tree (see step IX of the ProfileView pipeline in Methods). Representative models can be used to subdivide family or subfamily members into smaller groups, in order to capture differences in function-related features at a lower level, i.e. creating groups that preferably include only one function. We remark that all “functional” CPF subtrees corresponding to known subfamilies, highlighted by distinguished colors in **Fig.2C** and **Fig. S1**, are characterised by a representative model which separates at least 50% of the subtree sequences from all other sequences in the ProfileView tree. Moreover, we found representative models associated with several of the internal nodes of the ProfileView tree (**Fig. S1**, where the proportion of sequences supported by a model is indicated on the nodes), and many models separate subtree sequences sharply (100%) indicating functional diversity. An automatic procedure in ProfileView identifies representative models.

Given a representative model for a subtree, the set of conserved positions in the model univocally defines a motif for the corresponding subtree. Motifs associated with the 11 functional subtrees are reported in **Figs. S5, S6** with the exception of classes I and III CPD PL, known to share the same function, that we grouped together by considering the representative model of the minimal subtree including both classes. The only subtree where we found two distinct representative motifs, covering two different regions of the FAD binding domain sequence, is the light-independent transcriptional regulator tree (**Fig. 4AB**). When comparing with the other models, these two models are the only ones which do not cover the FAD binding domain region directly involved in proton or electron transfer to the FAD chromophore, as illustrated in **Fig. 4C** with the alignment of the two transcriptional regulator motifs, the (6-4) PL motif and the animal PR CRY motif. This alignment indirectly shows that proton/electron transfer is not involved in the function of light-independent transcriptional repressors (**Fig. 4C**) despite the importance of the FAD chromophore in their regulation (Hirano *et al*., 2017).

**Figure 4:**
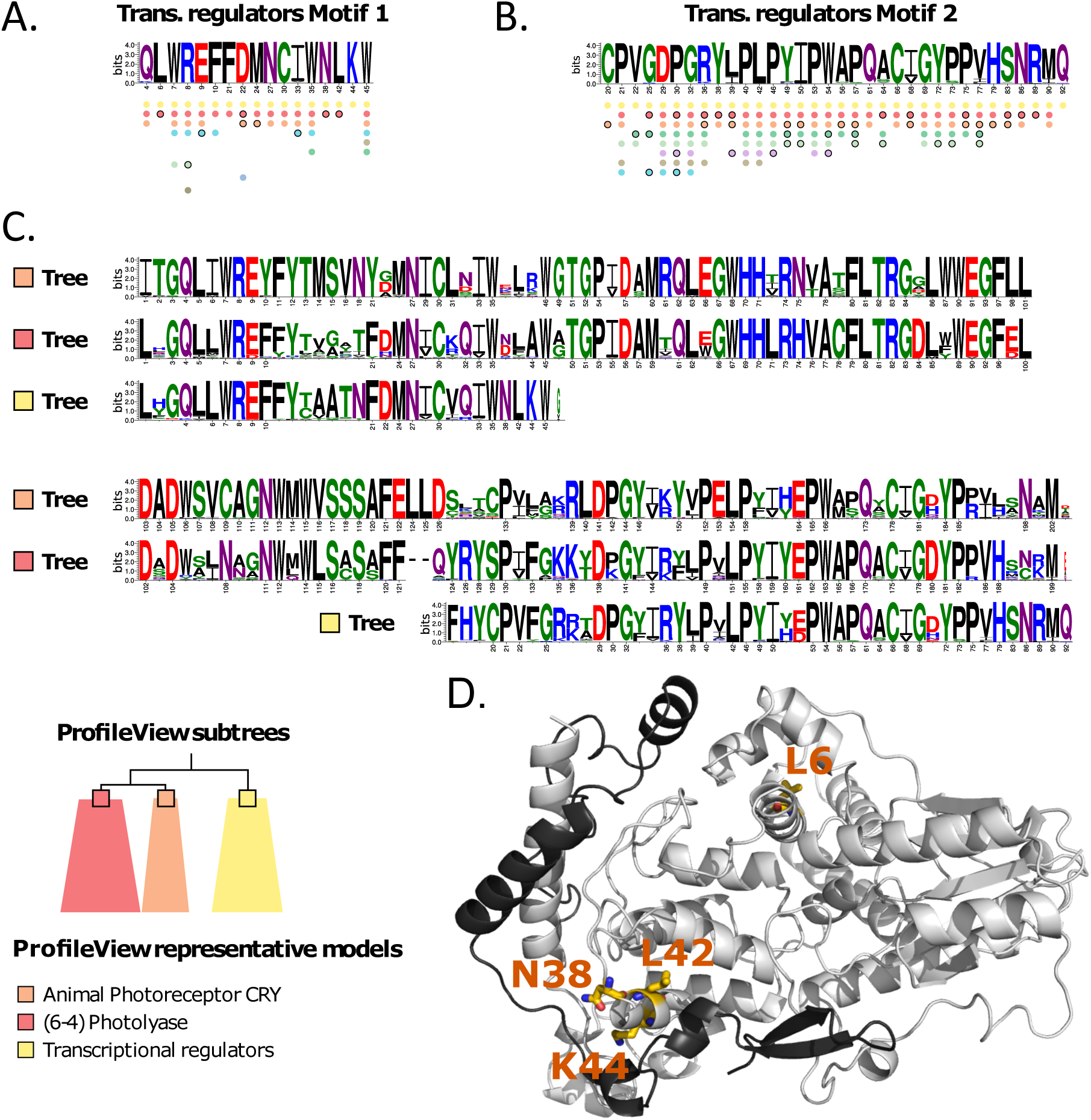
Trans. regulators motifs and their comparison with (6-4) PL and animal PR CRY motifs. **A, B**: two motifs of conserved residues present in light-independent transcriptional regulator sequences. They are extracted from two representative models of the sequences (described in **Fig. S7**) comprising the “yellow” subtree of Fig. 3AE (see also bottom). Numbers (under the letters) correspond to positions in a model, and they are not comparable between motifs. Coloured dots, piled below the motifs, indicate that the corresponding position is well-conserved (see Methods) for the subtrees with the same colour in Fig. 3A. Circled dots indicate positions that are less conserved (see Methods). For each motif, coloured dots are ordered, from top to bottom, depending on the best E-values given by hhblits to the pairwise alignments. **C**. Three representative motifs associated with the trans. regulators (yellow), (6-4) PL (red) and animal PR CRY (orange) subtrees of the ProfileView tree are aligned. Numbered positions correspond to conserved positions belonging to the associated representative motif. The absence of the number indicates less conserved positions. The alignment has been constructed using trans. regulators motifs as template models and all others as query models. The length of a motif depends on the length of the associated model, selected as best representing the sequences in a subtree. **D**. PDB structure (4CT0) of the interacting mouse cryptochrome mCRY1 (grey) and Period2 mPER2 (black) involved in the circadian clock. The four residues highlighted in the structure, N38, L42, K44 and L6, have been explained in the text.

To validate ProfileView motifs, we exploited the functional information derived by characterized mutations and looked whether their conserved amino acid positions would identify known functional natural variations, single amino acid residue replacements by site-directed mutagenesis or random mutagenesis, and structural specificity when structures were available. For this, we manually curated a list of experimentally characterized positions in CPF sequences (see **Supplemental File** “CPF mutants used for validation.xlsx”). Most of these positions display mutations causing loss of function or phenotypic changes. They are often involved in binding with other proteins, DNA substrates or with the cofactor FAD; active amino acids involved in catalytic or allosteric sites, such as DNA repair for PLs or post-translational modifications in CRY, are also identified. **Table S3** summarizes how many ProfileView positions are validated by current experimental evidence. Interestingly, it finds a number of highly specific positions for CPF functional classes that have not been reported in the literature before. We discussed these positions together with other observations in **Supplementary File**. They illustrate the great deal of functional information that can be extracted from representative motifs and be used to design tailored experiments for discovering new functional activities or novel biological mechanisms involving the FAD binding domain.

### How evolutionary close sequences are distinguished by motifs in ProfileView classification space?

ProfileView can distinguish very similar sequences associated to different functions. We illustrate this crucial feature with a concrete example, based on representation models and motifs. We consider the pair of sequences U5NDX3 and R7UL99, belonging to the CPF family. They are grouped together by phylogenetic analysis because very similar (sequence identity is 61.8% and sequence similarity is 74.7%) and are classified in different functional groups by ProfileView, as a photolyase and a transcriptional regulator respectively. The conserved positions belonging to the photolyase functional motif (motif called “(6-4) photolyase” in **Fig S5**) constructed from ProfileView analysis are shown in the alignment reported in **Fig. S8**. For almost all positions in the motif, the corresponding amino acid is conserved in both sequences (in green) as expected by the high sequence identity of the alignment. For positions 1, 33 and 135 in the motif, the amino acid is conserved only in the U5NDX3 sequence. This means that the photolyase representative model will provide higher matching values for U5NDX3 than for R7UL99. Moreover, two of these positions, 1 and 135, are highly conserved in the photolyase family and variable in the transcriptional regulator family (see dots below the motif in **Fig S5**) making the U5NDX3 sequence closer in classification space to the photolyase subgroup than R7UL99. Note that these observations concern the dimension of ProfileView classification space which is associated with the “(6-4) photolyase” model, but that is the contribution of all probabilistic models, one for each dimension of the classification space, that will define the position of the sequences bringing them closer either to the photolyase subgroup or the transcriptional regulator subgroup, in this specific example. A second example is reported in **Fig. S9** for sequences Q6MDF3, D8UF46 and Q485Z2, where the phylogenetic tree could wrongly suggest an ancestral function, conserved in paraphyletic groups separated by clades where neofunctionalization would occur. The analysis of the sequence alignment (see legend in **Fig. S9**) highlights those positions explaining functional classification.

### ProfileView on the WW domains

The WW domain family is found in many eukaryotes. WW domains are protein modules mediating protein-protein interactions through recognition of proline-rich peptide motifs and phosphorylated serine/threonine-proline sites. They are involved in a number of different cellular functions (Ingham *et al*., 2005) such as transcription, RNA processing, receptor signalling and protein trafficking, and in several human diseases such as muscular dystrophy, cancer, hypertension, Alzheimer’s, and Huntington’s diseases. Their functional classification is far from being straightforward because based on the sequence motif and the binding affinity of the peptides targeted by WW domains. In particular, the same WW domain can bind with variable affinity to multiple peptides (Sudol and Hunter, 2000; Otte *et al*., 2003; Russ *et al*., 2005), and it is the modulation of binding properties that make hundreds of WW domains to interact specifically with hundreds of putative ligands in mammalian proteomes (Sudol and Hunter, 2000). WW domains have been experimentally classified in six interaction groups by Otte *et al* (Otte *et al*., 2003) (Y, R*_a_*, R*_b_*, L, poly-P, poS/poT), in 3 groups by Ingham *et al* (Ingham *et al*., 2005) (A, B and C) and in 6 groups by Russ *et al* (Russ *et al*., 2005) (I, Im, I/IV, II, III, IV). These three functional classifications were based on target peptide sequence motifs and their binding affinity. (**Fig. S10** shows the localisation of known classified sequences on the phylogenetic tree for WW domains.)

#### Validation of ProfileView on the functional diversity of WW domains

All natural sequences (60) analysed in (Otte *et al*., 2003; Ingham *et al*., 2005; Russ *et al*., 2005) and upgraded with a set of 289 natural sequences randomly selected from (Tubiana *et al*., 2019) and not yet experimentally characterized, have been considered for classification by ProfileView. Of the 60 experimentally tested natural sequences, 54 of them have been experimentally functionally characterized (**Table I**).

ProfileView tree is organised in seven subtrees as illustrated in **Fig. 5** and **Fig. S11**. The three independent experimental characterisations classify the 54 sequences in groups/classes that turn out to belong to specific ProfileView subtrees as shown in **Table S4**, and in **Fig. 5** and **Fig. S11** by the three first internal layers of colored squares (corresponding to the 54 characterised sequences). In other words, ProfileView perfectly classifies the 54 known functionally characterised sequences: the 35 sequences in Ingham’s group A, Russ’s class I and Otte’s Y-group are grouped together in one large ProfileView subtree (*T*_1_ in **Table S4** and **Fig. S11**); the 2 sequences in Ingham’s group C, Russ’s class IV and Otte’s posS/posT-group are grouped in two main ProfileView subtrees (*T*_6_&*T*_7_ in **Table S4** and **Fig. S11**); the remaining 17 sequences are organised in 4 other ProfileView subtrees corresponding to 4 different Otte’s groups (*T*_2_ to the L-group, *T*_3_ to the Ra-group, *T*_4_ to the Rb-group and *T*_5_ to the posS/posT-group; see **Table S4** and **Fig. S11**).

**Figure 5:**
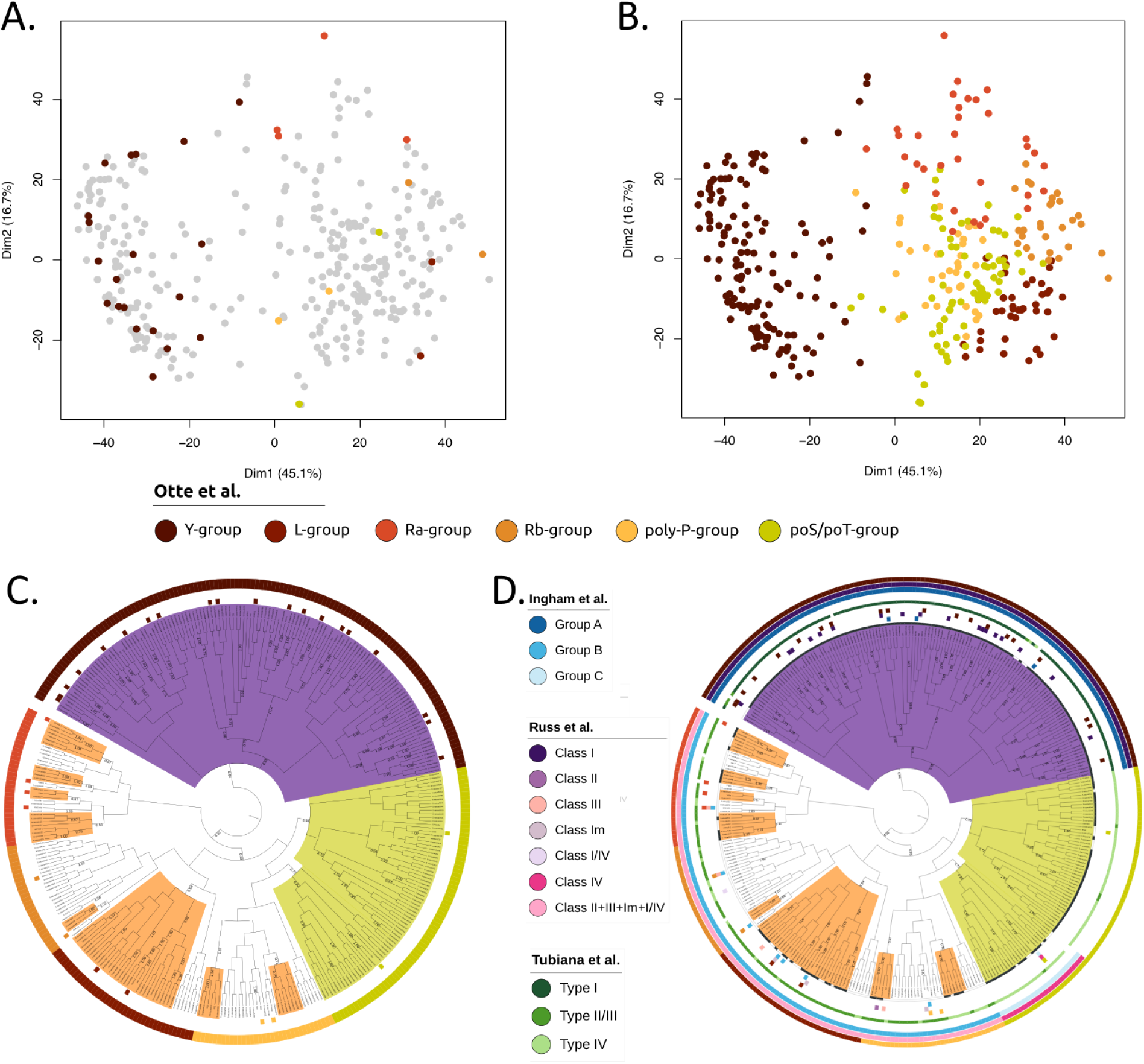
ProfileView representation space and ProfileView tree for WW domains; compatibility with experimental and computational classification. **A.** 2-dimensional projection of the ProfileView representation space for the WW domain natural sequences studied in (Otte *et al*., 2003; Ingham *et al*., 2005; Russ *et al*., 2005; Tubiana *et al*., 2019) obtained by PCA. The first and second PCA components explain 45.1% and 16.7% of the dispersion, respectively. Colors follow Otte’s functional classification of 32 WW sequences (Otte *et al*., 2003); sequences not classified in (Otte *et al*., 2003) are in light grey. **B.** As in A, where light grey sequences are classified by ProfileView in subsets associated with Otte’s functional classes (Otte *et al*., 2003) by hierarchical clustering (see main text). **C.** The ProfileView tree represents the hierarchical clustering, realised on the high dimensional ProfileView space; see B. All subtrees with a representative model are indicated by a root labeled by the percentage of sequences in the subtree best matching the model (see Methods). Among these subtrees, those containing at least 3 sequences are coloured. 54 coloured squares are reported as functionally characterised, and 6 more were left unclassified in (Otte *et al*., 2003; Ingham *et al*., 2005; Russ *et al*., 2005) (KIAA1052, PRP40-1, CA150-1, CA150-3, HYP109-2, IQGAP1). The 6 WW unclassified sequences are classified by us in different subtrees corresponding to different Otte’s groups: PRP40-1, CA150-1 in L-group; HYP109-2, IQGAP1 in Ra-group; CA150-3 in Rb-group; KIAA1052 in poS/poT-group. **D.** Same tree as in C, where sequences in a coloured subtree that are best matched by the representative model are highlighted in black in the first circular stripe surrounding the tree. As in C, the three experimental characterisations of natural sequences are reported in three layers made of small squares around the tree: “Otte” (Otte *et al*., 2003) (brown scale; as in C), “Ingham” (Ingham *et al*., 2005) (blue scale) and “Russ” (Russ *et al*., 2005) (purple scale). “Tubiana” computational classification (Tubiana *et al*., 2019) is reported in green scale; WW sequences not considered in Tubiana are left white. The three most external circular stripes show the compatibility between the grouping suggested by Otte, Russ and Ingham with ProfileView subtrees as described in **Table I**. The larger subtree comprising a given function, in the sense of either Otte, Russ and Ingham, gives the colour of the function to the corresponding portion of the stripe. Note that one of our trees is not functionally annotated by experimental data coming from Ingham nor Russ. (See **Fig. S11** and **Table I**.)

When comparing ProfileView classification to Russ’ study, it is interesting to consider the nature of this latter. Indeed, Russ *et al* look at combinations of binding specificities of WW domain sequences to different peptides and classifies them in 6 classes accordingly. Sequences in classes Im, I/IV, II and III display complex binding patterns involving different proline-rich peptide motifs whose differentiation is not obvious. Accordingly, these sequences belong to various ProfileView subtrees (*e.g.* class III sequences are spread on subtrees *T*_2_-*T*_5_), different from those involving class I and class IV. Therefore, in **Fig. 5** and **Fig. S11** (see pink color in the outer circle) and in **Table S4**, we grouped classes Im, I/IV, II and III together.

As shown in **Fig. 5**, not all the seven ProfileView subtrees are associated with a unique representative model (see legend). Multiple models might be associated to subsubtrees and describe different groups of sequences within the class, suggesting a finer functional organisation for this subfamily of WW domains. All identified representative models are reported in **Fig. S12** and their associated motifs in **Figs. S13**, **S14**.

### ProfileView on the GH30 family of the CAZy database

The glycoside hydrosylases (EC 3.2.1.-), in short GH, are a widespread group of enzymes which hydrolyse the glycosidic bond between two or more carbohydrates or between a carbohydrate and a non-carbohydrate moiety. Their classification, based on substrate specificity and occasionally on molecular mechanisms, turned out to be particularly difficult. For this, a vast knowledge about these enzymes has been meticulously curated in the CAZy database (Lombard *et al*., 2014). The GH30 is one of the GH families that has been organised in subfamilies in CAZy (http://www.cazy.org/GH30.html). It counts nine different subfamilies (GH30-1,…, GH30-9) corresponding to eleven different enzymatic chemical reactions. Some of these subfamilies are functionally classified by CAZy and some others are left unclassified.

#### Validation of ProfileView on the functional diversity of GH30 sequences

We considered the set of GH30 sequences and their classification in CAZy. ProfileView representation space and ProfileView tree for these sequences have been constructed using models coming from two similar PFAM domains, PF02055 (Glyco hydro 30) and PF14587 (Glyco hydr 30 2). The topology of the ProfileView tree (**Fig. S15** and **Fig. S23B**) perfectly separates sequences in the nine CAZy subfamilies GH30-1,…, GH30-9 into subtrees (only one GH30-3 sequence is placed within GH30-2 sequences). Furthermore, the subtrees well separate the EC numbers in CAZy functional annotation (see **Fig. S15**, **Table S5** and **Table S6**).

Four subfamilies (GH30-1, GH30-2, GH30-7 and GH30-9) are characterised by representative models directly explaining the separation of their sequences from all other GH30 sequences in the tree (**Fig. S23B**). In **Fig. S15**, the grey dots indicate the existence of representative models for many ProfileView subtrees, highlighting a possible functional sub-characterization for several CAZy subfamilies. For instance, note that the two CAZy reactions 3.2.1.45 and 3.2.1.21+3.2.1.37 for GH30-1 are identified in distinguished subtrees (green and violet labels are associated to reactions 3.2.1.45 and 3.2.1.21+3.2.1.37 in **Fig. S15**) separated by a representative model. Furthermore, for the GH30-3 subfamily, several sequences labelled by CAZy reaction 3.2.1.75 occur in different GH30-3 subtrees characterized by representative models, highlighting potential functional differences within this subfamily.

### ProfileView on the enzyme superfamilies of the Structure-Function Linkage Database

The Structure-Function Linkage Database (SFLD) is a manually curated classification resource describing structure-function relationships for functionally diverse enzyme superfamilies (Schnoes *et al*., 2009; Akiva *et al*., 2014). Despite their different functions, members of these superfamilies “look alike” making them easy to misannotate. We challenge ProfileView against these sets of sequences and show that its classification meets the functional information in SFLD.

SFLD is organised in superfamilies whose members are subdivided into subgroups using sequence information, and lastly into families, that is sets of enzymes known to catalyze the same reaction using the same mechanistic strategy. Subgroups are not organised by function, and the functional specificity of the sequences is detailed at the family level. We consider two different superfamilies, Haloacid Dehydrogenase and Radical SAM, because of their large variety of functions. Indeed, the Haloacid Dehydrogenase family is characterized by 25 subgroups organized in 22 families and 20 different reactions, and the Radical SAM family by 58 subgroups organised in 98 families and 85 reactions (see sfld.rbvi.ucsf.edu/archive/django/ superfamily/index.html for a detailed description). We analysed the HAD/*β*-PGM/Phosphatase-like sub-group of Haloacid Dehydrogenase and three subgroups of Radical SAM: B12-binding domain containing, Methylthiotransferase and SPASM/twitch domain containing. ProfileView functional classification has been validated on the SFLD families associated with the four subgroups.

#### ProfileView on the HAD/*β*-PGM/Phosphatase-like subgroup

Characterized functions included in this subgroup include 2-haloacid dehalogenase, beta-phosphoglucomutase, phosphonoacetaldehyde hydrolase, and phosphatases of various specificities (see sfld.rbvi.ucsf.edu/archive/django/subgroup/ 1129/index.html). We run ProfileView on a model library constructed from the two similar Pfam domains HAD and HAD 2 (see Table I). ProfileView groups all known sequences belonging to known characterized functions correctly, in separated subtrees, as illustrated in **Fig. S16** and **Table S7**. Moreover, for each subtree, it provides a model separating the set of sequences in the subtree from the rest of the set. The exception relies on one family, the 2-deoxyglucose-6-phosphatase which is grouped with the glycerol-3-phosphate phosphatase, represented by only two sequences correctly grouped together, and for which a model separates both functions from the rest of the tree.

#### ProfileView on the B12-binding domain containing subgroup

All the enzymes in this subgroup appear to have a Vitamin B12 Binding domain and are involved in many different reactions (see sfld. rbvi.ucsf.edu/archive/django/subgroup/1082/index.html). We run ProfileView on a model library constructed from the two similar Pfam domains B12-binding and B12-binding 2 (see **Table I**). **Fig. S17** and **Table S8** describe ProfileView classification in three large subtrees associated with three families, which are represented by tens of sequences. The remaining five families are underrepresented, four comprise exactly one sequence and the fifth one only three sequences (grouped together by ProfileView, see “paromamine deoxygenase” in **Fig. S17**). Underrepresented families are localised within two large subtrees of the ProfileView classification tree, the bacteriocin maturation and the hopanetetrol cyclitol ether synthase (see **Fig. S17**). Consistently, note that ProfileView does not propose a model separating these two large subtrees but it proposes one separating the anaerobic magnesium-ptotoporphyrin-IX monomethyl ester cyclase family from the rest.

#### ProfileView on the Methylthiotransferase subgroup (MTTase)

All enzymes of this subgroup are organised around 4 families (see sfld.rbvi.ucsf.edu/archive/django/subgroup/1061/index.html) that have been defined in SFLD by considering the domain architecture of MTTase sequences comprising an N-terminal MTTase domain, a central radical generating fold domain and the C-terminal TRAM domain, not shared by other Radical SAM outside the MMTase. In contrast, to classify MTTase sequences, we used the Radical SAM domain only, shared by all subgroups of the superfamily. This domain allowed ProfileView to split the sequences in 4 main subtrees corresponding to the four known families as reported in **Fig. S18** and **Table S9**. A few sequences are misplaced compared to SFLD classification. ProfileView proposes many models splitting the four families. In particular, it proposes two representative models splitting the MiaB-like and CDK5RAP1 families from the rest and viceversa (**Fig. S18**).

#### ProfileView on the SPASM/twitch domain containing subgroup

We used ProfileView to study this subgroup (see sfld.rbvi.ucsf.edu/archive/django/subgroup/1067/index.html) through two independent analysis, one based on the Radical SAM domain and other on the SPASM domain. This is an intrinsically difficult set, not only for functional annotation but also for domain annotation. Indeed, we identified the SPASM domain in 29 sequences based on the ProfileView model library of 4501 SPASM models, while only 6 of these sequences have been annotated by Pfam with a SPASM domain. ProfileView classification of the SPASM/twitch domain containing sequences based on SPASM domain organises the seven known SFLD functional families in distinct subtrees (see **Fig. S19** and **Table S10**). **Fig. S20** and **Table S11** describe ProfileView classification based on Radical SAM domain. Families are well organised in distinguished subtrees supported by representative models.

### Comparison of ProfileView with other computational approaches

ProfileView is compared with the PANTHER classification system (Mi *et al*., 2012, 2013), the state-of-the-art neural network approach based on Restricted Boltzman Machines (RBM) in (Tubiana *et al*., 2019), and the CUPP platform (Barrett and Lange, 2019). In all comparisons it proves to overcome or be on par with the functional classification considered.

#### ProfileView and PANTHER

PANTHER (Mi *et al*., 2012, 2013) is a large curated biological database of gene/protein families and their functionally related subfamilies which has been designed to classify and identify the function of gene products. PANTHER provides data and tools to group sequences in functional clusters. Contrary to ProfileView, it does not organise them in a distance tree, missing the possibility to identify large-scale functional properties for groups of sequences clustering together, like the light dependent/independent CPF sequences. Comparison was realised on the full CPF family. For easier visualization, we reported PANTHER classification on both the ProfileView classification tree and the CPF distance tree in **Figs S21** and **S22**. ProfileView and PANTHER agree on several functional classes: “SLR1343 PROTEIN” for PANTHER and CRY Pro for ProfileView; “ZGC:66475” PANTHER and Class II CPD PL for ProfileView. Other PANTHER groups are function specific but they do not recognise the full functional subgroup, such as “CRYPTOCHROME 1A”, “CRYPTOCHROME 2B-APOPROTEIN” and “CRY2AProtein” for PANTHER that characterize a part of the Plant Photoreceptor CRY sequences. Finally, other PANTHER groups collect functions from different functional classes: “(6-4) PHOTOLYASE ISOFORM A” recognizes (6-4) PL, Class I CPD PL and NCRY; “CRYPTOCHROME-1” recognizes Plant PR CRY and circadian rhythms transcriptional regulators, which are light independent; “SI:CH1073-390K14.1” recognises both PR and PL. In particular, experimentally characterized CPF sequences show PANTHER limitations: many known (6-4) PL (red subtree) are annotated as circadian regulators, Class I CPD PL is partly classified as PR instead. Note also that no distinction between Class I, II, III CPD PL is evident in PANTHER classification, and that sequences in the NCRY subtree are annotated as (6-4) PL while they are PRs according to us and to (Emmerich *et al*., 2020).

#### ProfileView and the RBM approach

The state-of-the-art neural network approach based on Restricted Boltzman Machines (RBM) in (Tubiana *et al*., 2019) relies on the generative modelling of correlations in sequence alignments, it extracts biologically interpretable features but demands a particularly heavy training and computational time. The RBM approach classifies the dataset of 349 WW domain sequences in 3 groups (I, II/III, IV). Associated protein binding motifs have been proposed (Tubiana *et al*., 2019).

The RBM approach correctly classifies 49 out of 52 functionally characterised sequences considered in (Tubiana *et al*., 2019) as described in **Table S4**. It misclassified 3 sequences, compared to ProfileView correct classification of 54 out of 54 sequences. Note that the large group II/III, indistinguishable in Tubiana *et al*, is organised into several ProfileView subtrees of sequences known to bind to specific peptides, as shown in (Otte *et al*., 2003), providing a refined analysis of binding motifs. ProfileView tree also classifies, within its subtrees, many experimentally uncharacterized WW domain sequences, largely agreeing with Tubiana’s classification but not always, as seen in **Fig. 5** and **Fig. S11**.

In conclusion, compared to RBM (Tubiana *et al*., 2019), ProfileView is more precise, it extracts from the analysis molecular determinants underlying protein functional diversity and it is much faster. Its computation time is measured in hours versus days for RBM (see Methods and **Table S2**).

#### ProfileView and CUPP

CUPP (Barrett and Lange, 2019) is a computational approach designed to classify by using short peptide sequences expected to be specific for functional characterization of carbohydrate-active enzymes. In CUPP, proteins sharing the same peptide profile are claimed to share the same function.

The set of GH30 sequences used to validate ProfileView was also used for the evaluation of CUPP (Barrett and Lange, 2019). CUPP split these sequences in 33 groups and organised them in a dendogram (Barrett and Lange, 2019) whose topology is reported in **Fig. S23A**. The dendogram is composed of 9 subtrees corresponding to the 9 CAZy subfamilies. A schematic comparison of CUPP dendrogram (Barrett and Lange, 2019) and ProfileView tree is given in **Fig. S23**. Both their topologies highlight the separation of the CAZy subfamilies GH30-1, GH30-2, GH30-3 and GH30-9 from the other subfamilies. ProfileView tree separates further subfamilies GH30-4 and GH30-5 from the remaining ones.

A detailed analysis of the CAZy subfamilies indicates similar sequence organisation for the two methods. For instance, CUPP organises GH30-1 sequences by splitting them in five clusters (Barrett and Lange, 2019) that are easily identified in ProfileView tree, where three representative models are associated to three of CUPP clusters (purple, fuchsia and dark blue in third circle of annotation in **Fig. S15**). In contrast, the classification of CAZy subfamilies GH30-4 and GH30-5 (**Fig. S15**) highlights a large number of CUPP clusters while ProfileView groups GH30-5 into three main subtrees and GH30-4 into one. Two of the three ProfileView subtrees grouping GH30-5 are characterized by representative models. Interestingly, the remaining sequences are clusterized by CUPP into several clusters and no representative model is found by ProfileView, indicating the difficulty of both methods to classify this group of sequences.

To test the general applicability of ProfileView versus CUPP, which was designed for enzyme proteins, we also compared the two approaches on the CPF sequences and on the WW domain sequences. This analysis highlights CUPP’s limitations in handling arbitrary protein families.

On the WW domain sequences, CUPP does not provide any insightful classification, as **Fig. S27** shows. This is probably due to the very short length of this domain, between 35 and 40aa long.

On the CPF family, CUPP was run using both FAD and PHR sequences. CUPP tree and its associated clusters are represented in **Fig. S24** for FAD sequences (see also **Fig. S25**). CUPP: 1. groups all together the CPF classes “Transcriptional regulators”, (6-4) PL and Animal PR CRY. Hence, distinguished functions are shared in the same subtree. In particular, it does not distinguish light dependent from light independent protein sequences; 2. does not distinguish Class I and III CPD PL; 3. places the CRYPro subtree far from the remaining subtrees while, in ProfileView, CRYPro is located closer to Class II CPD PL; 4. splits the CRY DASH tree into two subtrees. There is no known functional annotation for one of the subtree and, therefore, it is not clear whether it is a relevant sequence split or not. ProfileView organises sequences in this subtree differently.

Furthermore, CUPP succeeds to classify a larger number of sequences (corresponding to the leaves left uncoloured in **Fig. S24**) in the CPF family compared to ProfileView that did not find, among its models, sufficient confidence to include some input sequences in its tree. Viceversa, there are sequences that have been classified by ProfileView and that do not belong to CUPP classification (see uncolored sequences within CUPP clusters in **Fig. S24**). We also notice that, as ProfileView, CUPP: 1. groups Class II CPD PL in a single subtree, and 2. distinguishes NCRY sequences.

When CUPP considers the whole PHR sequence, the topology of the CUPP tree (**Fig. S26B**) gets closer to ProfileView topology even though CUPP keeps mixing Class I and III CPD PLs as well as light dependent (6-4) PL and Animal PR CRY sequences; the NCRY subtree locates close to photolyases (**Fig. S25**); the higher number of CUPP clusters fragments the functional organisation, as for instance for Class II CPD PL.

## Materials and Methods

### Datasets used to validate the method

The seven protein families used to evaluate ProfileView performance are listed in Table I (first and second column). Their sets of homologous sequences have been retrieved from publicly available databases (see below; see Table I, third column, for their number). All families present multiple functions. For each family, a subset of sequences has been functionally classified before (see Table I, fifth column) and it has been used for evaluation. Various protein sequence characteristics are reported in **Table I**, **Table S1 and Table S2**. CPF sequences were retrieved from UniProt, JGI projects (genome.jgi.doe.gov), and OIST projects (marinegenomics.oist.jp). The set was constructed following two main criteria: 1. it contains CPF sequences known to have a specific function according to experimental evidence reported in the literature (see **Supplemental File** for bibliographical references); 2. it contains CPF sequences that span the whole tree of life; they belong to 146 species, 74 classes, and 40 phyla (see **Supplemental File** for the detailed list). In the text, a “CPF sequence” refers to the full length CPF sequence comprising the PHR domain, including the FAD binding domain, and possibly the C- and N-terminal extensions, while a “FAD sequence” refers to the FAD binding domain sequence exclusively.

The set of WW domain sequences was constructed by combining the datasets of natural sequences analysed in (Otte *et al*., 2003; Ingham *et al*., 2005; Russ *et al*., 2005; Tubiana *et al*., 2019). 60 sequences have been experimentally characterized (Otte *et al*., 2003; Ingham *et al*., 2005; Russ *et al*., 2005) and the remaining ones have been randomly selected, in comparable proportion, from the three sets classified in Tubiana *et al* (types I, II/III, IV) (Tubiana *et al*., 2019).

The set of GH30 sequences is the same used in (Barrett and Lange, 2019) (file GH30.faa provided with the CUPP program v1.0.14 and containing 1803 sequences) and described in the Carbohydrate-Active Enzymes database CAZy (http://www.cazy.org/GH30.html). It is organised in different subfamilies of the CAZy classification. Some of these subfamilies are functionally classified by CAZy and some others are left unclassified. We used the annotation files in (Barrett and Lange, 2019), where 721 of the 1803 sequences have a mapping/label to the subfamilies from GH30-1 to GH30-9. Note that, of the 1675 sequences retained for analysis by ProfileView after filtering, 695 have a label in the GH30 ProfileView tree (**Table I**).

The set of sequences of the HAD/*β*-PGM/Phosphatase-like subgroup of the Haloacid Dehalogenase (HAD) superfamily and of the three subgroups of the Radical-SAM superfamily (B12-binding domain containing, Methylthiotransferase and SPASM/twitch domain containing) have been retrieved from the Structure-Function Linkage Database (SFLD) (Schnoes *et al*., 2009; Akiva *et al*., 2014). Namely, each sub-group is defined by the union of the sets of annotated sequences associated with its families in SFLD. Given a subgroup, we considered all its families, even if they were represented by very few sequences, possibly only one.

### Clade-Centered Models and a multi-source functional annotation

Widely used search methods (Altschul *et al*., 1997; Eddy, 2011; Remmert *et al*., 2011) are based on a mono-source annotation strategy, where a single probabilistic model (*e.g.*, a pHMM (Eddy, 1998)), generated from the consensus of a set of homologous sequences, is used to represent a protein domain. The mono-source strategy usually performs well for rather conserved homologous sequences, but when sequences have highly diverged, consensus signals become too weak to generate a useful probabilistic representation and global-consensus models do not characterize domain features properly. A *multi-source* domain annotation strategy (Bernardes *et al*., 2016), in which protein domains are represented by several probabilistic models, called *Clade-Centered Models* (CCM), was implemented in CLADE (Bernardes *et al*., 2016) and MetaCLADE (Ugarte *et al*., 2018) for genomes and metagenomes/metatranscriptomes respectively.

To construct CCMs (see below), we consider the *full* set of sequences *S^i^* associated with a Pfam domain *D^i^* (Finn et al., 2014) and, for each sequence *s_j_*∈ *S^i^*, we construct a *clade-centered* profile HMM (CCM) by retrieving a set of homologous sequences close to *s_j_*from UniProt. Such a model displays features characteristic of *s_j_*and that might differ from other domain sequences *s_k_*∈ *S^i^*. The rationale is that the more *s_j_*and *s_k_*are divergent, the more clade-centered models are expected to highlight different features. It has been shown that CCMs significantly improve domain annotation (both for full genomes (Bernardes *et al*., 2016) and for metagenomic/metatranscriptomic sequences (Ugarte *et al*., 2018)) and, due to their closeness to actual protein sequences, they are more specific and functionally predictive than the canonical global-consensus approach. In this work, however, we build and use CCMs differently aiming at better resolve the functional organisation of sequences within protein families, whose sequences likely share the same domain architecture. In order to capture conserved motifs likely to be of functional relevance, we built highly specific clade-centered models. They will likely belong to protein interaction sites, be made of conserved positions on subsets of homologs, and be determinants of functional specificity.

### The ProfileView method

A flowchart describing ProfileView pipeline is provided in **Fig. 6** and its ten main steps are explained in detail below. A hands-on description of the ten steps for the CPF family is given in the **Supplementary File**. ProfileView takes as input a Pfam domain *D* and a set of homologous sequences *S* to be classified. If similar Pfam domains exist (Pfam usually names them with a numerical extension, as for instance HAD and HAD 2), then the user can decide to provide several alternative domains as input and construct the model library *M*_D_ accordingly.

**Figure 6:**
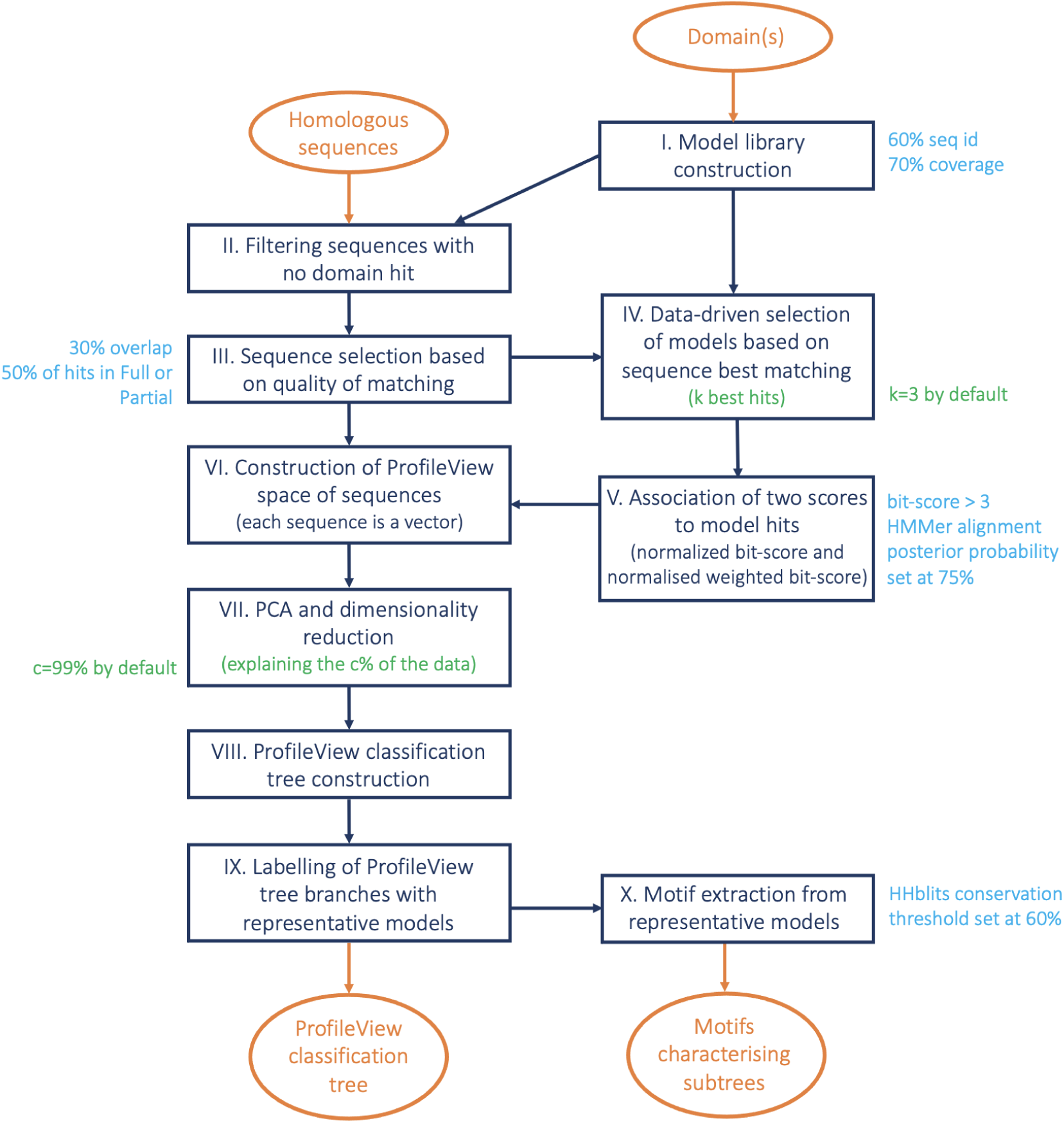
ProfileView flowchart. The ProfileView pipeline is organised in ten main steps: (I) building the model library for a domain or a few similar domains chosen by the user, (II) sequence filtering based on matching/unmatching of the models on a sequence, (III) sequence selection based on the quality of a match, (IV) filtering of models to reduce model redundancy, (V) association of two scores to each model hit, (VI) construction of the ProfileView space of sequences, (VII) dimensionality reduction of the sequence space, (VIII) construction of the ProfileView classification tree, (IX) identification of the best representative models for subtrees, (X) extraction of functional motifs from representative models. ProfileView parameters that the user can modify are highlighted in green, and those that remain fixed are highlighted in cyan.

#### I. Model library construction

To construct a library of models *M*_D_ for the domain *D*, we considered sequences from the FULL dataset in Pfam database (Finn *et al*., 2014) and, for each sequence, we built a CCM (Bernardes *et al*., 2016) by searching in Uniclust30 (which is UniProtKB clustered at 30% identity and for which a HH-blits database is provided; Mirdita *et al*. (2017)) for highly significant matches of homologous sequences having at least 60% identity with the query domain sequence and covering at least 70% of it. More precisely, a multiple sequence alignment is built using the command hhblits of the HH-suite (Remmert *et al*., 2011) (with parameters -qid 60 -cov 70 -id 98 -e 1e-10 and database uniclust30_2017_10) and subsequently converted into a pHMM with HMMER (Eddy, 1998) in order to perform a sequence-profile comparison. Moreover, a pHMM is considered only if it is trained with a minimum number of 20 sequences.

Note that the sets of Pfam sequences in the FULL dataset might be very large (some tens of thousands of sequences) and that we reduced their number to a few thousand sequences, by applying MMseq2 (at https://github.com/soedinglab/MMseq2; the easy-cluster command of mmseqs was used with parameter --min-seq-id, to set the minimum sequence identity for clustering, and parameter -c 0.8, to consider matches above this fraction of aligned/covered query/target residues) off the ProfileView pipeline, to cluster close sequences together and select, from each cluster, a representative sequence from which to generate a model, as above. We asked sequences in a cluster to have more than either 40 or 60% sequence identity (default set at 50%) depending on the protein family, in such a way that around 3000-4000 representative sequences could be identified for building the model library.

If several similar Pfam domains are considered, the procedure above will be applied to the Pfam sequences associated with all domains.

#### II. Sequence filtering

After building the set of models for *D*, we discarded from the input set of sequences *S*, all sequences against which no domain hit was found (independently of the hit score). *S* domain annotation is carried out by considering HMMER best hits (version 3.1b2) for models in *M*_D_. Note that this step, based on multiple probabilistic models, is able to identify domains in divergent sequences where the consensus Pfam model cannot provide a hit. For all protein families, Table I (fourth column) reports the number of sequences after filtering. (See **Fig. S28A** for an illustration of sequence filtering.)

#### III. Sequence selection

Each CCM in *𝓜*_D_ is mapped against the set *S* of all input sequences using HMMER. Let *H* = *{h_s,m_ | s* ∈ *S, m* ∈ *𝓜*_D_*, score*(*h_s,m_*) *>* 0*}* be the set of hits *h_s,m_* provided by hmmsearch, where *s* is a sequence of *S*, *m* is a model of *𝓜*_D_ and *score*(*h_s,m_*) is the bit-score assigned to *h_s,m_*. The bit-score is a log-odds ratio score (in base two) comparing the likelihood of the pHMM to the likelihood of a null hypothesis (*i.e.* an i.i.d. random sequence model). More formally,

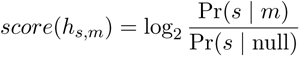

where Pr(*s | m*) is the probability of the pHMM *m* generating the sequence *s* and Pr(*s |* null) is the probability of *s* being generated by the null model (Barrett *et al*., 1997).

We partitioned the hit set *𝓗* in three subsets *Full*(*𝓗*), *Overlap*(*𝓗*), *P artial*(*𝓗*), where *Full*(*𝓗*) contains all hits that fully cover the associated model, *Overlap*(*𝓗*) contains all hits involving the extremes of a sequence covered only partially by the associated model (this situation corresponds to an “incomplete” sequence), and *P artial*(*𝓗*) contains all remaining hits. (See **Fig. S28B** for an illustration of the three matching types.) More formally, given a hit *h_s,m_* ∈ *𝓗*, it belongs to *Full*(*𝓗*) if the aligned region of *m* to *s* (excluding gaps) is at least 90% of the length of *m*. If *h_s,m_* represents an overlap between *s* and *m* (allowing an overhang length of at most the 10% of the sequence length) then *h_s,m_* ∈ *Overlap*(*𝓗*). Otherwise, *h_s,m_* ∈ *P artial*(*𝓗*).

To eliminate potentially incomplete sequences, a sequence *s* is retained only if:

1. either at most the 30% of its hits belong to *Overlap*(*𝓗*),
2. or, at least the 50% of its hits belong to either *Full*(*𝓗*) or *P artial*(*𝓗*).

These two conditions have been introduced in order to take into account the fact that Pfam might also contain partial sequences that could lead to the construction of very short models (that could be fully aligned in potentially incomplete sequences). We refer to the reduced set of sequences as *S*\*.

#### IV. Data driven selection of models based on sequence best matching

In order to restrict the analysis to a reduced set of models that remains representative of *𝓜*_D_, we kept only those models that achieve one of the *k* best scores for at least one sequence of *S*\*, for *k* = 3 (default). The rationale of this model filtering is to get rid of “noisy” models and, at the same time, significantly reduce the size of *𝓜*_D_, from some thousands down to a few hundreds. We refer to the reduced set of models as *𝓜*\*_D_. *k* is a parameter that can be set by the user.

#### V. Association of two ProfileView scores to model hits: the normalized bit-score and the normalized weighted bit-score

Let *L_s_* be the number of positions in a sequence *s* that match to a model *m* in a sequence/model alignment (that is, no gap is considered in the counting). Given a hit *h_s,m_* we define the following two scores for it:

• a normalized bit-score 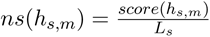

• a normalized weighted bit-score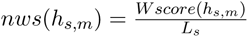, where *W score*(*h_s,m_*) is the sum of bit-scores over the positions in the sequence-profile alignment where the bit-score is greater than 3 (that is, the positions where *m* and *s* strongly agree). More formally, let 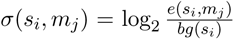 be the log-odds ratio of a residue *s_i_* being emitted from a match state *m_j_* with emission probability *e*(*s_i_, m_j_*) and with null model background frequency *bg*(*s_i_*), defined by HMMER during the model construction and differing between amino acids (Eddy, 1998). Given the list ((*s_i_*_1_ *, m_j_*_1_)*,…,* (*s_iK_, m_jK_*)) of the aligned residues of *s* against the model states of *m* and such that the posterior probability, computed by HMMER, of each aligned pair is greater than 75%, we define 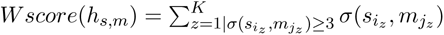.

Both scores are computed for all hits *h_s,m_* and used to construct the ProfileView space of sequences.

#### VI. The construction of a ProfileView space of sequences

For each sequence *s* ∈ *S*\*, we construct a vector *v_s_*, where the dimension of *v_s_*is 2*|𝓜*\*_D_| and *|𝓜*\*_D_| is the number of models in *𝓜*\*_D_. The vector *v_s_* contains the pairs of values *ns*(*h_s,m_*) and *nws*(*h_s,m_*), for each *m* ∈ *𝓜*_D_* . If a model *m* does not have a hit on the sequence *s* ∈ *S*\*, then we assume that *h_s,m_ /*∈ *𝓗* and let *ns*(*h_s,m_*) = 0 and *nws*(*h_s,m_*) = 0. Hence, we say that the ProfileView space *PV* is a 2*|M*\*_D_|-dimensional space, where each dimension is associated with either the normalized bit-score or the normalized weighted bit-score for some model *m* ∈ *𝓜*\*_D_. Each sequence is a point in *PV* and its position reflects the proximity of the sequence to CCMs in *𝓜*\*_D_.

#### VII. PCA and dimensionality reduction for ProfileView space of sequences

After constructing the ProfileView space *PV* for the sequences *s* ∈ *S*\*, Principal Component Analysis (PCA) is performed to reduce its number of dimensions. More precisely, *PV* is reduced to a *p*-dimensional space *PV*\*, where *p* is the minimum number of principal components that explain the *c*% of variance for the set *S*\*. By default, *c* = 99%. This value should decrease the number of dimensions to a few dozens. If a protein family is characterised by a large diversity of representative sequences, the user may have to loosen the constraints on variance by setting *c* to smaller values. *c* is a parameter that can be set by the user.

#### VIII. The ProfileView tree construction

Sequences are clustered in *PV*\* using a hierarchical agglomerative strategy. Namely, we considered the Euclidean distance between vectors and Ward’s minimum variance method for merging clusters. The logic of this criterion is to select, at each step, the pair of clusters that minimize the total variance within the cluster after the merging. Starting from all clusters being singletons, this bottom-up algorithm completes in *|S* \**| −* 1 agglomerative steps and allows to represent clusters in a hierarchical way and to define a rooted tree. More precisely, it produces a binary tree where each internal node defines a cluster of two or more elements (according to the chosen merge criterion). Moreover, in such a tree, the distances/dissimilarities between the merged clusters are encoded as edge weights.

#### IX. Association of representative models to ProfileView subtrees

To better explore subtrees in the ProfileView tree, potentially associated with known functions, we associated a *representative model* to the sets of sequences that label their leaves. Intuitively, a representative model separates a subset of sequences *𝓒* from the rest of the sequences of the tree (this set is designated *S*\* *\ C*) in the ProfileView space *PV*\*. Given a model *m* in the library, let us call *𝓒_m_*\* the maximal subset of *𝓒* where the model assigns higher scores to sequences in *𝓒_m_*\* than to sequences in *S*\* *\ C*. This must apply to at least one of the metrics – *ns* and *nws* – which define *PV*\* (see step III). For each model *m* in the library, we compute *𝓒_m_*\* and choose the model with a *𝓒_m_*\* of largest cardinality as the *representative model* of *𝓒*. If two models have the same maximum cardinality, we choose the model *m* that provides the best separation, i.e. the model that maximizes the distance between the centroids of the sets *𝓒_m_*\* and *S*\* *\ C* (again, computed according to the *ns* and *nws* metrics). If *𝓒* is the set of sequences of a subtree *T* of the ProfileView tree (which is not the entire tree), then a *representative model m for C is associated with the root of T* when the following two conditions are met: 1. *𝓒_m_*\* includes at least half of the sequences in *𝓒* and 2. *𝓒_m_*\* contains at least one sequence from each of the child subtrees of *T* . Note that a node in the ProfileView tree might be left without a representative model. When ProfileView returns a representative model for a node of the tree, it also returns a list of suboptimal models covering either the same amount of sequences *|𝓒_M_*\* | or 90% of *|C|*.

#### X. Motifs extraction from representative models

A motif extracted from a representative model is the set of all amino acids characterizing well-conserved columns (*i.e.* match states) in the sequence alignment associated with the model, according to the hhblits’ definition. That is, given a column of the multiple sequence alignment related to the model, an amino acid is *well-conserved* if it occurs with a probability *≥* 0.6 before adding pseudo-counts and including gaps in the fraction count.

### Parameters used in ProfileView analysis of the seven protein families

For all protein families, the domain(s) considered for model construction, their accession code and the number of constructed models are reported in **Table I**. For the model construction, representative sequences for FAD and GH30 were retrieved from Pfam v31 while for all other domains, we used Pfam v32. For the three families, Glycoside Hydrolase family 30 (GH30), HAD/*β*-PGM/Phosphatase-like and B12-binding domain containing, Pfam contains two similar domains (see Table I, sixth column) and we used Pfam sequences from both of these domains, we clustered them and built the associated models. For the three families, using two domains instead of one improves the classification slightly.

ProfileView was run with the same default parameters *k* = 3 and *c* = 99%, for all protein families in Table I, with the exception of the WW domain, which is characterised by a wide variability of sequences, that run with *k* = 5 and *c* = 80% (see steps IV and VII). For WW domains, note that with *c* = 80%, we have obtained a space of 11 dimensions, against 206 dimensions obtained with a threshold of *c* = 99%, starting from a total of 2488 dimensions. Also, the number of best matching models increased to *k* = 5 allowed us to obtain 1244 models versus 845 obtained with k=3. The idea being that when the dataset of sequences to be classified is very diversified, as for the WW domain family, the number of models should be large (*>* 1000) to explain diversity.

For the SPASM/twitch domain containing family subgroup, we performed two independent analyses, one based on the Radical-SAM domain and the other based on the SPASM domain. This is because all the original sequences contained the Radical-SAM domain but not all contained the SPASM domain.

### Motifs graphical representation

Model logos were built using the python package of Weblogo (Crooks *et al*., 2004) (version 3.7) which allowed us to easily export sequence logos (Schneider and Stephens, 1990). Amino acids are colored according to chemical properties: neutral polar amino acids (G, S, T, Y, C) show in green, acidic polar (Q, N) violet, positively charged (K, R, H) blue, negatively charged (D, E) red, and hydrophobic (A, V, L, I, P, W, F, M) black.

The graphical representation of a motif associated with some representative model was augmented by extra information helping to easily compare the motif across representative models. Namely, we highlighted, by a coloured “dot”, positions in it found to be well-conserved in other representative models. Given a reference model *M_r_* and a query model *M_q_*, a dot is put under a well-conserved column of *M_r_*, if there exists a column in the query model *M_q_* : 1. aligning in hhblits with a score greater than +1.5 (*i.e.* fairly similar amino acid profiles) and posterior probability greater than 0.8; 2. containing a most conserved amino acid which is the same as in *M_r_* and is also well-conserved. A circled dot indicates an aligned column in *M_q_* satisfying 1 but not 2. This means that the most conserved amino acid in *M_r_* shows *<* 60% frequency in *M_q_* . Note that, in this case, *M_r_* and *M_q_* might display different most conserved amino acids.

It is important to notice that given two models and a position, the score assigned to that position in the hhblits pairwise alignment of the models depends on the reliability of the query-template alignment (https://github.com/soedinglab/hh-suite/wiki). Depending on which one of the models is considered as a template, the scores assigned to the same position might vary (the confidence values are obtained from the posterior probabilities calculated in the Forward-Backward algorithm of hhblits). In particular, hhblits is warning that the confidence score for an aligned position depends on the confidence on the alignment of the close by region. As a consequence, the alignment score of certain conserved position might decrease because of the presence of a very variable region in their vicinity, possibly containing gaps. This explains why, for aligned positions of two motifs, we might miss to indicate related positions or we might display different color dots. An example of missing related positions is illustrated by position 102 in the NCRY motif and position 103 in the plant PR CRY motif of CPF. The two motifs clearly diverge within the region just following positions 102/103, justifying a difficult model alignment and a low confidence score for 102/103. A second example, illustrating the asymmetry of the coloured dots, is position 102 in the NCRY motif aligned with position 95 in CRY Pro. While the CRY Pro motif records the coloured dot for a matching with NCRY, this is not true for the NCRY motif. Indeed, while the two positions align together with a confidence score of 0.8 for the CRY Pro model taken as a template, they also align together when the NCRY model is taken as the template but with a confidence score dropping at 0.6.

### Phylogenetic tree construction for CPF, FAD and WW sequences

The multiple sequence alignments of CPF sequences and FAD sequences were computed using MUSCLE version v3.8.31 (Edgar, 2004), and were then trimmed using trimAl version 1.4.rev22 (Capella-Gutíerrez *et al*., 2009) with a gap cutoff of 0.01 (*i.e.* columns containing more than 99% of gaps were removed). Then, for each sequence alignment, we selected the best evolutionary model using ProtTest (version 3.4.2) (Darriba *et al*., 2011). More precisely, the evolutionary model best fitting the data was determined by comparing the likelihood of all models according to the Akaike Information Criterion (AIC). The model optimisation of ProtTest was run using a maximum-likelihood-tree strategy and the tree generated for the best-fit model (VT+G+F) was considered as input for the construction of the final phylogenetic tree (with parameter *α* = 1.061). In particular, the construction of a maximum-likelihood phylogenetic tree has been carried out with PhyML 3.0 (Guindon *et al*., 2010) that optimized the output tree with Subtree-Prune-Regraft (SPR) moves and considering the SH-like approximate likelihood-ratio test. Finally, branches with a support value smaller than 0.5 were collapsed. The phylogenetic tree for the set of homologous CPF sequences used to validate ProfileView is reported in **Fig. S3** and contains 307 leaves corresponding to the 307 CPF sequences containing the FAD binding domain. The phylogenetic tree for the set of 307 FAD sequences is reported in **Fig. S4**.

The procedure used to generate the phylogenetic tree for WW domain sequences is the same as the one used for CPF and FAD sequences. The best-fit model (computed with ProtTest) is RtREV+I+G, with parameters *α* = 1.647 and *p − inv* = 0.028.

Phylogenetic and ProfileView trees have been generated with iTOL (Letunic and Bork, 2019).

### Output files of ProfileView

ProfileView produces several output datasets: the model library, the ProfileView tree, the list of representative models associated with internal nodes of the tree.

Also, ProfileView provides to the user the possibility to choose a list of representative models to be compared. The first model of this list is considered as a reference model. A first output describes and provides the logo reporting all conserved positions together with a list of coloured dots (possibly circled) obtained after a pairwise comparison of a model in the list with the reference model (see Methods above; see for example **Fig. 1D**). A second output describes and provides the logo reporting an intermediate representation of the positions in the reference model, namely reporting all conserved positions in the associated motif and all positions that are not conserved in the reference model but that are conserved in some other model in the list.

### 1.1 Comparison with other tools

CUPP (Barrett and Lange, 2019) and PANTHER (Mi *et al*., 2012, 2013) have been used for comparison. CUPP v1.0.14 was run with CUPPclustering.py and parameter -cluster to execute the clustering (http://www.bioengineering.dtu.dk/CUPP). The PANTHER HMM library version 15.0 and the pantherScore2.2 tool (scoring protein sequences against the library) were retrieved at http://www.pantherdb.org. We used pantherScore2.2.pl with parameters -l [PANTHER15.0 library] -D B -n, where -D B allows to visualise the best hit in the output and -n allows to visualize family and subfamily names in the output.

### Evaluation

For each protein family, we considered functionally characterised pools of sequences collected from the literature and classified them in groups with ProfileView. To evaluate what proportion of sequences with a specific function is correctly classified by ProfileView, we used the Recall measure, defined as TP/TP+FN, where TP is the number of sequences that ProfileView classifies correctly and FN is the number of sequences that it classifies in the wrong group. The idea is to evaluate in which manner ProfileView captures as many positives as possible.

### Computing time

The most costly computational part of the pipeline is the construction of the probabilistic models for a protein sequence. The program was tested using 16 threads on a single machine equipped with an Intel Xeon E5-2670 CPU running at 2.60GHz, with 128 GB of RAM, and a Linux operating system (CentOS release 6.5). **Table S2** summarizes, for each protein family, the time complexity for the model library construction and the classification step. The time used for the model library construction depends on the number of models and the length of the domain. Once a library is constructed, it can be used for the analysis of different protein families. Note that the same library constructed for the Radical SAM domain was used for both the analysis of the Methylthiotransferase family and the SPASM/twitch domain containing family. Note that, for the WW domain family, (Tubiana *et al*., 2019) indicates about 1-2 days of computing time on an Intel Xeon Phi processor with 2 *×* 28 cores to run RBM analysis. ProfileView classifies this family in less than 9 hours (**Table S2**).

### Implementation and software availability

ProfileView has been developed and tested under a UNIX operating system, using Bash, Python, and R scripts. It exploits GNU parallel (Tange, 2018), if available on the system, in order to perform some jobs in parallel. It is implemented in three main parts carrying out the following modules of the pipeline: the construction of a single-domain model library, the generation of the ProfileView tree along with its representative models, the comparison of selected representative models and the identification of conserved positions/motifs. ProfileView is available at http://www.lcqb.upmc.fr/profileview/ under the version 2.1 of the CeCILL Free Software License.

### Data accessibility

The set of sequences used in the analysis and the model libraries, distance trees, ProfileView trees generated and discussed in the article (Cryptochrome/Photolyase Family, the WW domain family, the glycoside hydrolase enzymes family GH30, the four enzyme superfamilies of the Structure-Function Linkage Database) are available at http://www.lcqb.upmc.fr/profileview/.

## Discussion

The availability of large quantities of (meta)genomic data is allowing for a deeper exploration of living organisms and of the processes underpinning their genetic, phylogenetic and functional diversification. Computational approaches, able to highlight these diversities and to identify what is functionally new within the realm of sequence information, will make the first fundamental step in the discovery of new candidates to be experimentally tested for their functional activity. Moreover, due to the huge quantity of sequences to be acquired in years to come (1 zetta-bases/year are expected in 2025 (Stephens *et al*., 2015)), there will be no more way to look into this mass of data with an “expert eye” and computational approaches will play a key role on the extraction of novel information and in functional classification.

Today, we can characterise homologs based on their similarity through distance measures modelling the evolution of the entire sequences. However, as shown here and elsewhere (Schnoes *et al*., 2009; Mi *et al*., 2012; Akiva *et al*., 2014; Barrett and Lange, 2019), this computational approach is insufficient to provide insights on protein functional activities, and a large number of sequences remain not yet functionally annotated. Some of these protein families, like the seven families discussed in this study, are extremely important in medicine, biology, environmental science and biotechnology due to their key roles in cancer biology, DNA repair, drug delivery strategies, chronobiology and photobiology, specific enzymatic reactions, the formation of protein-protein interaction networks, optogenetics. Thanks to their key role, for decades now, experiments have accumulated a huge amount of functional information that we used to validate the ProfileView approach. ProfileView functional organisation of these seven families agrees with experimental evidence.

ProfileView highlights that protein functional classification depends on a non-linear contribution of many probabilistic models and that conserved patterns in sequences are not sufficient alone to discriminate diversified functions of complex protein families. This change of perspective in functional classification, underlies the complexity of the question and explains why this problem is wide open today despite the clear interest in classifying protein families that have been amply studied in molecular biology, like transporters, signalling, transcription factors.

By constructing multiple probabilistic profiles characterising different conserved motifs in homologous domain sequences, ProfileView captures functional signals and, by combining them, is able to successfully classify large datasets. The main advantages of ProfileView approach compared to those developed before are as follows: (i) it is alignment-free and avoids errors due to the difficulty of comparing distant homologues; (ii) several probabilistic models represent more precisely than a single consensus models the functional variability of protein families; (iii) large quantities of data are not needed to learn features and run the classification; (iv) functional annotation of many sequences does not need to be known to explore with precision the space of sequences and classify them; (v) it is a general approach applicable to proteins of arbitrary length and function. Moreover, once a domain library is constructed, ProfileView is computationally efficient in screening very large sets of homologous sequences in a reasonable time.

ProfileView demonstrated to discover potentially interesting CPF proteins whose function could be experimentally tested with the purpose of enlarging our understanding of the mechanisms exploiting light to perform functional activities in natural environments. These proteins are of interest for biotechnology and any computational approach to highlight them is desired. It also organised the WW domain family in subtrees of sequences, corresponding to a large spectrum of differences in binding affinity to various ligands, which have been experimentally observed. It demonstrates that a large variety of sequence motifs covers this spectrum and it identifies these motifs. It could classify protein superfamilies in the manually curated CAZy and SFLD databases by accurately identifying differences in their multiple enzymatic reactions. Compared to Tubiana *et al* (Tubiana *et al*., 2019), a computational approach also based on sequence analysis, it describes differences among binding motifs in much greater detail, opening new avenues in the discovery of alternative binding patterns in protein-protein interaction networks. It has been compared favorably to other classification tools like PANTHER and CUPP, on the CPF, the WW domains and the GH30 family classified in the Carbohydrate-Active Enzymes database CAZy.

On the methodological side, ProfileView addresses the problem of extracting biological information on protein families from the huge space of natural sequences, and the sampling of distant sequences could be realised using different distance measures. This is an important direction of investigation possibly leading to more refined biological information extracted from sequences.

From the algorithmic point of view, ProfileView is surprisingly simple compared to the Restricted Boltzmann Machines (RBM) model used in (Tubiana *et al*., 2019) to classify WW domain homologs. RBM, is a generative stochastic (single layer) artificial neural network that learns collective modes by extracting short sequence motifs from sets of sequences based on correlation patterns among alignment positions. These motifs might reveal structural, functional and phylogenetic features and they are used to define a representation space where to classify sequences. RBM generative nature makes training challenging by an algorithmic point of view since it requires intensive sampling from large training sets. In contrast, ProfileView constructs probabilistic profiles from close neighbours of distant homologous sequences (demanding a very small number of sequences, a minimum of 20) in sequence space, making no use of positional correlations nor of their generative modelling. Its probabilistic models encode conserved patterns ignoring those parts of the homologous sequences appearing variable (see discussion on the two CPF sequences U5NDX3 and R7UL99 above). The number of models is not a restriction for the construction of the classification space. A possible direction of investigation is the design of multiple layers (of models) for an architecture that analyses finer motifs as well as proteins comprising multiple domains.

The fine understanding of functional mechanisms might need more sophisticated computational approaches than ProfileView. For instance, for CPF classification which is based on the FAD binding domain, ProfileView highlights functional differences between large classes of CPF sequences, helping to model the proximity between these classes with an appropriate identification of a functional tree topology. To find functional differences within classes and to anticipate the existence of a double function (see **Fig. S1**), the entire CPF sequence might be necessary, possibly because of the interaction between domains which might have functional consequences as highlighted in (Rosensweig *et al*., 2018).

Last, even if ProfileView has been applied here to the classification of entire protein sequences, it can handle metagenomics sequences as well. In this respect, it is important to highlight that the majority of metagenomics and metatranscriptomics data come from organisms that cannot be cultured and that will, possibly, never be isolated. Hence, conceptual new approaches to explore their biology in complex ecosystems is desperately needed. ProfileView allows to increase knowledge on the biology of organisms whose ecological role has been recognised (*e.g.* marine microbes) but that are still not accessible to functional investigations, opening a new avenue to functional exploration.

## Availability of data and materials

The set of sequences for CPF, FAD, WW domain, Glyco-hydro-30 and Glyco-hydro-30-2 for GH30, HAD and HAD 2 for Haloacid Dehalogenase, B12-binding and B12-binding 2 for B12-binding domain containing, Radical SAM for Methylthiotransferase and SPASM/twitch domain containing subgroups, and SPASM for SPASM/twitch domain containing subgroup, model libraries, phylogenetic trees, ProfileView trees are available at http://www.lcqb.upmc.fr/profileview/.

## Authors’ contributions

RV and AC conceived and designed the experiments. RV performed the experiments. EL performed the structural analysis of CPF classes. RV, JPB, AF and AC analyzed the data. AC, RV, JPB and AF wrote the paper. All authors read and approved the final manuscript.

## Acknowledgements

We thank Simona Cocco and Jérôme Tubiana for providing to us the dataset of WW sequences used in their study.

## SUPPLEMENTARY FILE

### 1. Functional diversification of CPF

Photolyases are photoactive enzymes that bind DNA and use blue light to mend two different types of UV-induced DNA damage, either ss/dsDNA cyclobutane pyrimidine dimer (CPD) or (6-4) pyrimidine-pyrimidone photoproducts, and are thus classified as either CPD or (6-4) photolyases. Moreover, some photolyases such as CRY-DASH can only bind and repair ssDNA CPD. The weaker DNA photolyase activity of CRY-DASH has been reported as a consequence of lower DNA binding affinity than CPD photolyase (Sato *et al*., 2018). Cryptochromes (CRY) do not bind DNA and are mainly photoreceptors (PR) (specifically noted PR CRY in the following) involved in many biological responses to light (*e.g.* photomorphogenesis, entrainment of the circadian clock). However, some CRYs are also light-independent transcriptional regulators taking part in the central circadian oscillator generating biological rhythms.

In the last decade, new CPF variants, exhibiting different photobiological properties or functions, have been discovered, changing current views on their evolution (Jaubert *et al*., 2017) and functional diversification (Coesel *et al*., 2009; Heijde *et al*., 2010; Fortunato *et al*., 2015; Essen *et al*., 2017). The initially proposed functional separation between CRYs and PLs has gradually started to vanish, as there are now several examples of CPF members exhibiting both functions (Coesel *et al*., 2009; Heijde *et al*., 2010; Franz *et al*., 2018). Some CPF members have even been used for optogenetic applications (Ozkan-Dagliyan *et al*., 2013; Liu *et al*., 2012) or proposed as magnetoreceptors (Rodgers and Hore, 2009).

Although a lot of experimental progress has been made, CPF functions could not be anticipated by the analysis of domain organisation due to a very simplified architecture of the CPF sequences, nor by structural properties due to the high similarity of their protein structures, nor by primary protein sequences. Tools employed for the phylogenetic reconstruction of this family (Chaves *et al*., 2006; Lucas-Lledö and Lynch, 2009; Mei and Dvornyk, 2015; Ozturk, 2017) did not allow to resolve different functions (*e.g.*, light-dependent DNA photolyases and light-independent transcriptional regulators) or to anticipate the function of new CPF sequences.

### 2. ProfileView algorithm applied to the CPF family

A hands-on description of our methodological approach is provided here for the analysis of the cryptochrome/photolyase family (CPF). ProfileView bases the analysis on the FAD binding domain, occurring in all CPF sequences, and considers the set *S*_CPF_ of 397 CPF sequences spanning the whole phylogenetic tree, of which 69 are functionally characterized CPF homologs and the remaining ones are known functionally uncharacterised sequences. The ProfileView pipeline, comprising ten main steps, is illustrated by the flowchart in **Fig. 6**.

#### I. Model library construction

ProfileView constructs a library, *M*_FAD_, of probabilistic models (Eddy, 1998) for the *FAD binding domain of DNA photolyase* from Pfam version 31 (accession code PF03441), due to its functional importance for CPF activity (**Fig. 1** and **Fig. 6**). More in detail, we considered all 4615 sequences which belong to the FULL alignment in Pfam. For each one of them, a CCM has been constructed with the command mentioned above. Finally, our model library *M*_FAD_ for the FAD-binding domain comprises 3735 CCMs, because for some sequences we could not collect a minimum of 20 homologs. The pipeline for *M*_FAD_ construction is depicted in **Fig. 1A**.

The set of Pfam sequences used to construct the ProfileView’s model library for CPF is mainly different from the set of classified sequences: among the 240 models taken into account for the classification of the 307 sequences, just 17 of these models were built from a (Pfam) sequence in *S*_CPF_ and only one is a representative model (for the (6-4) PL subtree) in the ProfileView tree. Moreover, the average identity and similarity (based on pairwise alignments) between the set of 307 sequences to classify and the set of the 240 sequences generating the models are 26.35% and 36.73%, respectively.

#### II. Sequence filtering

After building *M*_FAD_, we discarded from *S*_CPF_ all sequences against which we were not able to find any domain hit (independently of the hit score). *S*_CPF_ domain annotation was carried out by considering HMMER best hits (version 3.1b2) for all models in *M*_FAD_. An a posteriori phylogenetic analysis of the original set of CPF sequences has been carried out with RAxML version 8.2.11 (with parameter -m PROTGAMMAAUTO). We observed that the set of discarded sequences, presenting no FAD binding domain match, correspond to long branches in the tree (see **Fig. S28A**). This preliminary filter led us to consider a set of 386 CPF sequences over the 397 we started with.

#### III. Sequence selection

The 386 CPF sequences are then selected further by considering the full set of models and evaluating the strength of their hits against the sequences. This testing is intended to discard sequences that end with just a fragment of the domain (see **Fig. S28B**). The corresponding hits are expected to be very weak and this concerned 79 CPF sequences. We remained with 307 sequences corresponding to the leaves of the ProfileView tree. We refer to this reduced set as *S*_C_* _PF_.

#### IV. Models filtering

The rationale of this model filtering is to get rid of “noisy” models and significantly reduce the size of *M*_FAD_ down to a few hundred models. For the 307 CPF sequences, we extracted the three models in *M*_FAD_ that best match the sequence and make the union of all of them. Many CPF sequences are best matched by the same models and the final set is comprised of 240 models that best identify the presence of the FAD binding domain in CPF sequences. We refer to this reduced set as *𝓜*\*_FAD_.

#### V. Association of two ProfileView scores to model hits: the normalized bit-score and the normalized weighted bit-score

To each hit, between a sequence *s* in *S*_C_* _PF_and a model *m* in *𝓜*\*_FAD_, we applied the definitions of the two scores given in Methods and used all the scores to represent the sequence *s* as a vector of 480 dimensions.

#### VI. The construction of a ProfileView space of sequences

This step contains the central idea of the ProfileView method: each CCM is matched to each sequence to be classified and the scores of the hits (see columns of real numbers in **Fig. 1B**) will provide a description of how close the model is to each sequence. In its turn, a sequence can be represented by how close all models are to it through a vector of scores (see rows of real numbers in **Fig. 1B**). In this way, we define a representation space of sequences that does not reflect sequence similarity but, instead, the closeness of each sequence to each model. Since a match of a model is evaluated by two scores (see V), the space will be a 480-dimensional space and each sequence will be a point in the space.

#### VII. PCA and dimensionality reduction for ProfileView space of sequences

For this step, we used the parameter *c* = 99%. From 480 dimensions, the reduction produced a space of 37 dimensions.

#### VIII. The ProfileView tree construction

This step classifies the set of protein sequences in the 37-dimensional space. For the generation of our ProfileView tree, we use a hierarchical clustering algorithm which allows to build a tree that groups together the 307 sequences. The ProfileView tree built for the CPF sequences is depicted in **Fig. S1**, where internal colours are identified by representative models (see below) and external strips are associated with known functions (according to the literature, see **Supplemental File** for the detailed list of publications).

#### IX. Association of representative models to ProfileView subtrees

We associate several representative models to subtrees of the ProfileView tree following the procedure detailed in Methods. **Fig. S1** indicates which nodes of the CPF tree are represented by a model.

#### X. Motifs extraction from representative models

We extracted from each representative model in **Fig. S1** associated with the colored functional subtrees, their corresponding functional motifs. They represent the specificity of the sequences within each subtree.

### 3. Identification of known key residues by comparison of representative motifs in CPF

The comparison of ProfileView motifs’ positions versus experimentally characterized positions is reported in our manually curated list “CPF mutants used for validation.xlsx”. **Table S3** indicates how many ProfileView positions are validated by current experimental evidence.

Some positions in a motif might be conserved also in other motifs (corresponding to other subtrees), but some positions are motif specific, as illustrated by the colored dots in the logos of **Fig. S5** and **Fig. S6** (see Methods). A number of observations are given below, especially for those highly specific positions that have not been reported in the literature before.

#### The transcriptional regulators motifs

As expected, most of the positions in motif #1, associated with the light-independent transcriptional regulator sequences (**Fig. 4A** and yellow subtree in **Fig. S1**), are conserved in other subtrees as well, notably (6-4) PL and animal PR CRY, because of the proximity of these subtrees in the phylogenetic tree. Highly conserved positions in most, if not all, models are clearly identified as highly conserved also in the Pfam model (**Fig. S7**). However, four positions (L6, N38, L42 and K44 in **Fig. 4A**) appear to be specific to light-independent transcriptional regulators. Three residues belong to the same helix (*α*12) and two of these positions (N38, K44) are known to belong to the interaction site with a partner and to the ubiquitination site (Hirano *et al*., 2013; Schmalen *et al*., 2014). The two remaining conserved positions (L6, L42), at the best of our knowledge, have not been identified before and open ways to new investigations. Similar considerations can be drawn on motif #2 associated with the light-independent transcriptional regulator sequences (**Fig. 4B**).

#### The (6-4) photolyase motif

The (6-4) photolyase motif generated by ProfileView highlights the highly specific amino acid L115 which interacts with DNA and belongs to the site of the damage DNA strand binding (**Fig. 4C**; see its specificity in **Fig. S5**). This position has not been discussed previously in the literature.

#### Comparison of classes I and III CPD PL motifs

By comparing the motif representing classes I and III CPD PL with those representing either class I CPD PL or class III CPD PL (see **Fig. S29**), we notice that there is almost no amino acid which is motif-specific among the three models. The strong closeness between motifs of class I and class III agrees with their shared function. We especially notice the conserved amino acids involved in CPD lesion binding sites such as W7, N71, M75, W114 and possibly F129 and Q134 (where numbers refer to the motif accounting for both classes I and III CPD PL; see “Class I & III CPD PL” motif and structure in **Fig. S29**), or those involved in FAD binding or FAD binding pocket such as R74, D102, D104, N108 (directly involved in the proton transfer to the FAD; see “Class I & III CPD PL” motif and structure in **Fig. S29**; see **Supplemental File**). This example demonstrates that the analysis of the different motifs at different nodes might be used to deduce common functions. However, it is possible to extract some differences among the two motifs, where specific amino acids such as W60 (for class III CPD PL) versus Y56 (for class I CPD PL) were suggested to make an alternative electron transfer pathway possibly important in some specific condition (Scheerer *et al*., 2015). Other interesting differences are W29, D63 and T64 from class III CPD PL which have been identified as interacting with the MTHF in a specific binding site of MTHF from class III CPD PL (Scheerer *et al*., 2015).

#### Comparison of classes I and III CPD PL and plant PR CRY motifs

Classes I and III CPD PL and plant PR CRY are well-studied families in terms of function and molecular mechanisms, and present numerous specific mutants leading to a loss of function. Remarkably, by crossing the functional characterisation of specific residues in the collections of mutants described in the manually curated list with our representative models of classes I and III CPD PL and plant PR CRY, we could validate 26 and 35 of the ProfileView positions from the 33 and 47 positions in the list, respectively (see **Table S3**).

#### Two new conserved positions for classes I, II and III CPD PL

Some promising new information can be extracted by the comparative analysis. Indeed, despite many studies on these PL classes, we could identify 2 specific amino acids (F27 and I55) with yet undefined function. When looking at their position in the structure, these amino acids do not seem to be directly involved neither in DNA nor in FAD binding. Nevertheless, they are highly specific suggesting their involvement in the CPD repair mechanism.

#### Comparison of classes I, II and III CPD PL and NCRY motifs

By comparing the three CPD PL motifs/models with the one from NCRY, we essentially remark the absence of the CPD binding site. In NCRY motifs, four amino acids involved in the interaction of the CPD lesions by photolyase, E4, N71, M75 and W114, are absent and respectively replaced by R4, F71, A75 and M114 suggesting the absence of binding affinity for CPD substrate.

#### Comparison of classes I, II and III CPD PL and plant PR CRY motifs

By comparing the three CPD PL motifs/models with the one from plant PR CRY, we also remark the absence of the CPD binding sites in plant PR CRY motif. Interestingly, at two specific positions of the CPD binding sites (M75, W114), two specific amino acids (V74, Y114) are found in the motif of Plant PR CRY which have been involved in the ATP binding site described up to now as specific for Plant PR CRY (Orth *et al*., 2017; Brautigam *et al*., 2004). Moreover, despite very conserved amino acids in class I CPD PL model such as D45, D49 or E54 (involved in the proton transfer to the FAD), the latter ones are present but not fully conserved in class III CPD PL and are clearly absent in the plant PR CRY model. This observation suggests that these amino acids might also be involved, directly or indirectly, in the CPD repair function, and that some variability is not expected to disrupt the function.

All these examples, experimentally validated by the genetic and functional analysis of selected mutations, illustrate the strength of ProfileView representative models in extracting important amino acids information from sequences that can be used to design tailored experiments for discovering new functional activities or novel biological mechanisms involving the FAD binding domain.

---

**Figure S1:**
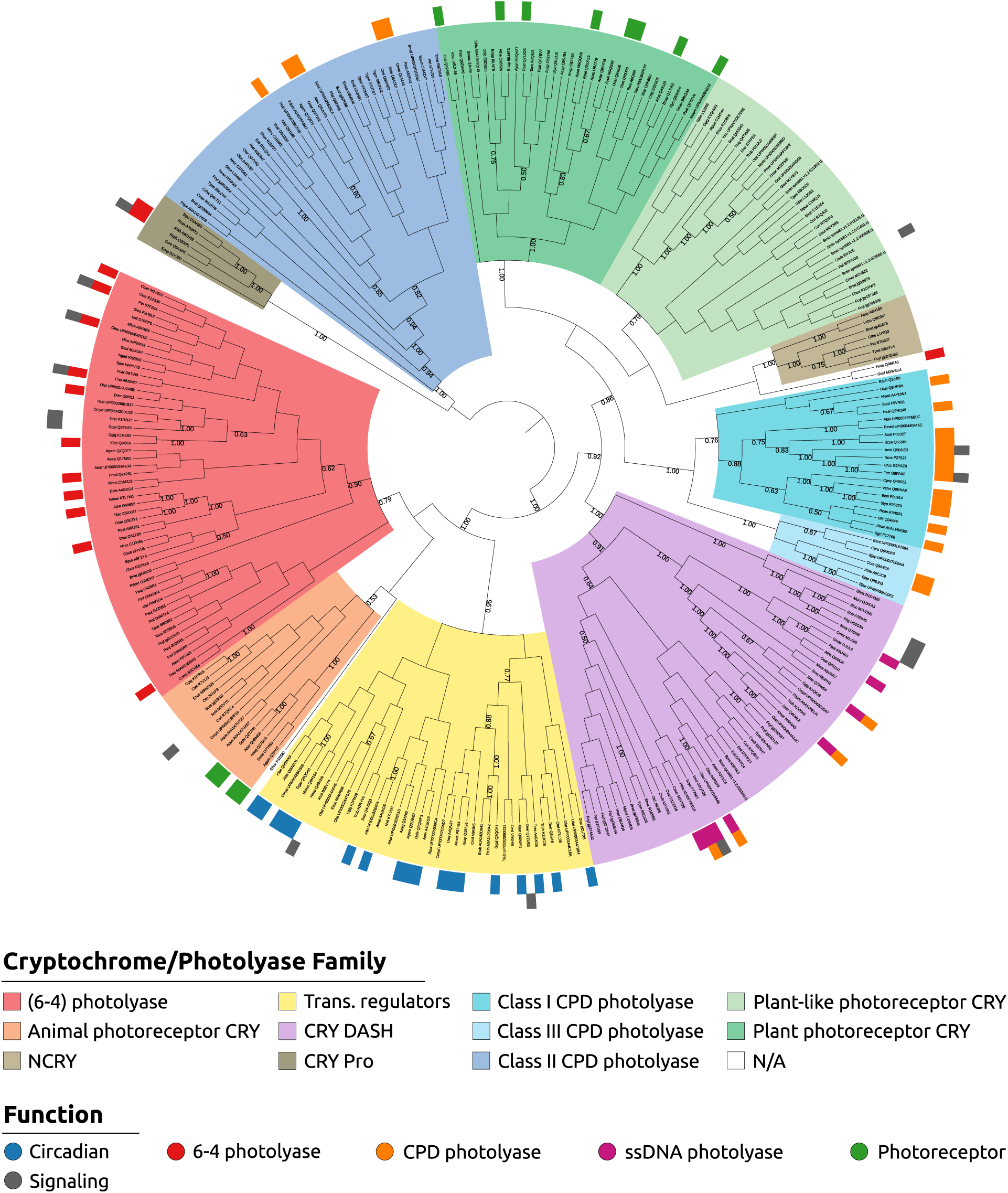
ProfileView tree of 307 FAD-binding domain sequences built from FAD-binding domain (PF03441) models from Pfam v31 using a hierarchical agglomerative clustering strategy. Colors of subtrees are identified by representative models and correspond to known CPF classes, with the exception of the NCRY subtree. External coloured labels define known functions for the sequences. Some of the 307 sequences are known to hold multiple functions and are labelled by two colors. The function “signalling” (grey) refers to signalling processes of different nature (photoreceptor, transcription, unknown). Numbers on the internal nodes correspond to the percentage of sequences in the corresponding subtree that are separated from the remaining sequences in the tree by the best representative model occurring in the model library.

**Figure S2:**
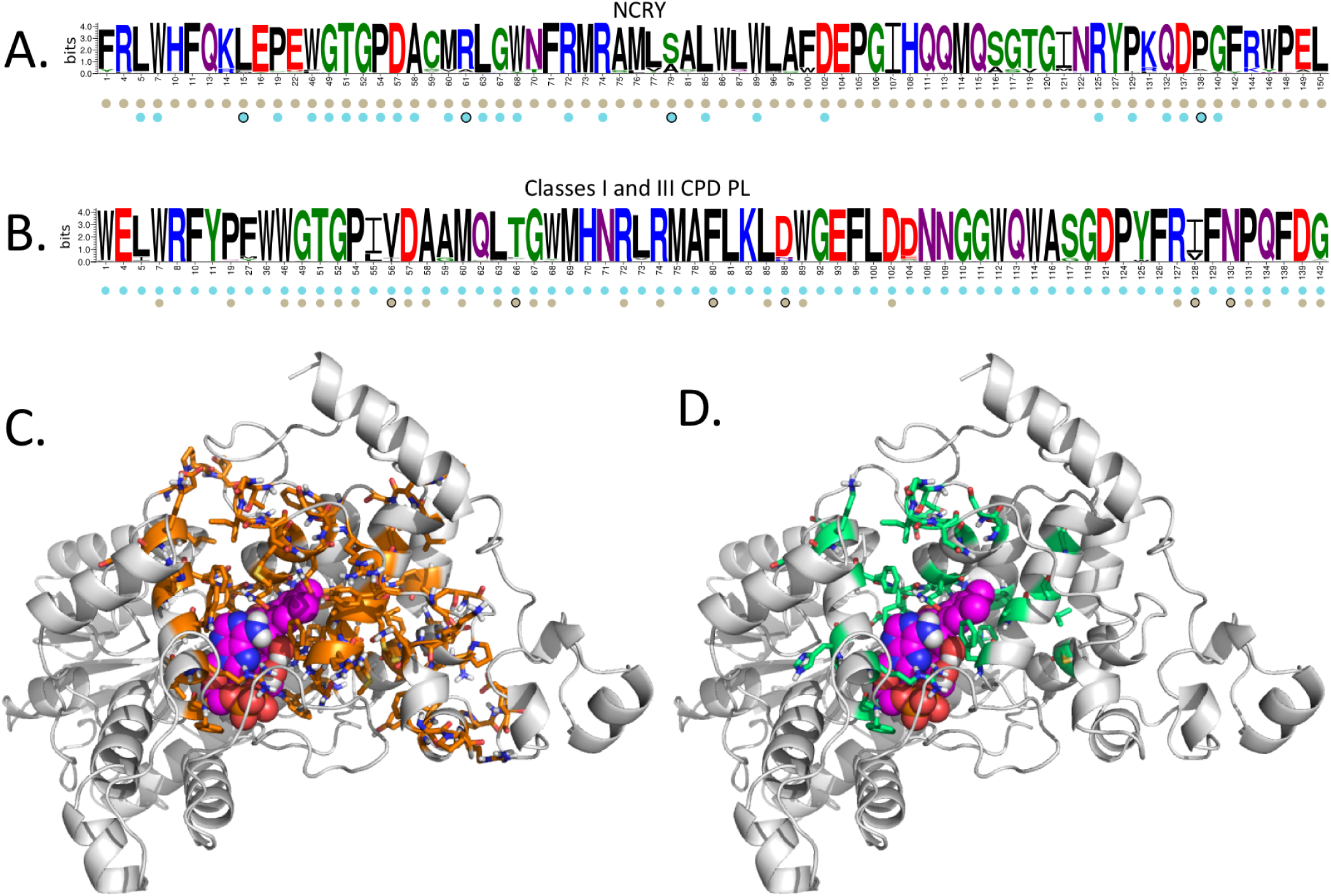
Motif for the NCRY sequences of the CPF family and their structural modelling. **A**. Motif resulting from the representative model of NCRY sequences (see **Fig. 3**). The positions of this model that are also conserved in the model representative of the class I CPD PL sequences in B are indicated with cyan bullets, below the motif. **B**. Motif resulting from the representative model of classes I & III CPD PL sequences (see **Fig. 3**). The positions of this model that are also conserved in the model representative of the NCRY sequences in A are indicated with beige bullets, below the motif. **C.** Homology model of the FAD *Pt*NCRY structure where all conserved residues in the NCRY motif (all positions making the motif in **A**) are highlighted in orange. Compare with the NCRY specific residues highlighted on the same homology model in **D**. **D.** All conserved residues in the NCRY motif that are NCRY specific (all positions in the motif in A marked with a beige bullet but not with a cyan one) are highlighted in green. Compare with **C**.

**Figure S3:**
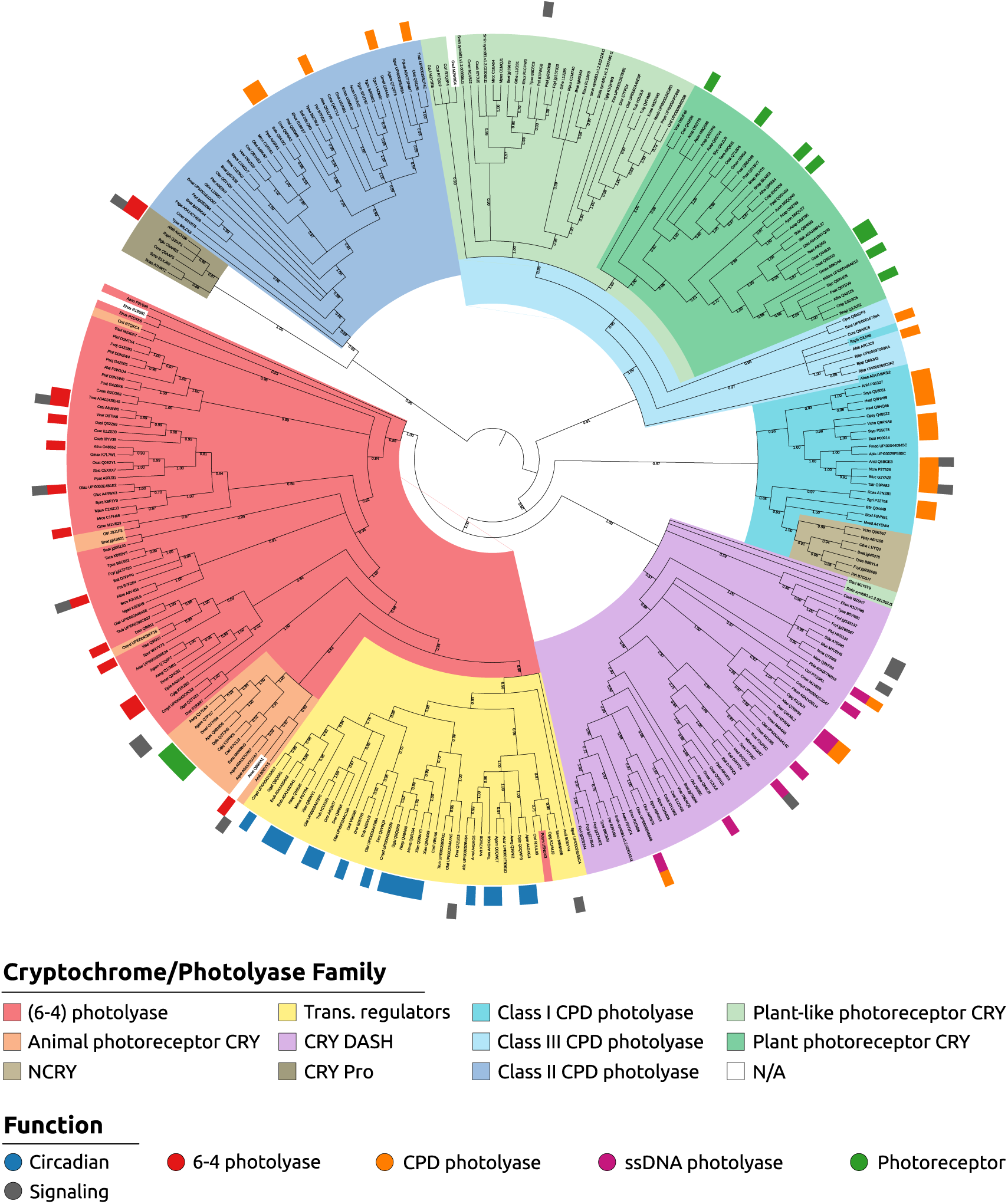
Phylogenetic tree constructed from 307 CPF sequences. Each sequence in the phylogenetic tree is coloured as in the ProfileView tree. Colors of internal subtrees are induced by sequence coloring. External labels report known functions for the sequences (see legend of **Fig. S1**). Numbers on the branches are bootstrap values.

**Figure S4:**
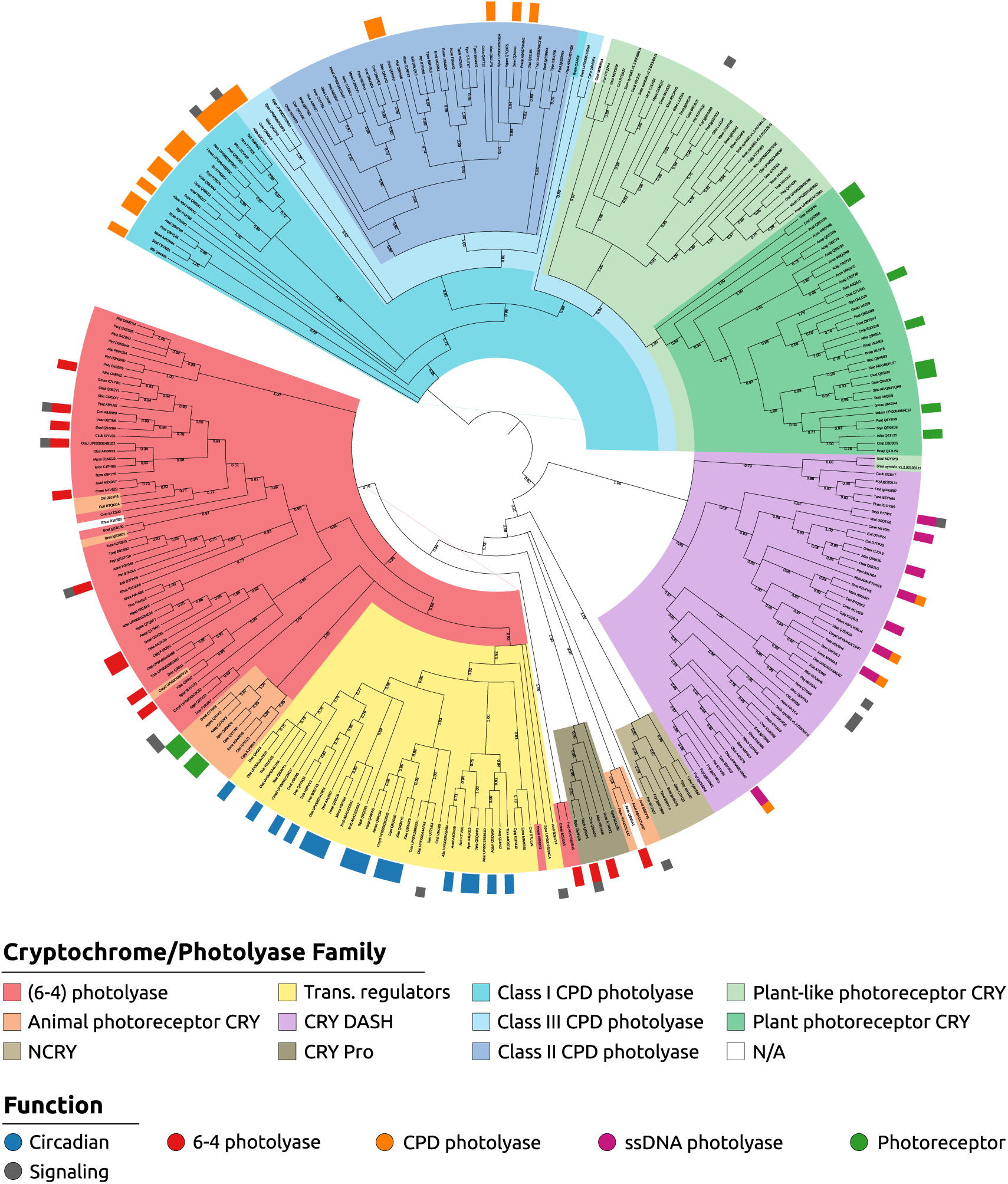
Phylogenetic tree constructed from 307 FAD-binding domain sequences. Each sequence in the phylogenetic tree is coloured as in the ProfileView tree. Colors of internal subtrees are induced by sequence coloring. External labels report known functions for the sequences (see legend of **Fig. S1**). Numbers on the branches are bootstrap values.

**Figure S5:**
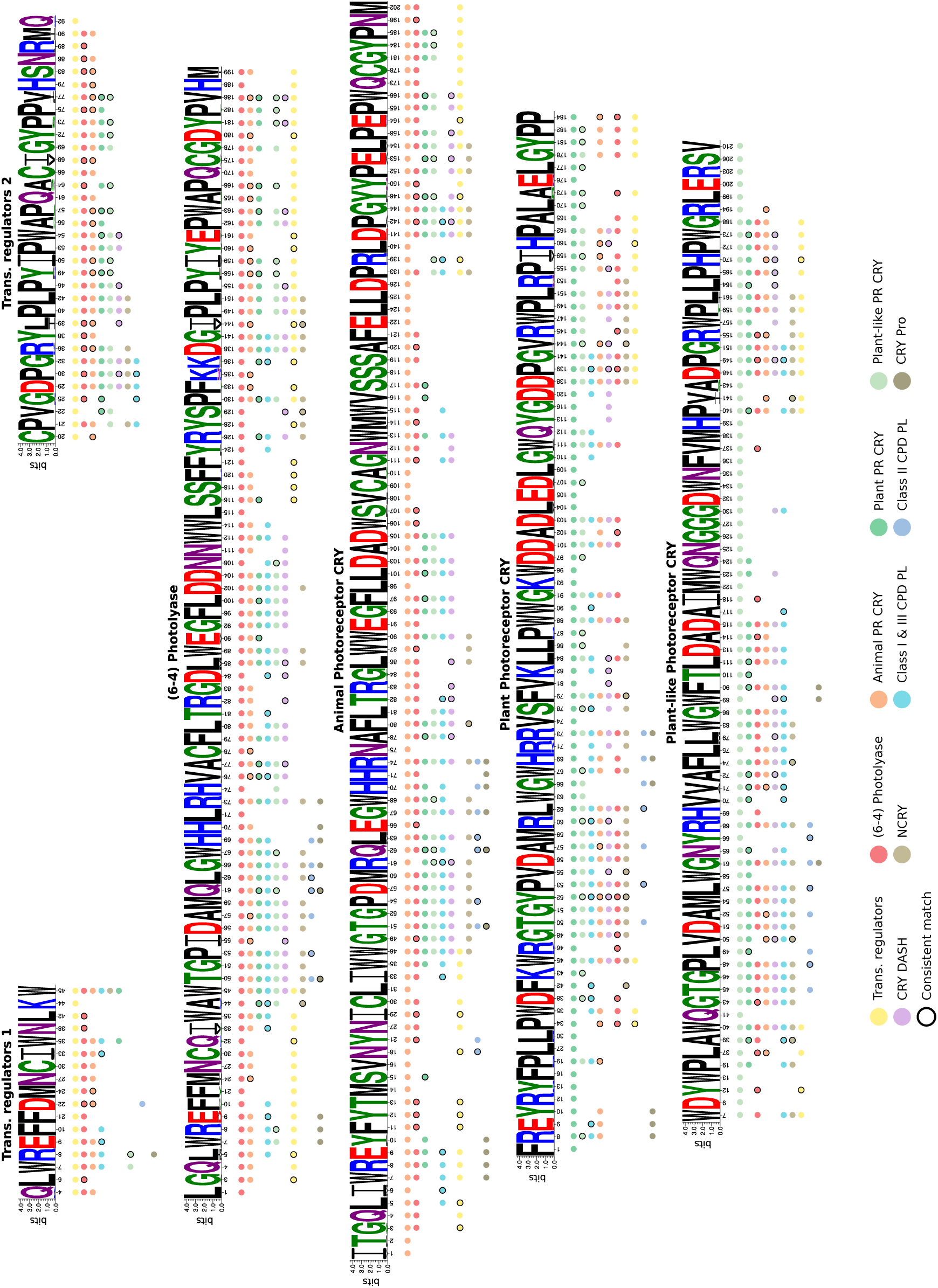
Eleven motifs for 10 subtrees in the ProfileView tree of CPF. Each motif for a class is represented by the most conserved positions in the corresponding representative model, that is positions showing *>* 60% frequency in the associated alignment (see Methods). Below each position, the coloured dots indicate that the position is well-conserved in other motifs (after their alignment; see Methods). Circled dots indicate that the position in the motif is not conserved as much in another motif (see Methods). The possible asymmetric distribution of color dots or an absence of dots between comparable positions in motifs is explained in Methods. Specific positions in a motif have no additional dot. The transcriptional regulators’ subtree, represented by two distinct representative models, is provided with two independent motifs. For each motif, coloured dots are ordered, from top to bottom, depending on the best E-values given by hhblits to the pairwise alignments.

**Figure S6:**
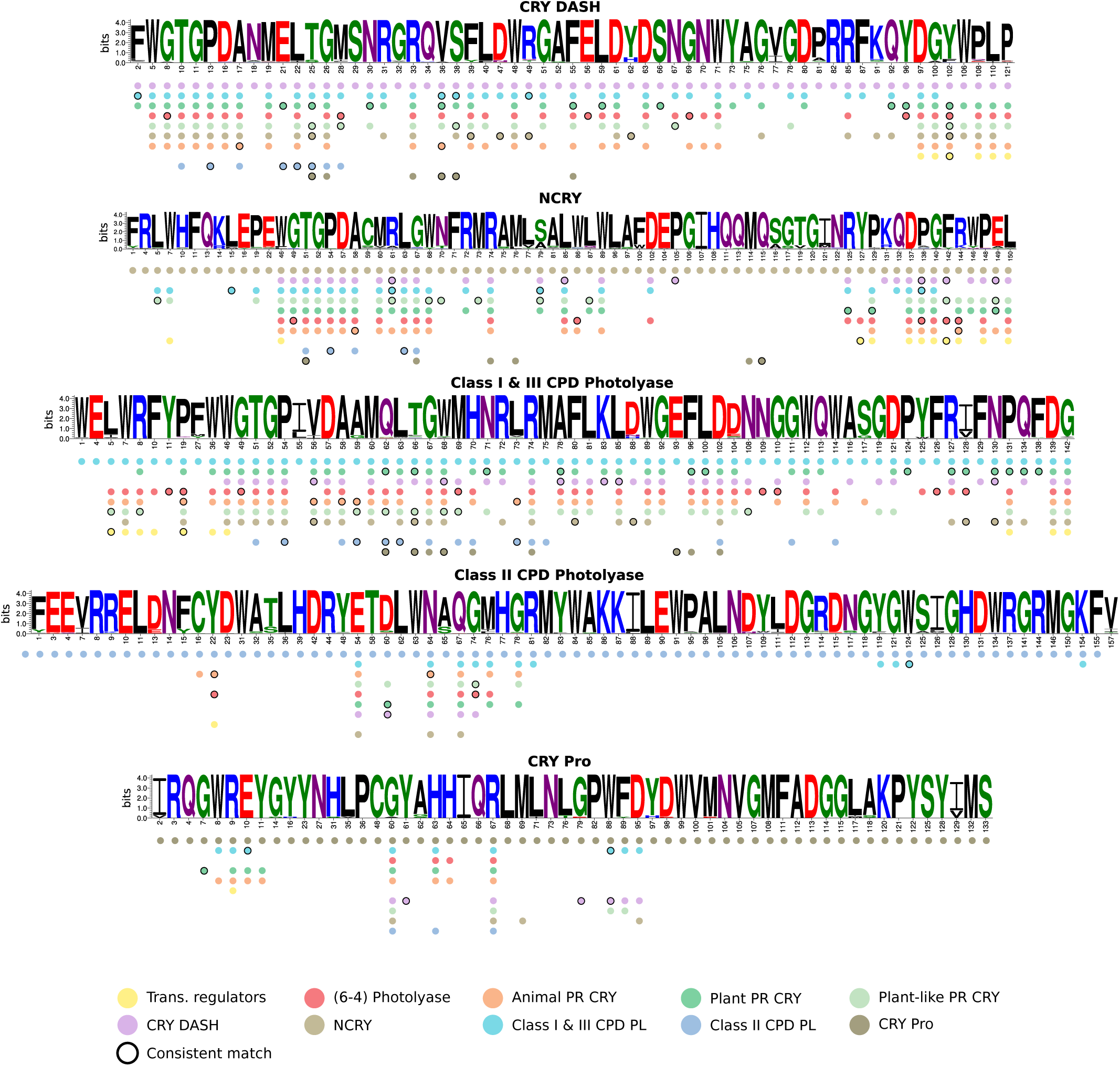
Eleven motifs for 10 subtrees in the ProfileView tree of CPF (continued). See legend in **Fig. S5**.

**Figure S7:**
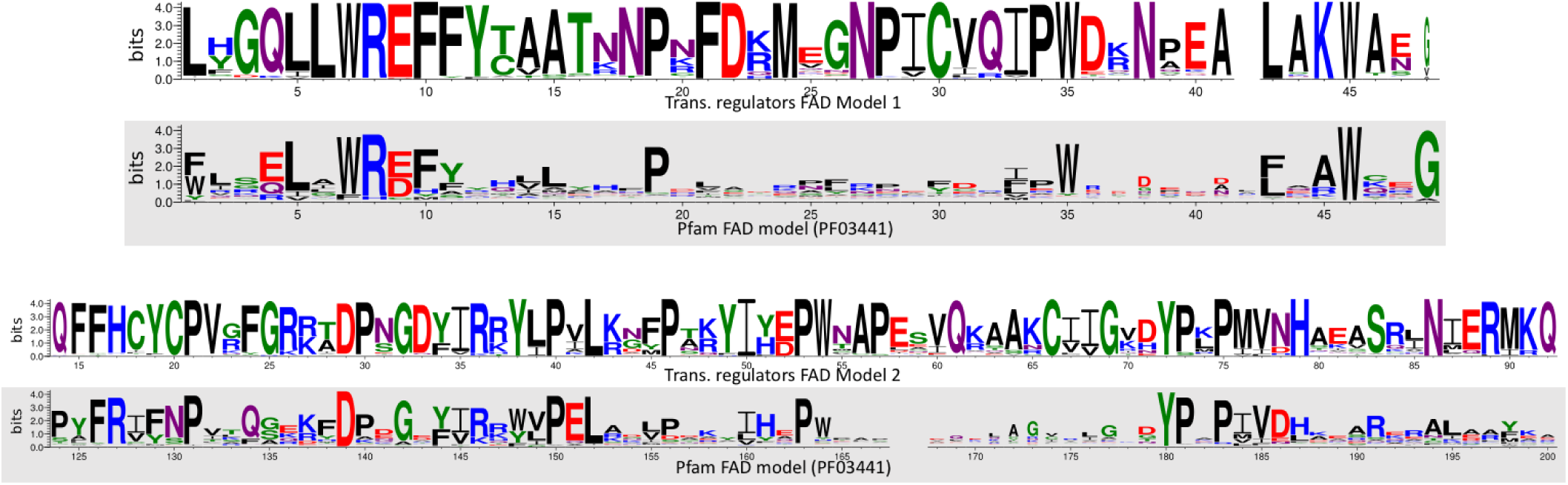
Comparison between trans. regulators models and Pfam models for the CPF family. Alignment of the two full models, corresponding to the trans. regulators motif 1 in A (top) and the trans. regulators model 2 in B (bottom), on two distinct regions of the Pfam FAD model PF03441 (grey background). The regions do not overlap.

**Figure S8:**
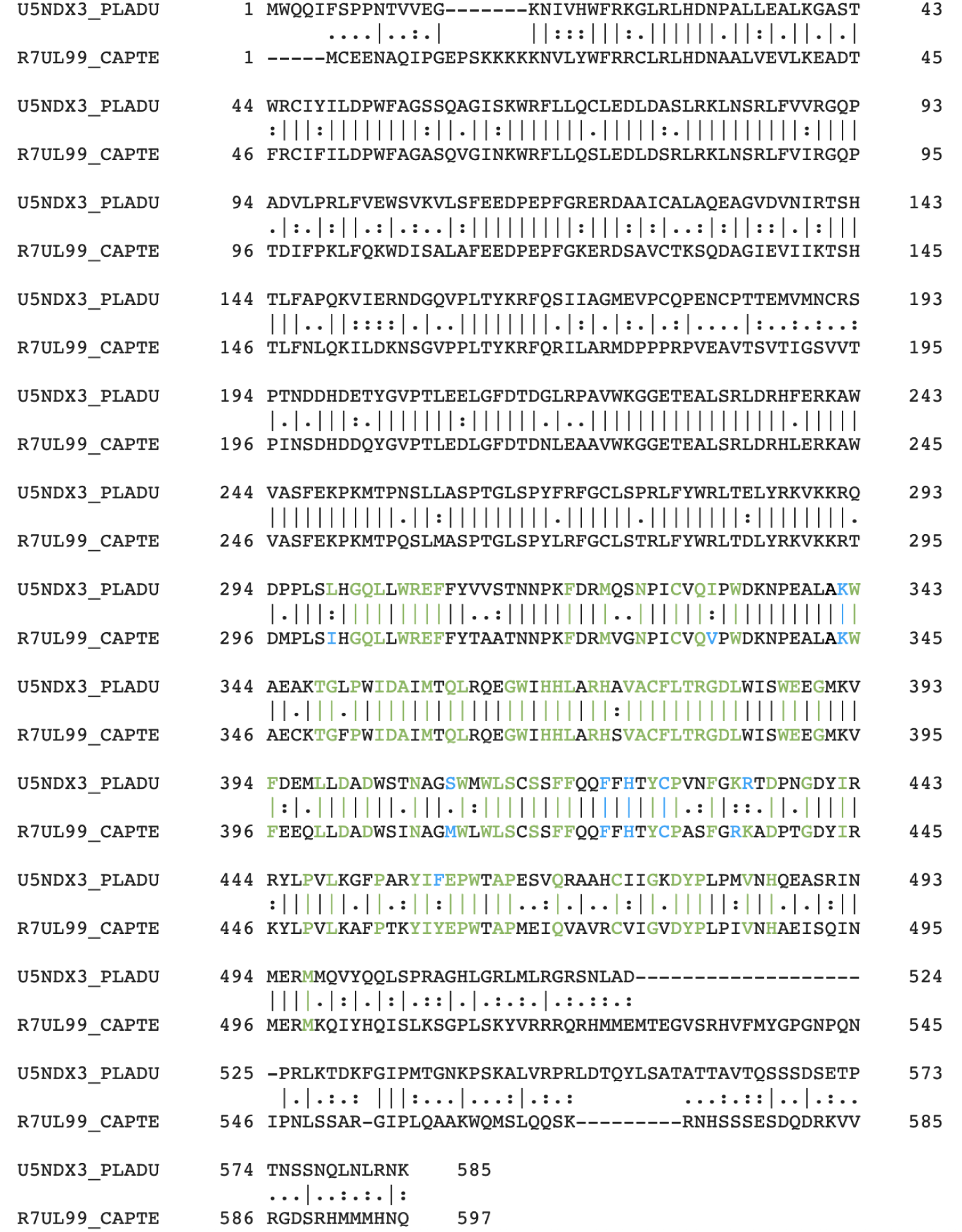
Sequence alignment for the pair of CPF sequences U5NDX3 and R7UL99. The alignment has been realised at https://www.ebi.ac.uk/Tools/psa/emboss_needle/. The sequence U5NDX3 has been classified by ProfileView as a “(6-4) PL” and the sequence R7UL99 as a “transcriptional regulator” sequence. In green: positions matched by the conserved motif, associated with the representative model of the “(6-4) PL” subgroup of CPF, and displaying the same amino acid of the motif. In blue: as in green but where the amino acid is different in the motif.

**Figure S9:**
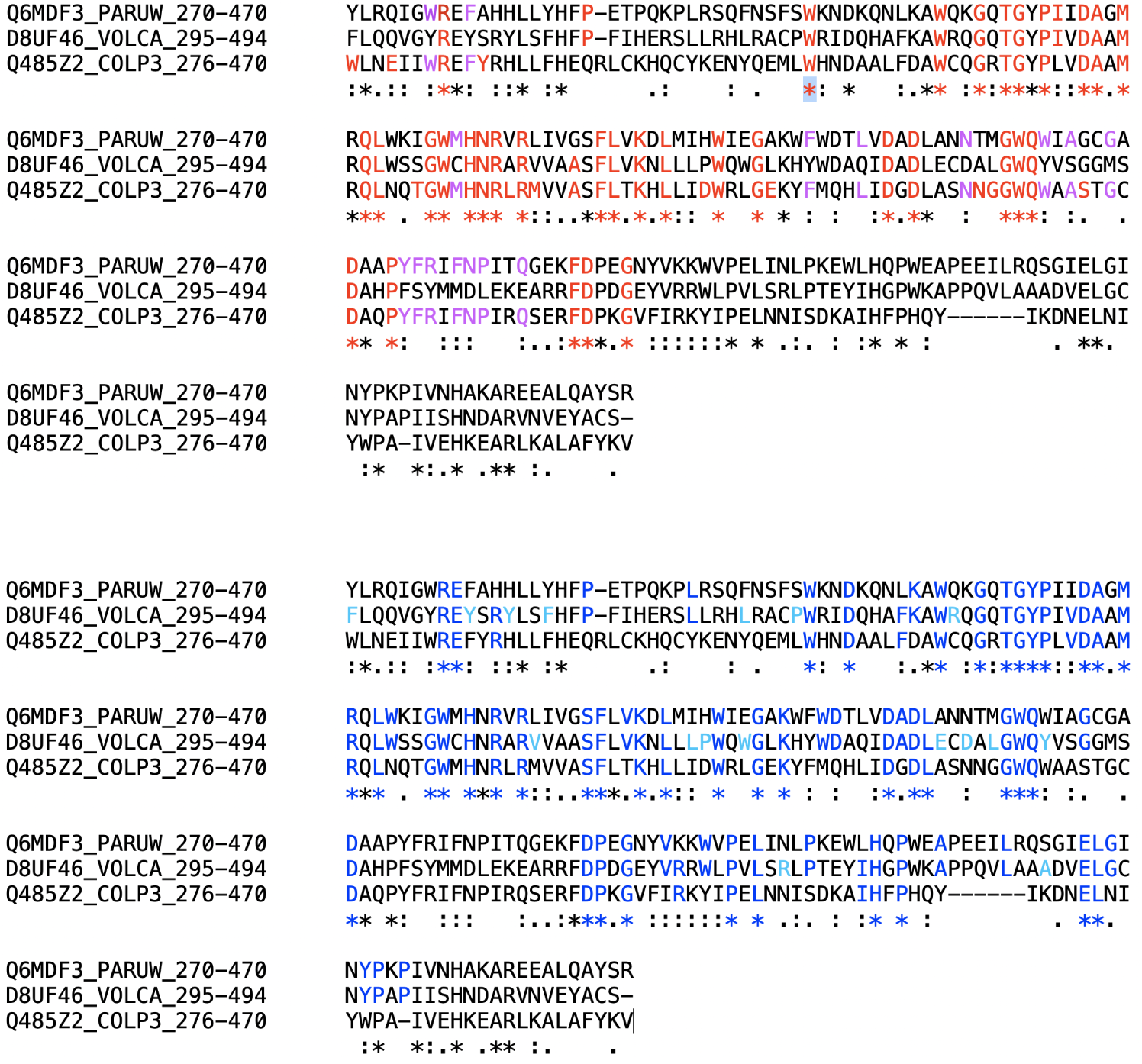
Sequence alignment for the three CPF sequences Q6MDF3, D8UF46 and Q485Z2. The alignment has been realised at https://www.genome.jp/tools-bin/clustalw. It displays 67.2% of sequence similarity (43.8% of sequence identity) between Q6MDF3 and D8UF46, and 62.6% of sequence similarity (45.6% of sequence identity) between Q6MDF3 and Q485Z2 which explain the topology of the CPF and FAD phylogenetic trees. Sequence Q6MDF3 is classified as “Class III CPD Photolyase” by ProfileView, D8UF46 as “Plant Photoreceptor CRY” and Q485Z2 as “Class I CPD Photolyase”. Top: 66 positions in the alignement are colored in red/purple because they are positions in the **representative motif for Classes I & III CPD photolyase** (**Fig. S6**). In 34 of these positions, the three sequences share the same amino acid and in 16 of them (purple), sequence D8UF46 does not share the amino acid with Q6MDF3 and Q485Z2, suggesting that Q6MDF3 and Q485Z2 are functionally closer sequences than Q6MDF3 and D8UF46, as suggested by the phylogenetic tree of CPF sequences (**Fig. S3**). Bottom: 83 positions in the alignement are colored in blue/cyan because they are positions in the **representative motif for Plant Photoreceptor CRY** (**Fig. S5**). In 43 of these positions, the three sequences share the same amino acid and in 17 of them (cyan), sequence D8UF46 does not share the amino acid with Q6MDF3 and Q485Z2, suggesting that D8UF46 is functionally distinct from Q6MDF3 and Q485Z2.

**Figure S10:**
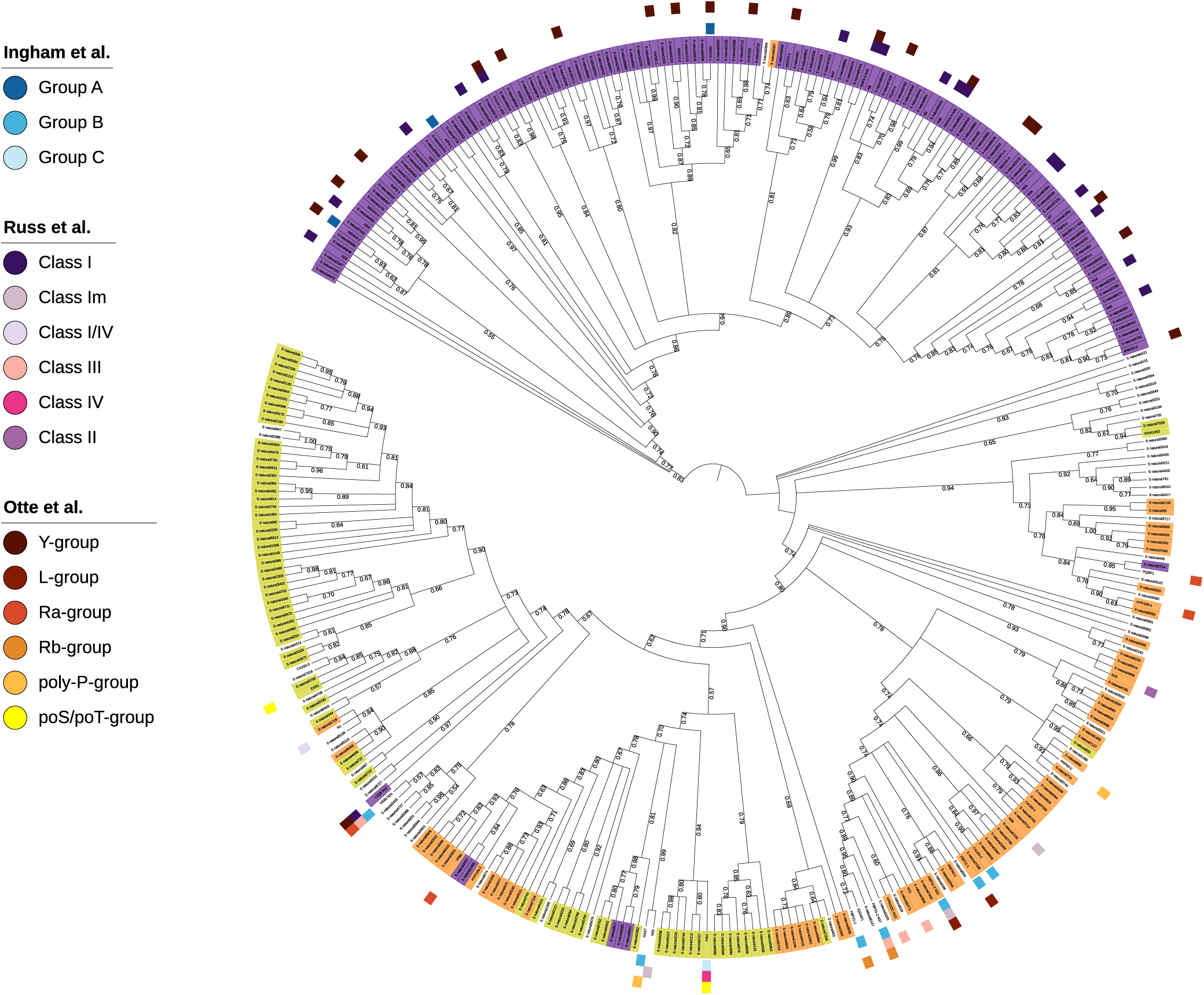
Phylogenetic tree for WW domains. The phylogenetic tree for WW domains shows that sequences classified in the same group by ProfileView appear in different subtrees, often scattered in the tree. The same holds true for those sequences known to represent the same functional class for Ingham, Russ and Otte’s classifications.

**Figure S11:**
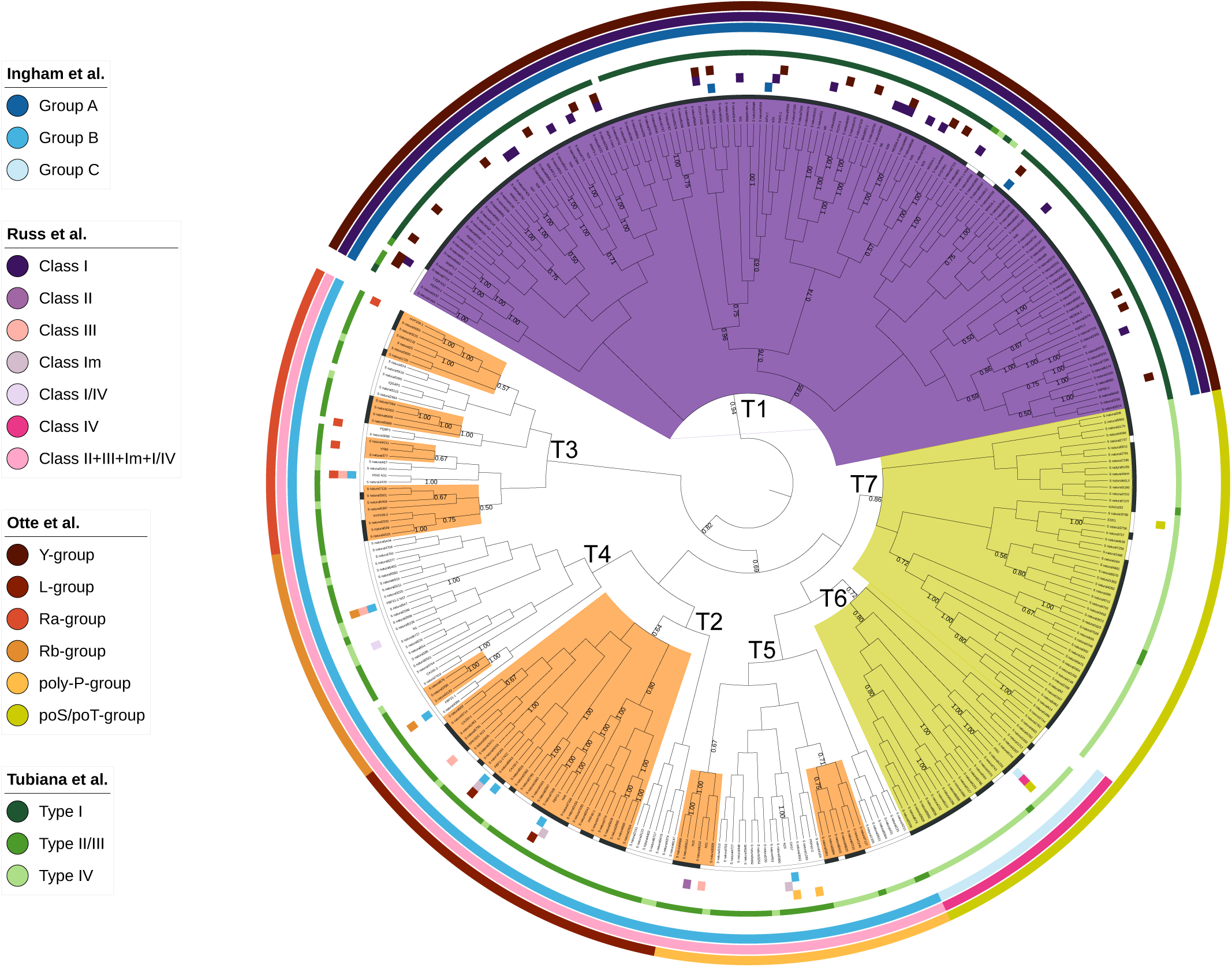
ProfileView tree for WW domains; compatibility with experimental and computational classification. Larger size of **Fig. 5D**.

**Figure S12:**
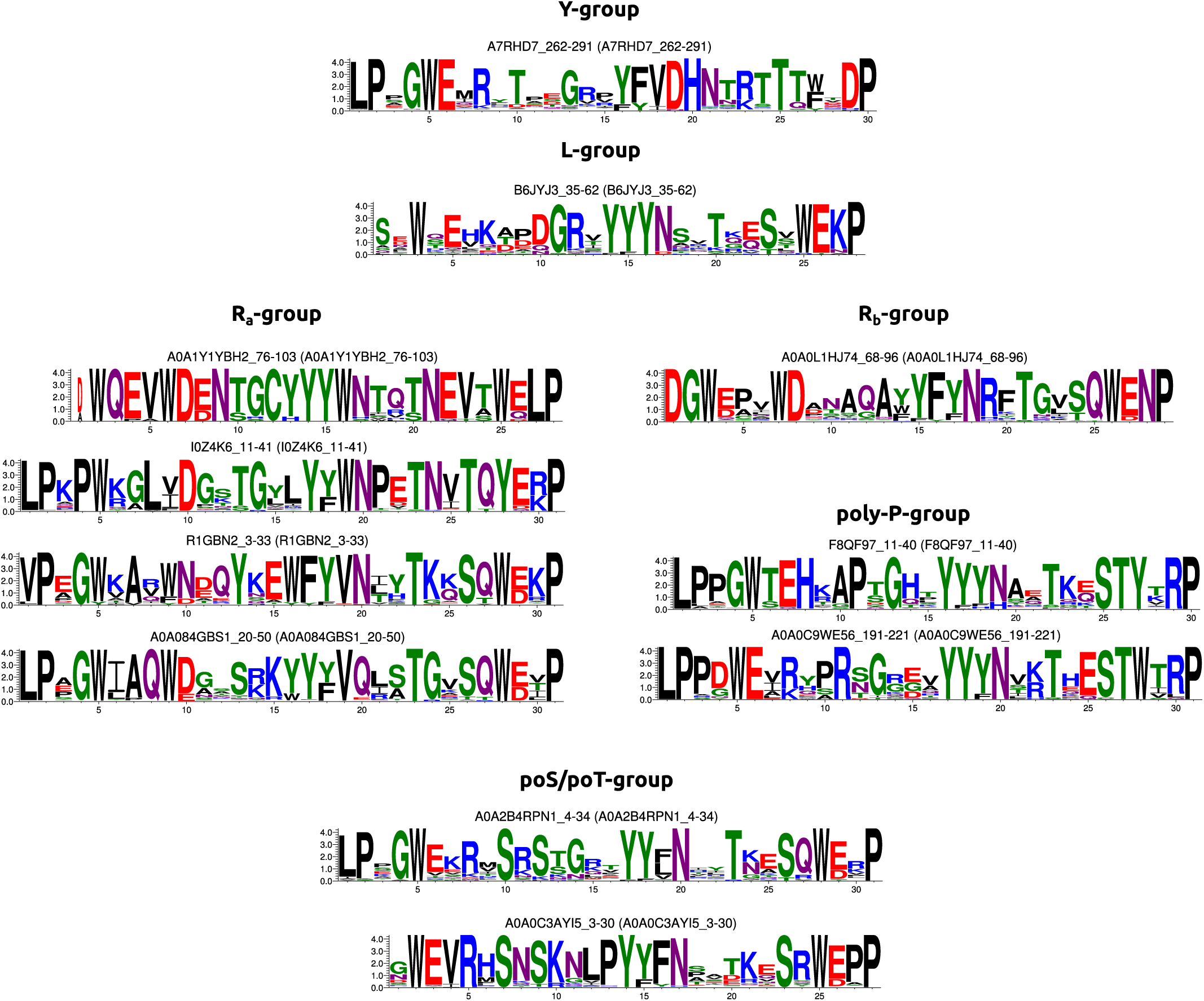
Representative models in ProfileView tree of WW domains. Models are representative of the sequences organised in the colored subtrees of **Fig. 5**. For Otte’s classes containing more than one representative model, the order, from top to bottom, corresponds to subtrees read anticlockwise in the outer circle (brown scale) of **Fig. 5**, corresponding to Otte *et al* classification.

**Figure S13:**
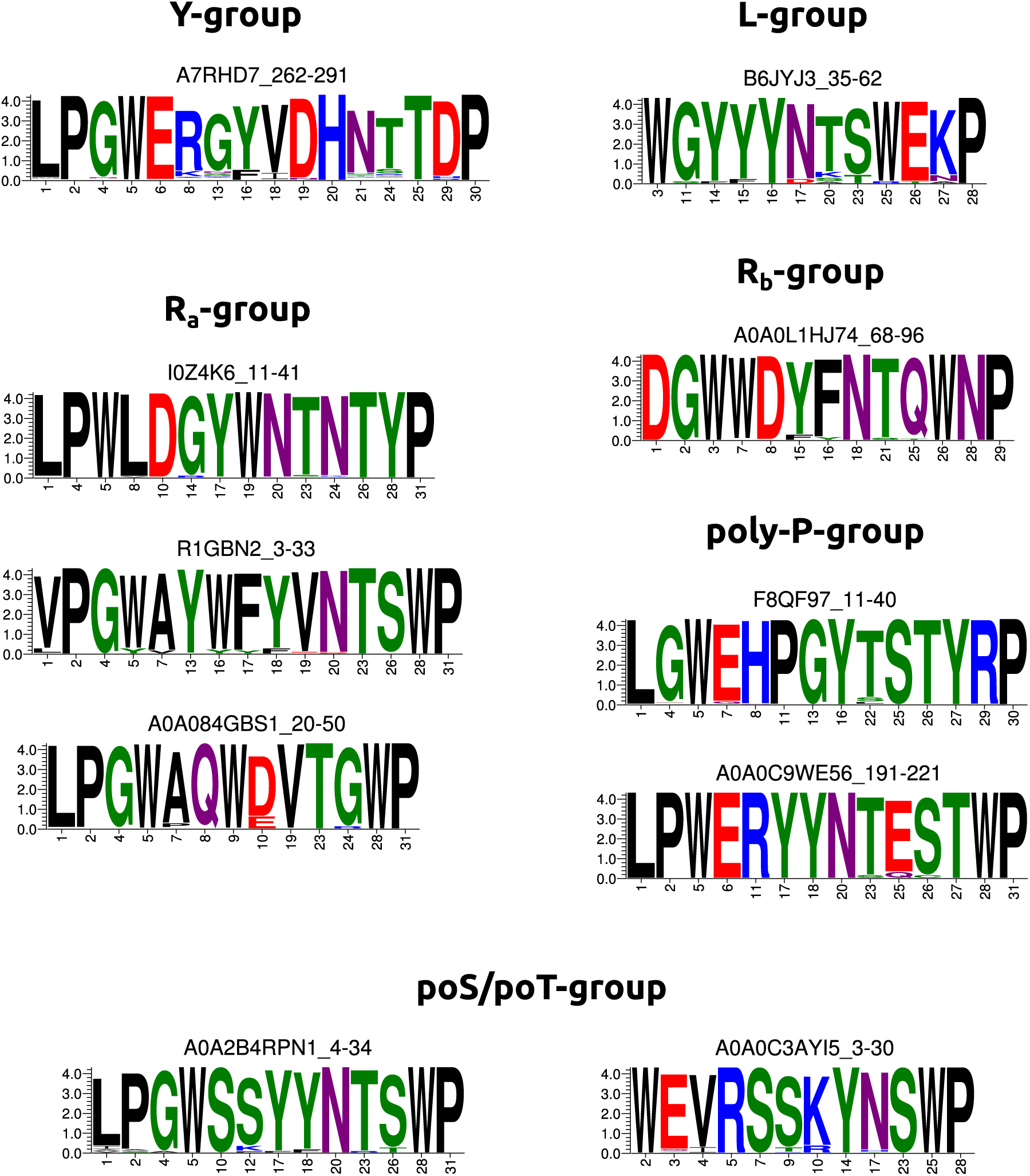
Ten motifs for 11 subtrees in the ProfileView tree of WW domains, based on hhblits conservation criteria. Each motif for a group is represented by the most conserved positions in the corresponding representative model, that is positions showing *>* 60% frequency for hhblits in the associated alignment (see Methods). Notice that by using the hhblits criteria, one of the R_a_ models does not provide any conserved motif (this is due to the very low number of sequences in the alignment generating the model, 20, and to the length of the model). Compare to **Figure S14**.

**Figure S14:**
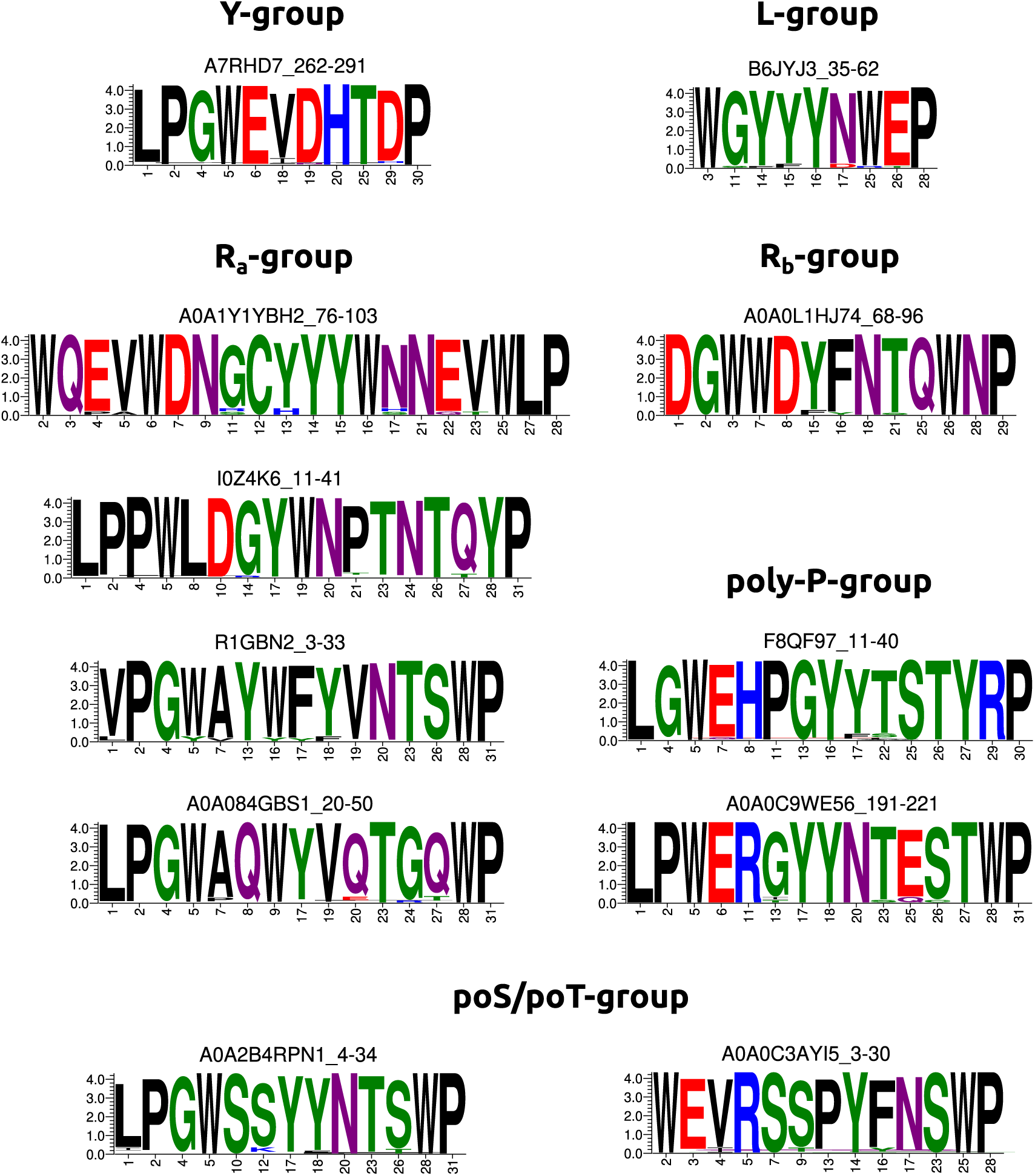
Eleven motifs for 11 subtrees in the ProfileView tree of WW domains, based on amino acid counting. Each motif for a group is represented by the most conserved positions in the corresponding representative model, that is positions showing *>* 90% amino acid frequency (excluding gaps) in the associated alignment (see Methods). Note that this conservation criteria recovers 4 motifs for the R_a_ group. Compare to **Figure S13**.

**Figure S15:**
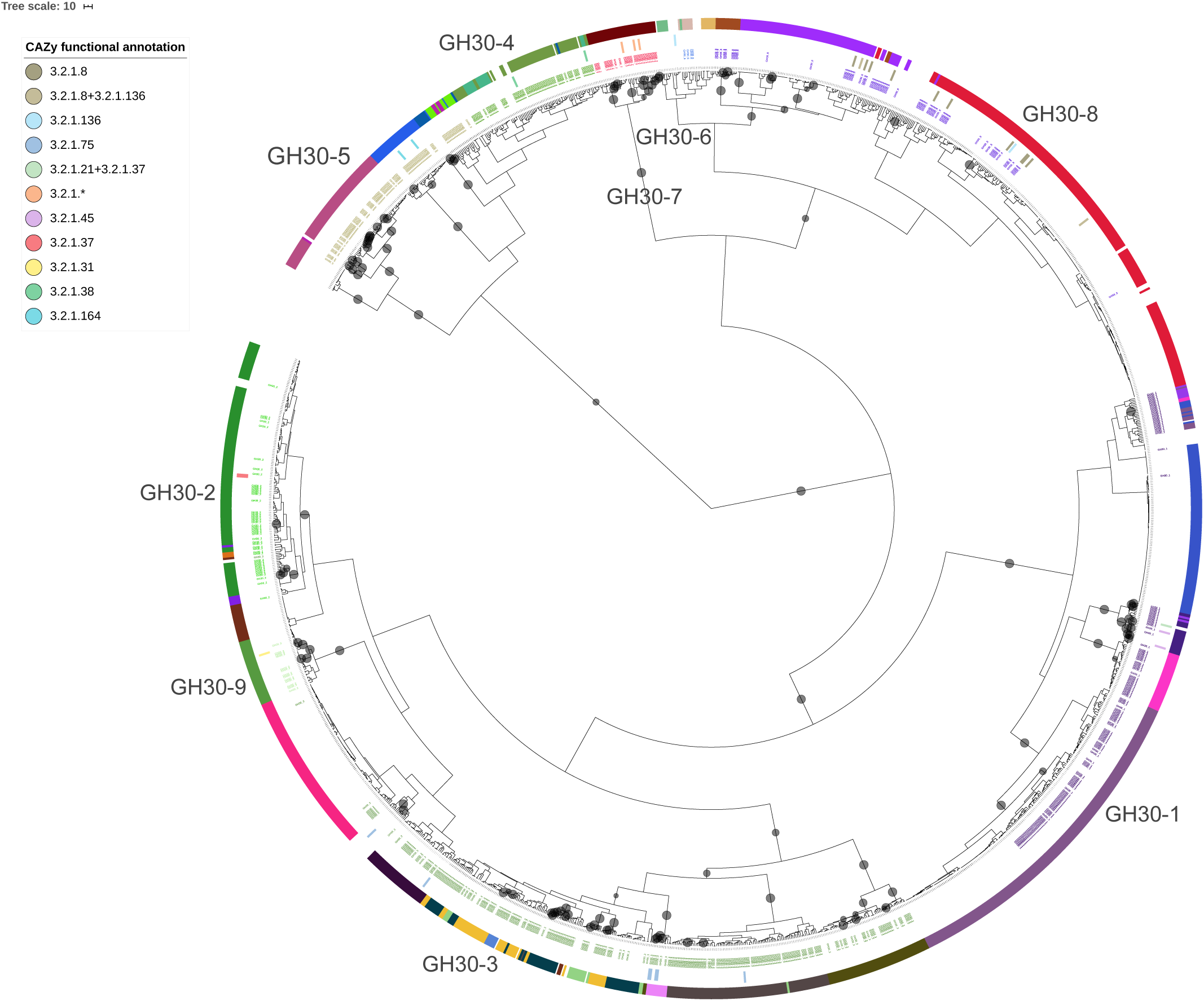
ProfileView tree of GH30 sequences. The tree is based on the construction of models for the two pfam domains PF02055 (Glyco hydro 30) and PF14587 (Glyco hydr 30 2). Black dots in the tree indicate the existence of representative models separating at least 75% of the sequences in the subtree (note that lowering the threshold to 50% provides comparable results). The first external ring contains the labels of CAZy subfamilies (GH30 1,…, GH30 9), also indicated in larger characters on the annotated tree for an easier reading. Sequences and their classification correspond to those used in Figure 3 of (Barrett and Lange, 2019). The second ring reports the existence of a “EC number” providing the functional annotation in CAZy. The EC numbers and their associated colours are indicated on the top left (GH30-1: 3.2.1.45 and 3.2.1.21+3.2.1.37; GH30-2: 3.2.1.37; GH30-3: 3.2.1.75; GH30-4: 3.2.1.38; GH30-5 3.2.1.164; GH30-6: –; GH30-7: 3.2.1.*; GH30-8: 3.2.1.8, 3.2.1.136, 3.2.1.8+3.2.1.136; GH30-9: 3.2.1.31). The third and most external ring reports CUPP clustering (Barrett and Lange, 2019). Different colours are used to indicate different CUPP clusters. See Table S5.

**Figure S16:**
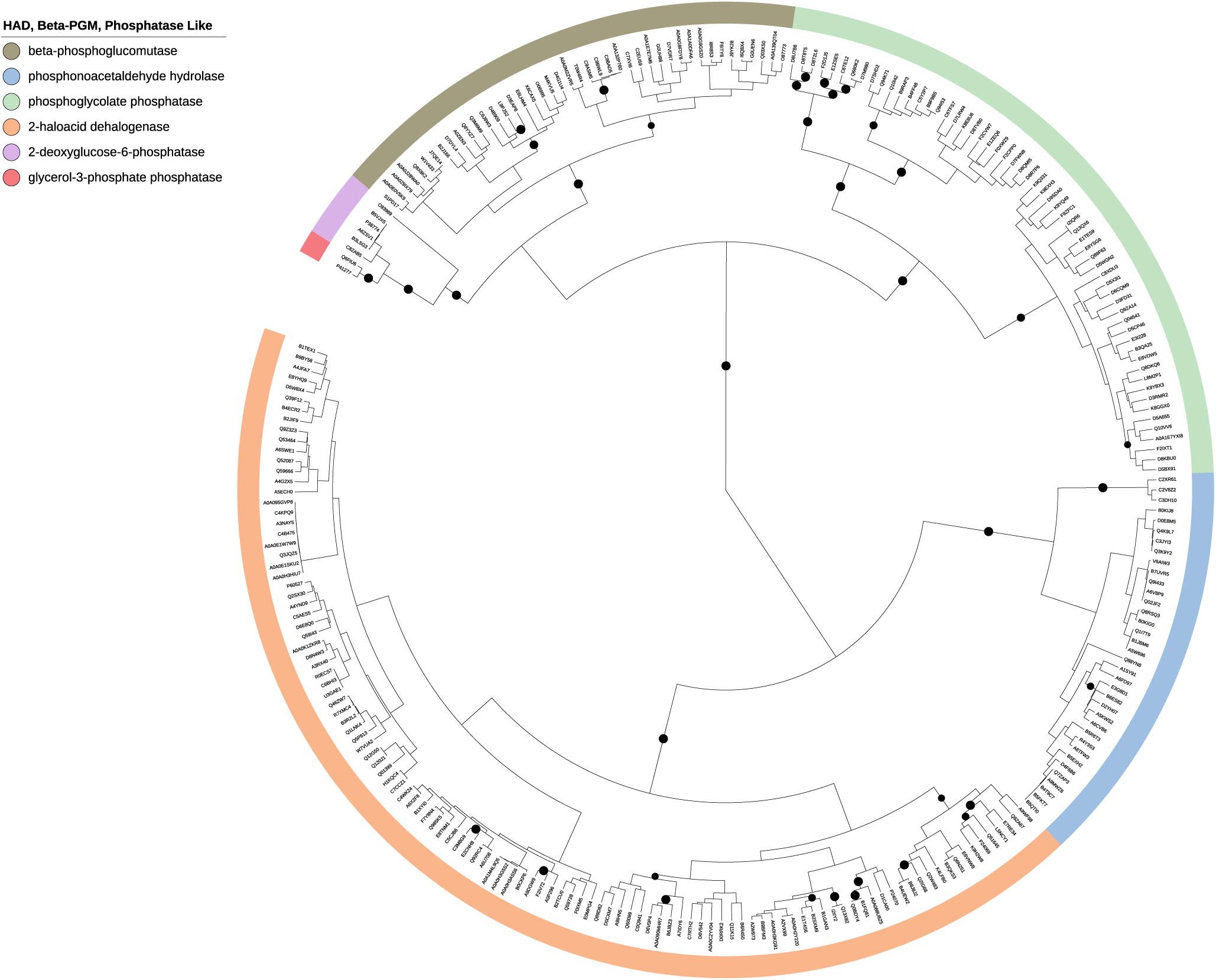
ProfileView classification tree of the HAD/*β*-PGM/Phosphatase-like subgroup of Haloacid Dehydrogenase in SFLD. Validation test of ProfileView performance. See Table S7.

**Figure S17:**
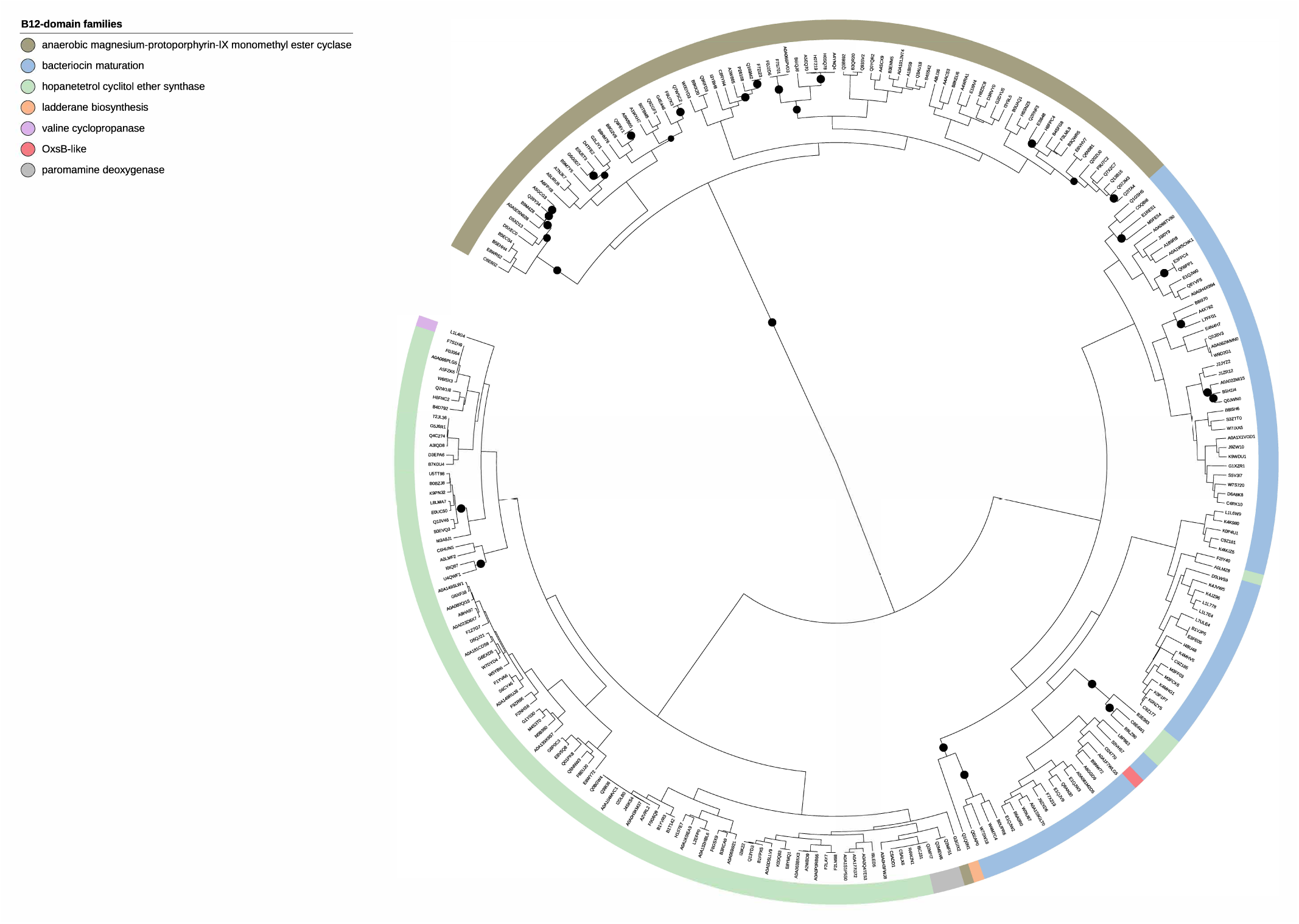
ProfileView classification tree of the B12-binding domain containing subgroup of Radical SAM in SFLD. Validation test of ProfileView performance. See Table S8.

**Figure S18:**
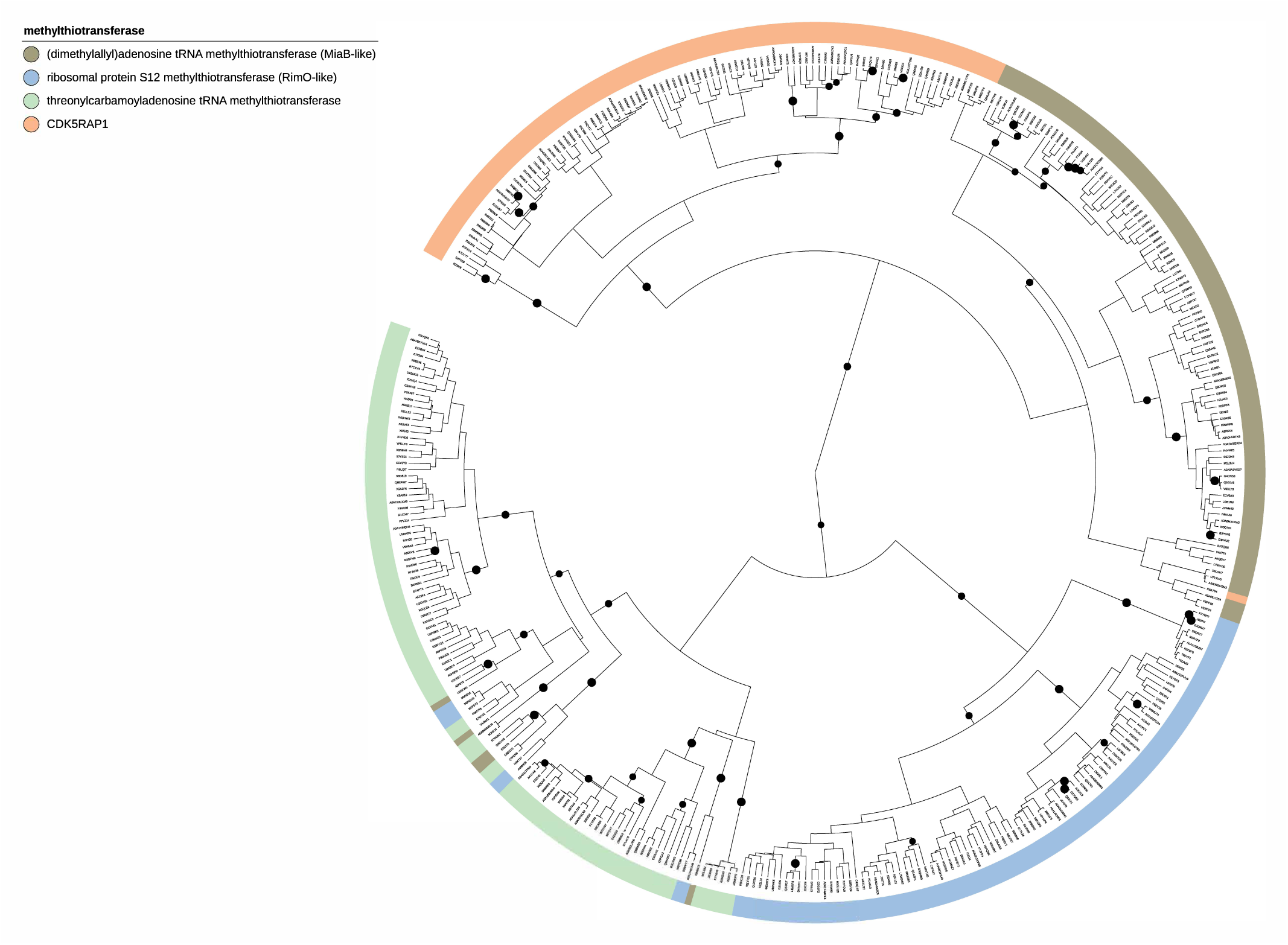
ProfileView classification tree of the Methylthiotransferase subgroup of Radical SAM in SFLD. Validation test of ProfileView performance. See Table S9.

**Figure S19:**
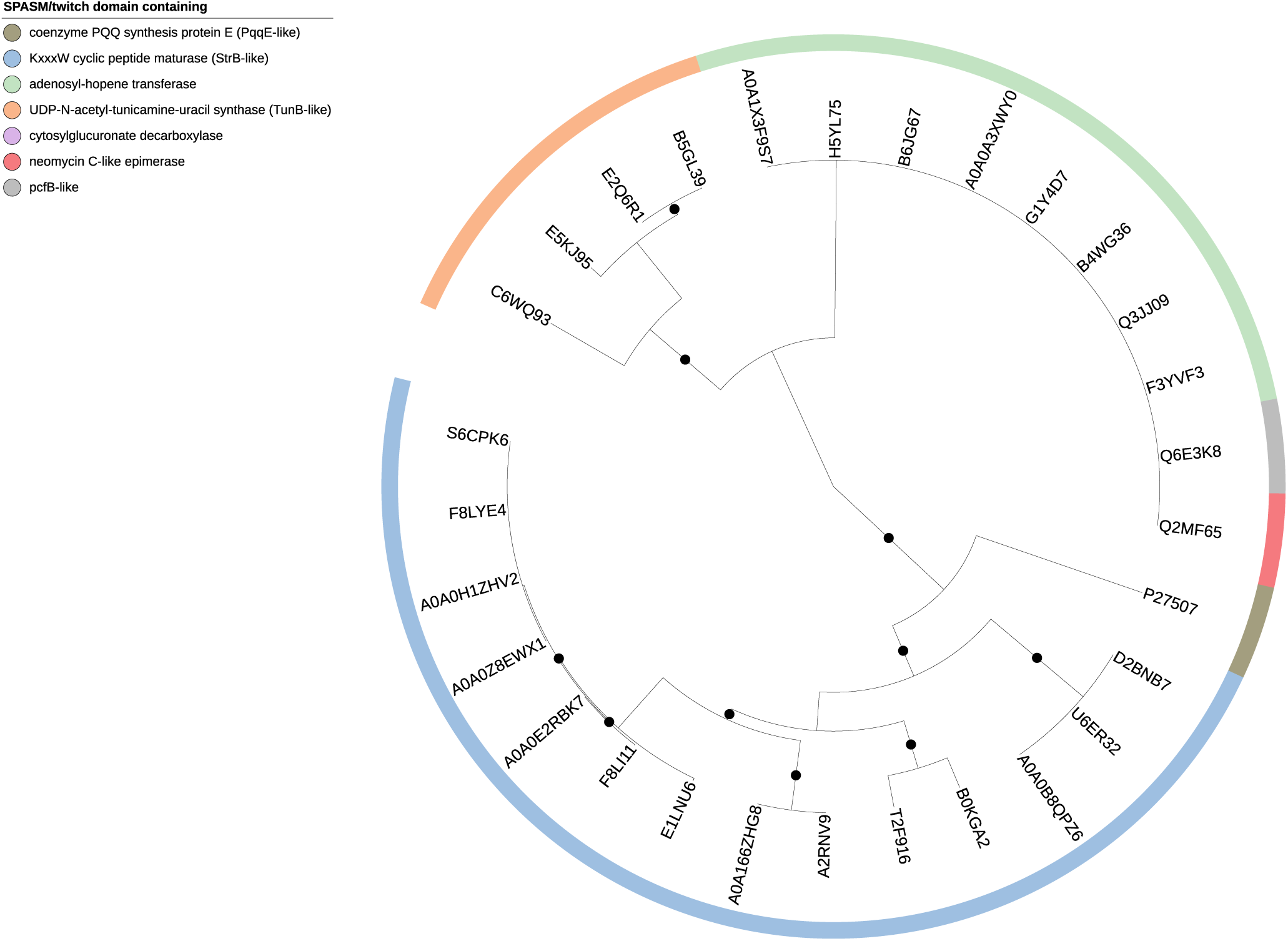
ProfileView classification tree of the SPASM/twitch domain containing subgroup of Radical SAM in SFLD based on the SPASM domain. Validation test of ProfileView performance. See Table S10.

**Figure S20:**
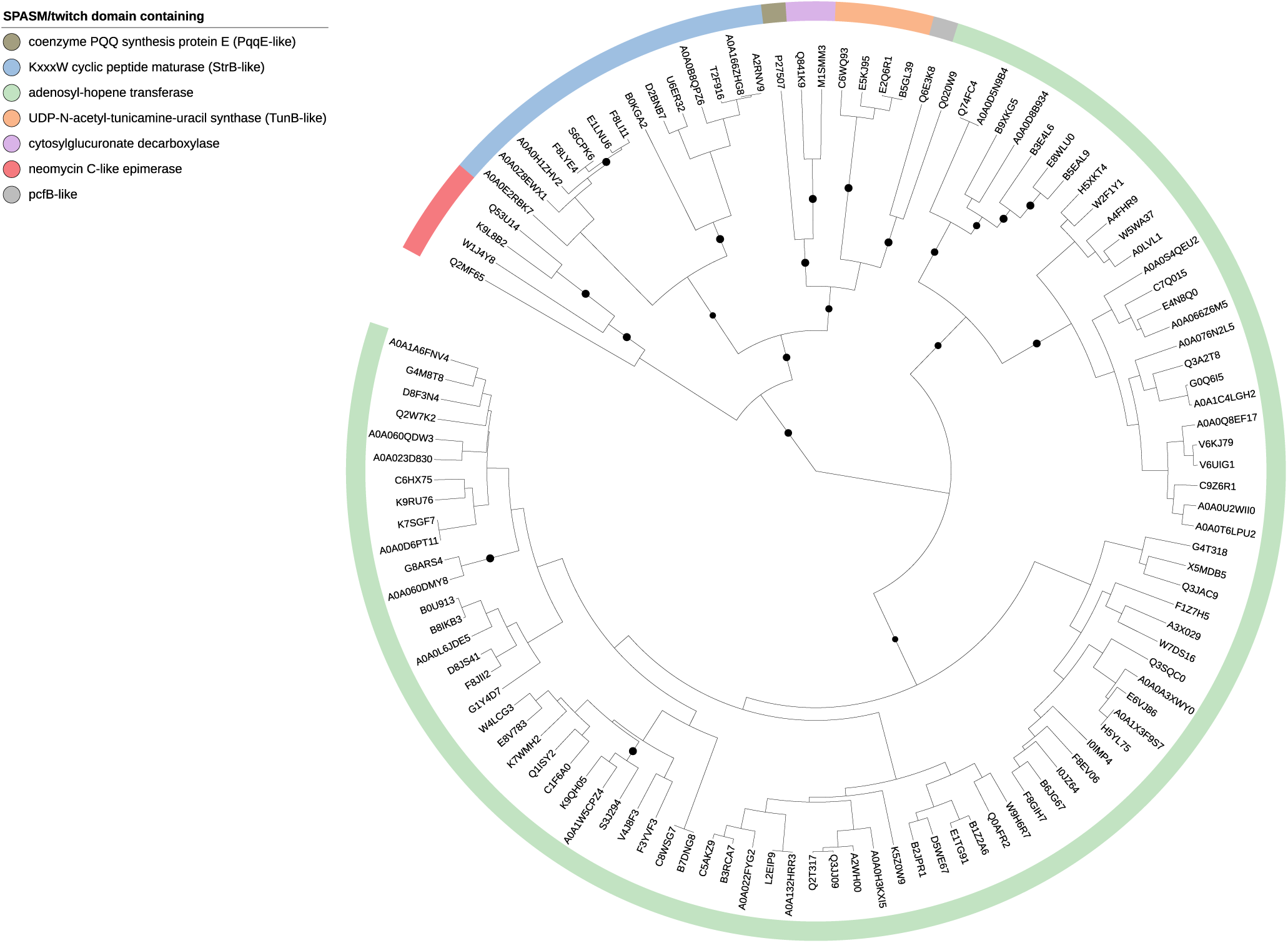
ProfileView classification tree of the SPASM/twitch domain containing of Radical SAM in SFLD based on the Radical SAM domain. Validation test of ProfileView performance. See Table S11.

**Figure S21:**
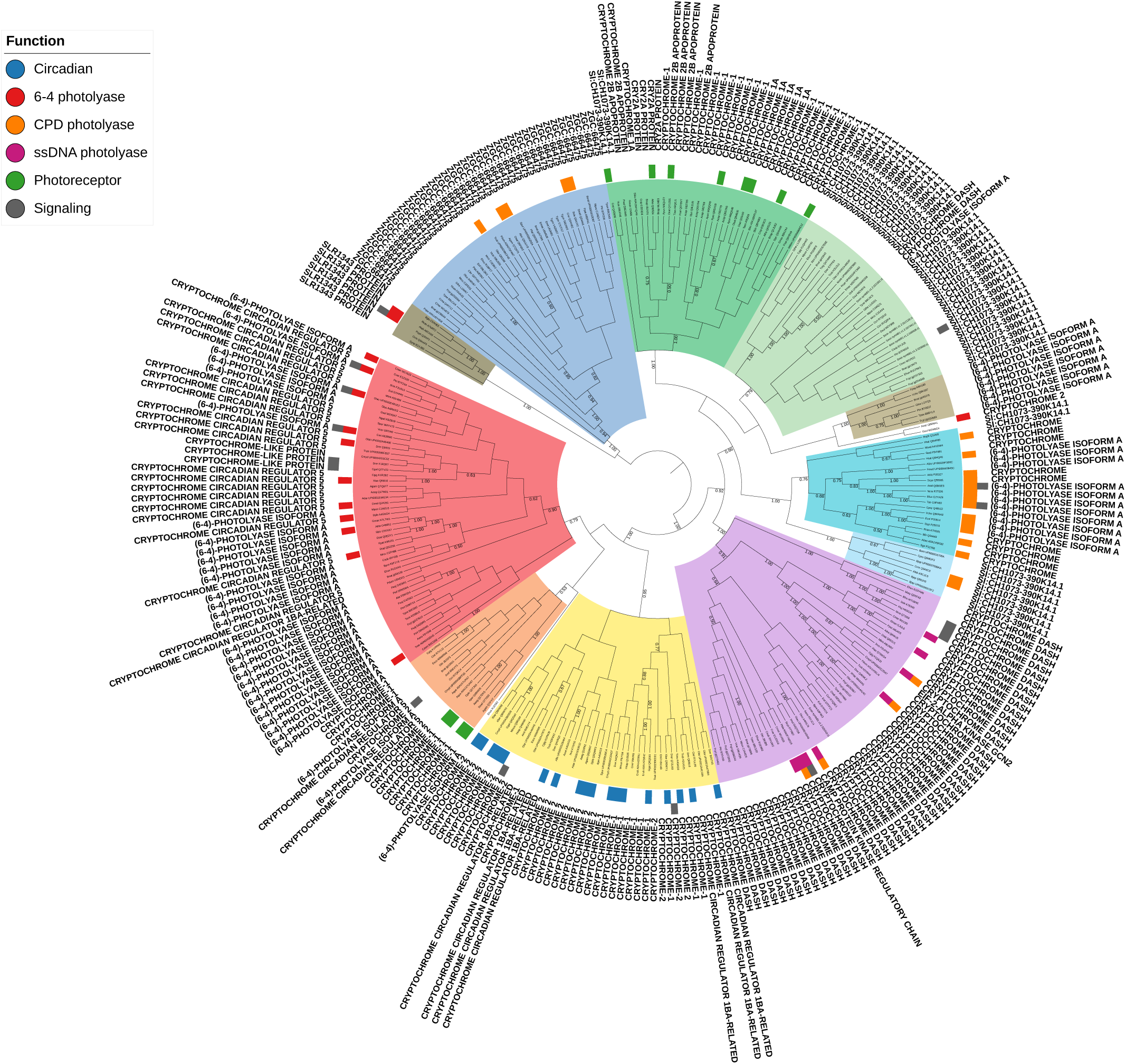
ProfileView classification tree of CPF sequences and PANTHER classification. The PANTHER classification of CPF sequences is plot on the external ring of the CPF ProfileView classification tree (**Fig. S1**).

**Figure S22:**
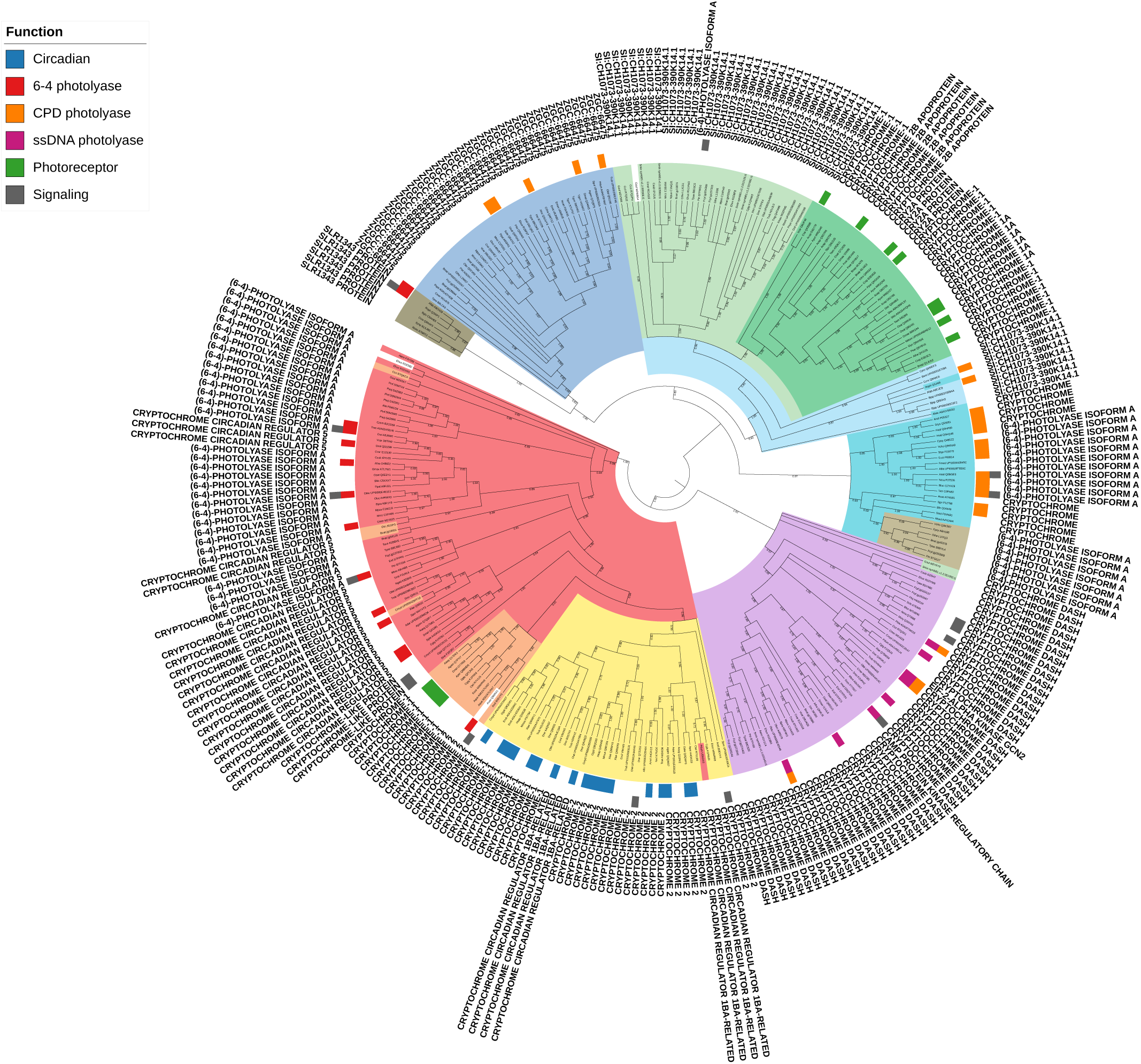
Phylogenetic tree of CPF sequences and PANTHER classification. The PANTHER classification of CPF sequences is plot on the external ring of the CPF phylogenetic tree (**Fig. S3**).

**Figure S23:**
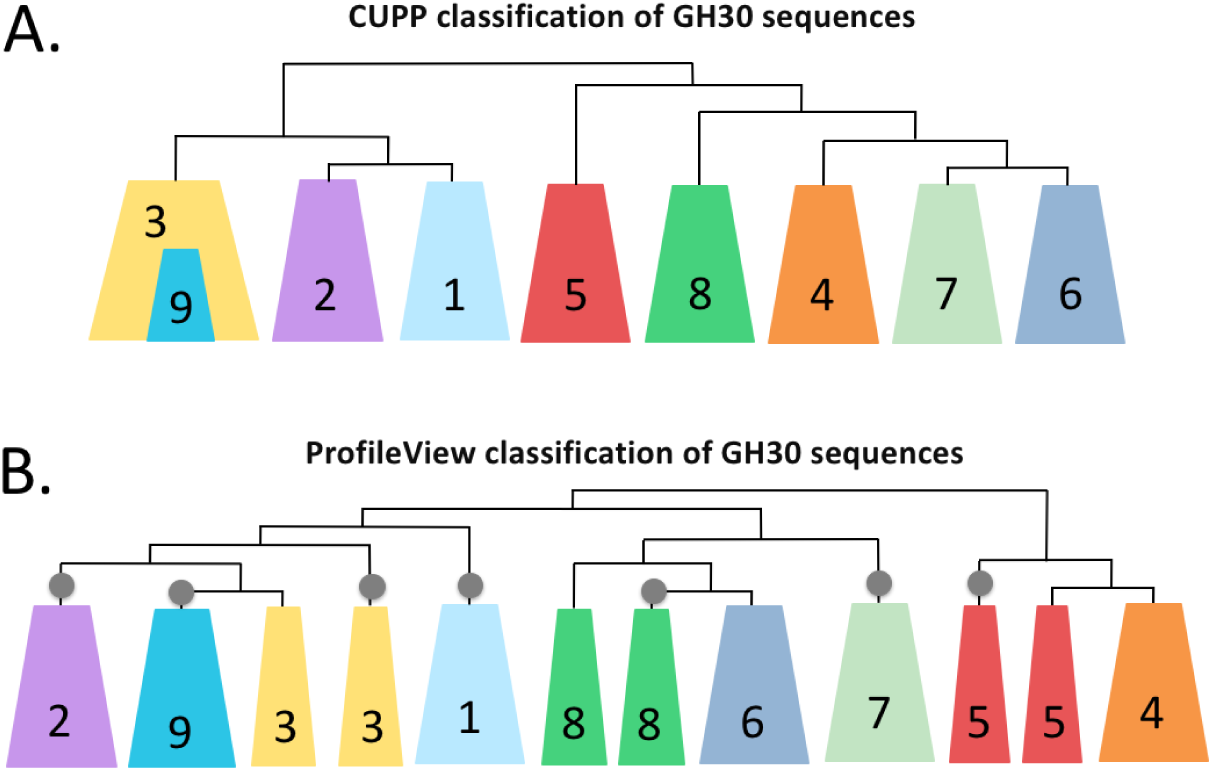
Schemas of the CUPP and ProfileView trees on GH30 sequences. A. Topology of the CUPP tree reported; reproduced from Figure 2 in (Barrett and Lange, 2019). B. Topology of the ProfileView tree. Colors and numbers correspond to GH30 subfamilies.

**Figure S24:**
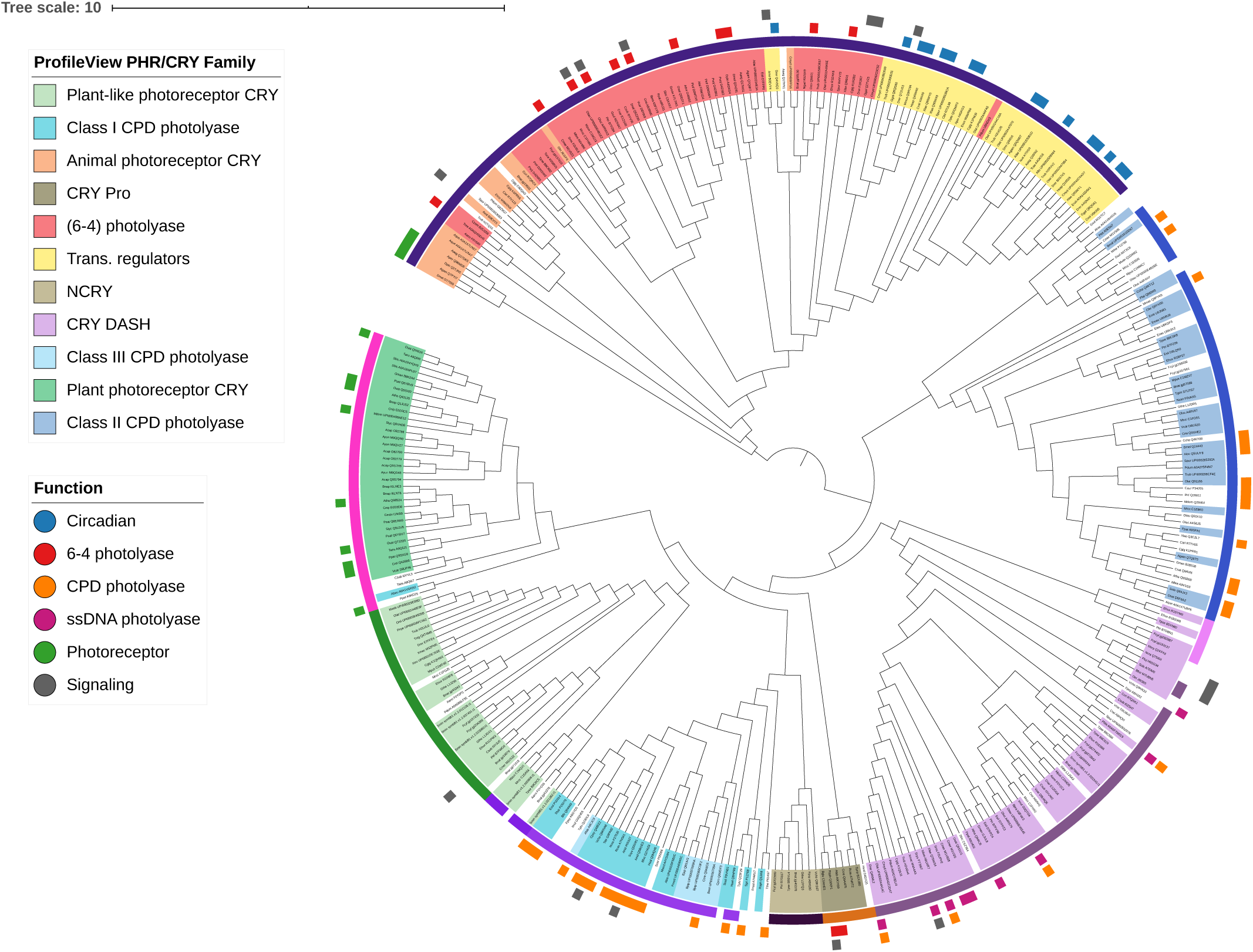
CUPP tree of CPF sequences, constructed with FAD sequences. Sequence names in the tree are coloured with ProfileView classification (with the same colour assignment of **Fig. S1**). CUPP clusters are represented by the first layer of colours around the tree (in clockwise order: dark purple, blue, pink, light purple, brown, black, violet, green, fuchsia). The two most external layers correspond to the experimental classification coming from the literature. The same information was used for ProfileView performance analysis. Compare to **Fig. S1**.

**Figure S25:**
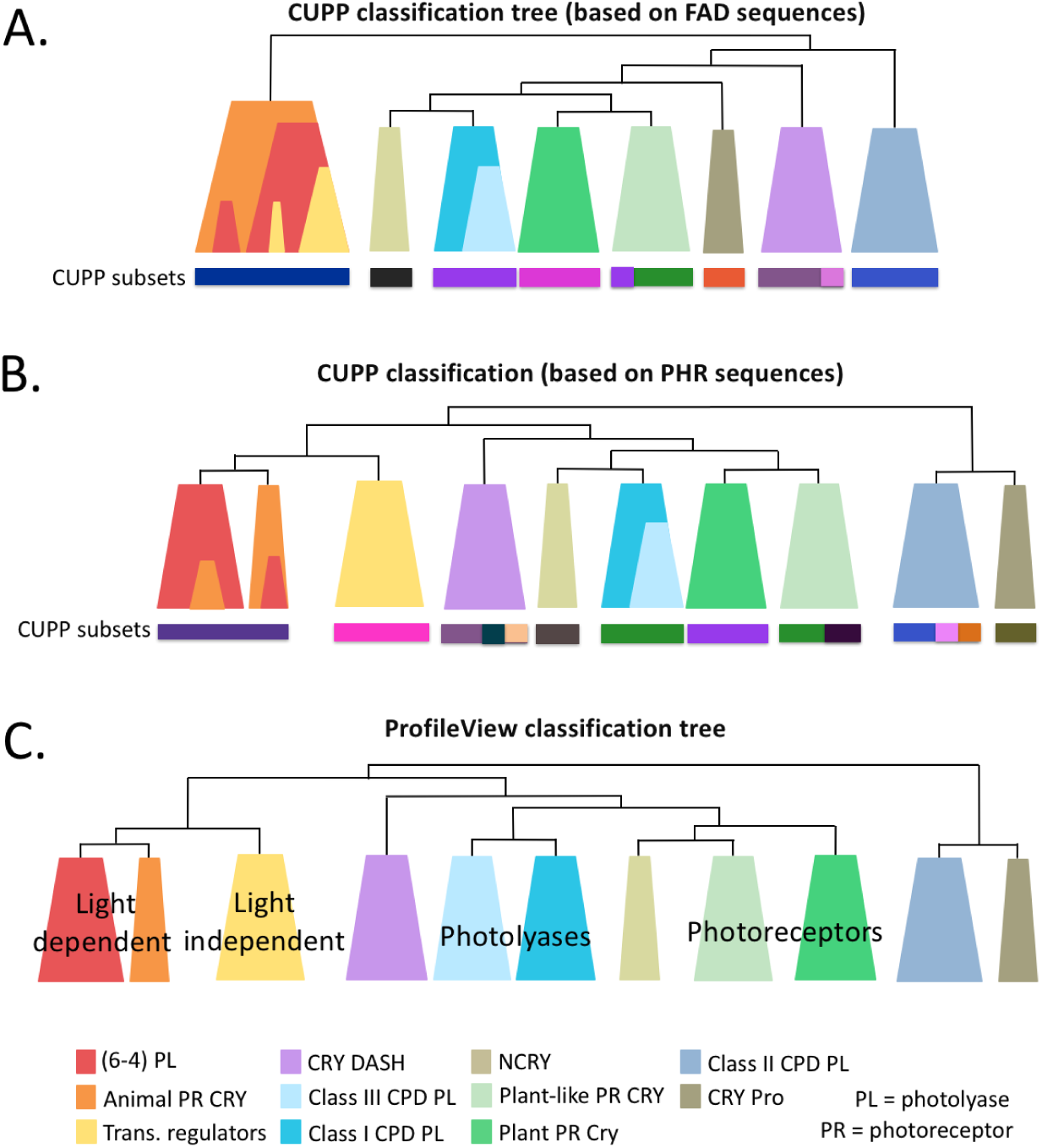
Schemas of the CUPP and ProfileView trees on CPF sequences. A. Topology of the CUPP tree constructed on FAD sequences. B. Topology of the CUPP tree constructed on PHR sequences. C. Topology of the ProfileView tree constructed on FAD sequences; taken from **Fig. 3** for an easy comparison.

**Figure S26:**
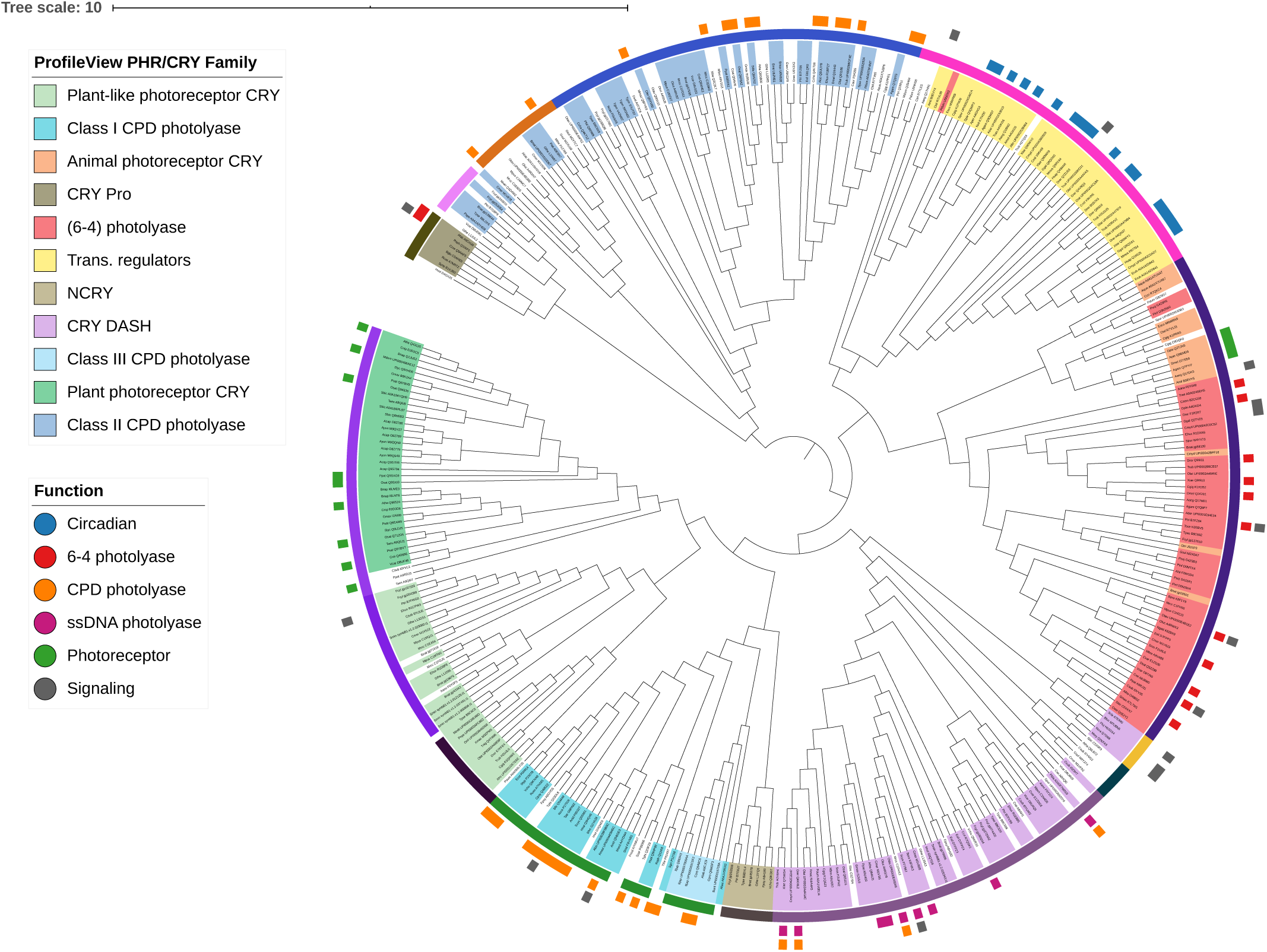
CUPP tree of CPF sequences, constructed with PHR sequences. See legend of **Figure S24**.

**Figure S27:**
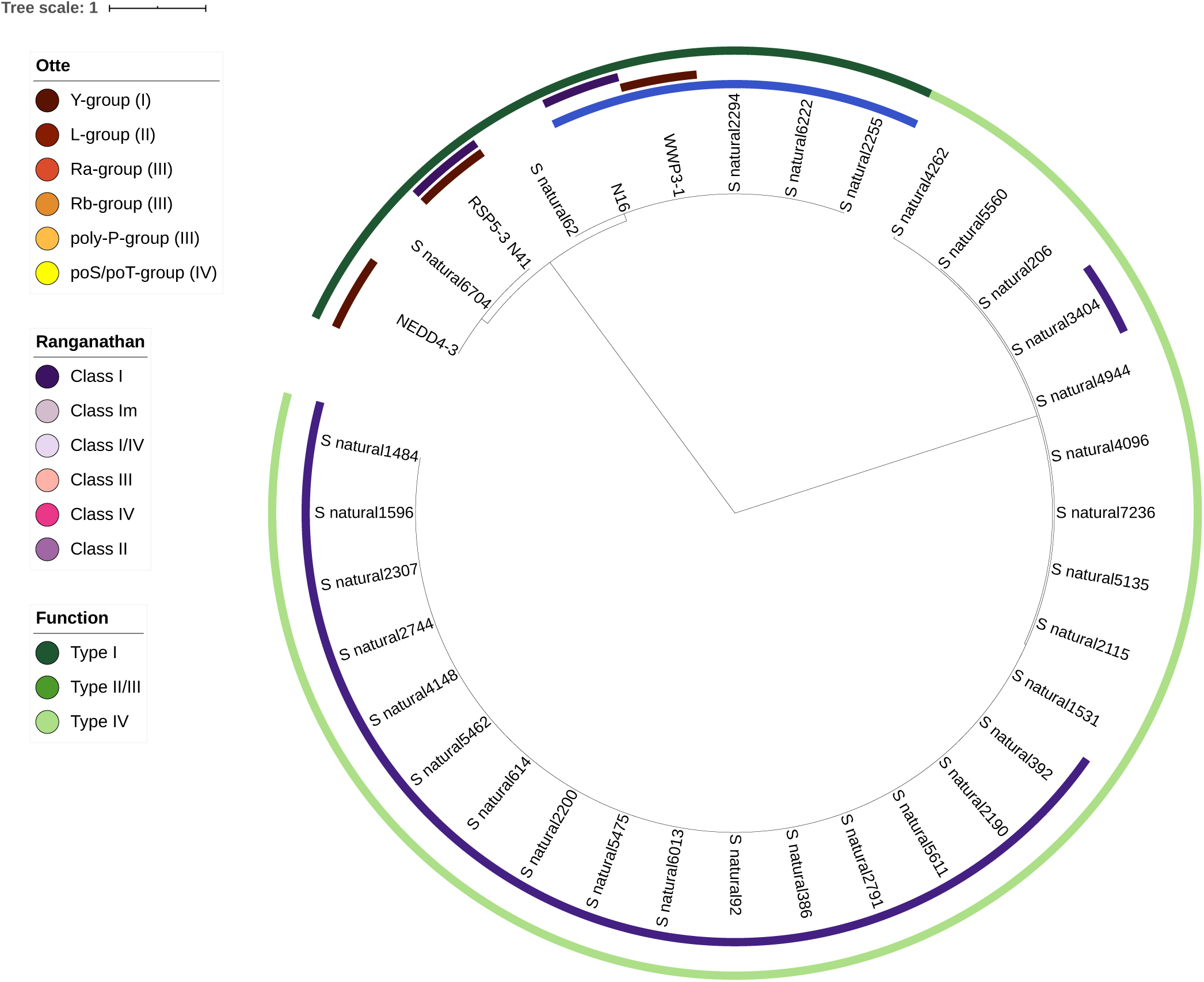
CUPP tree of WW domain sequences.

**Figure S28:**
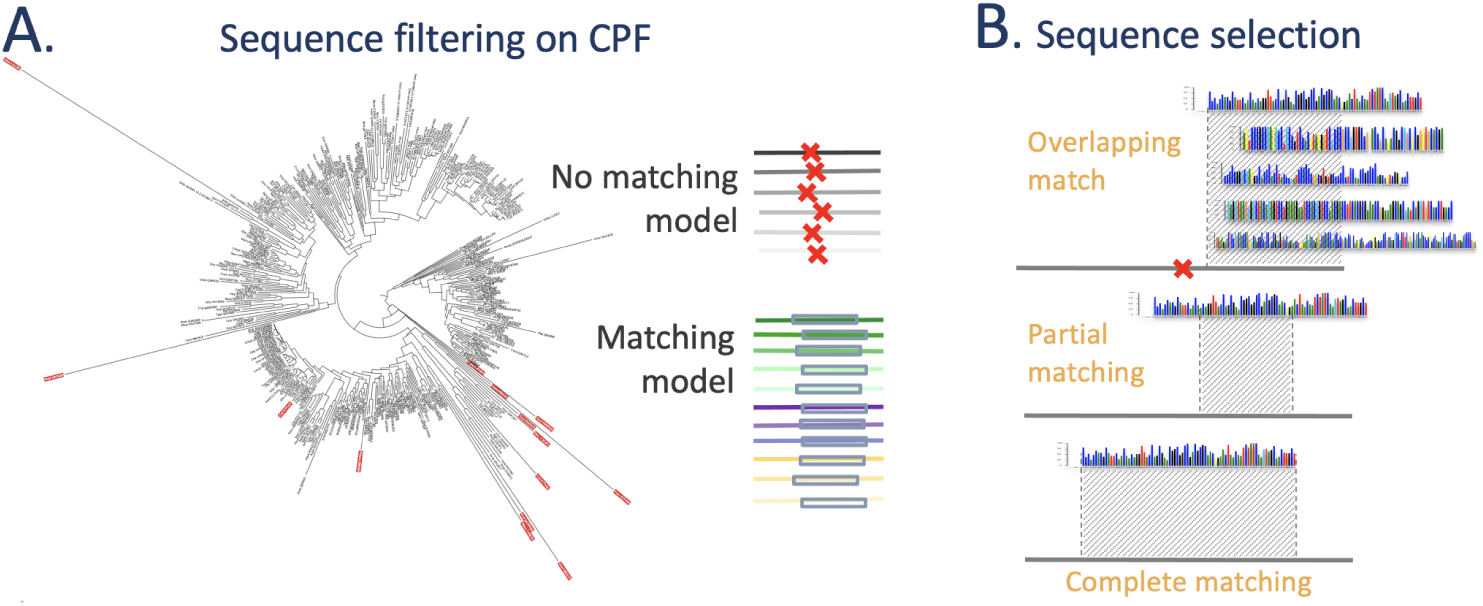
Sequence filtering and sequence selection in ProfileView pipeline. A. Phylogenetic tree of CPF input sequences where filtered sequences (that is sequences with no match of the FAD domain) are highlighted in red. Note their long branch length. B. The three types of matches between a model and a sequence used to select sequences in ProfileView. The hit, between a motif and a sequence, might involve the extreme of a sequence (top - overlapping match) or an internal region of the sequence (middle and bottom). For this latter, the motif might match the sequence only partially (middle - partial matching) or fully (bottom - complete matching).

**Figure S29:**
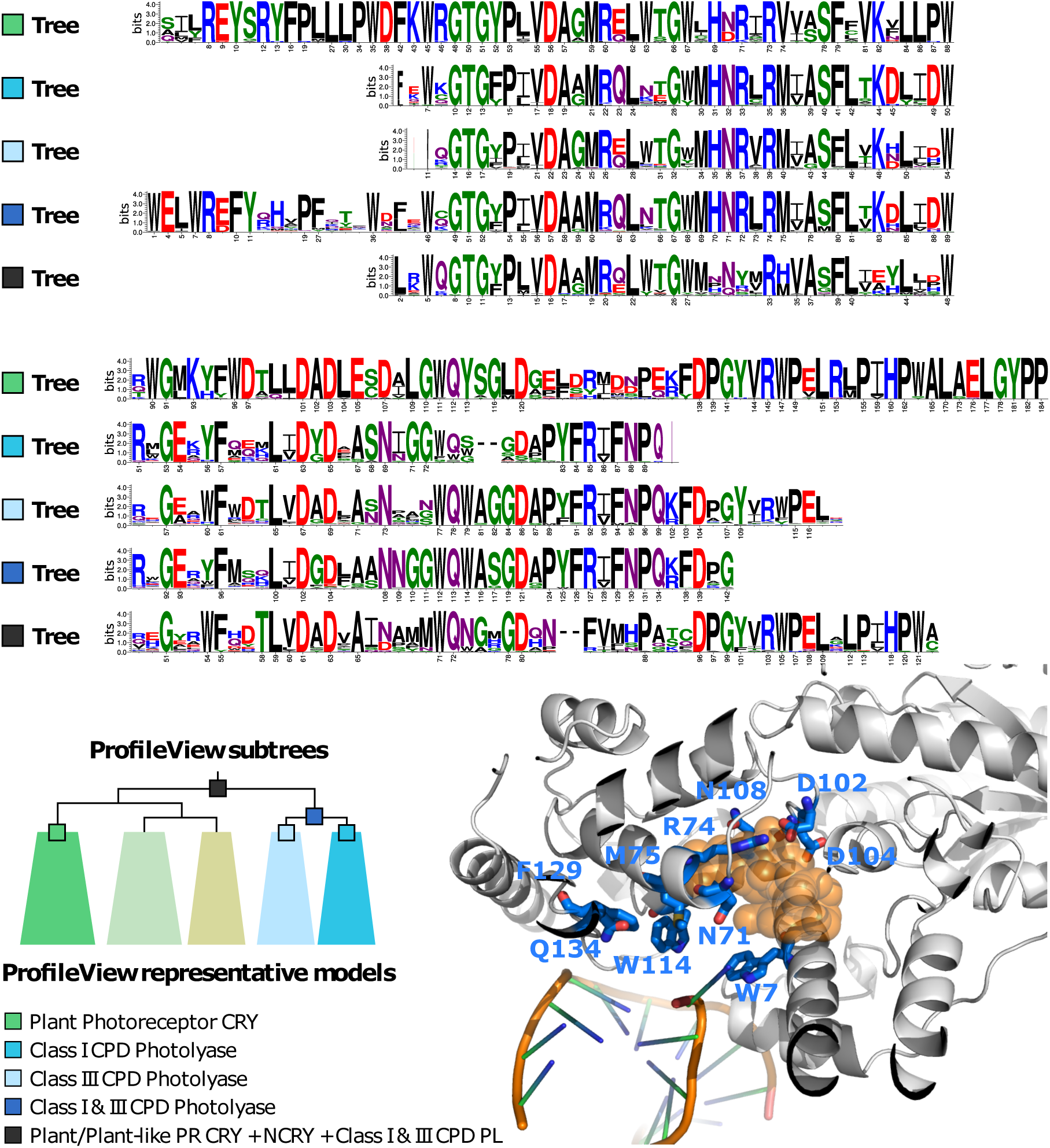
Five motifs for 5 subtrees in the ProfileView tree of the CPF family. Five representative models associated with internal nodes in the ProfileView tree are aligned. Numbered positions correspond to conserved positions belonging to the associated representative motif. The absence of the number indicates less conserved positions. The alignment has been constructed using plant PR as a template model and all others as query models. Neither plant-like PR CRY nor NCRY models were considered because no functionally characterised sequences are known for these models. The NCRY motif (associated with the beige subtree on the bottom) was not added because no functional information is available for comparison (see **Fig. S2**). The length of a motif depends on the length of the associated model, selected as best representing the sequences in a subtree. The PDB structure (1TEZ) highlights residues in interaction with DNA (W7, N71, W114 at *<* 5Å) and the FAD substrate (W7, N71, R74, M75, D102, D104, N108 at *<* 5Å). All residues highlighted in the structure have been explained in the text.

**Table S1:** Characteristics of the sets of sequences considered for classification. To be added to the list of features in Figure I: protein family name, lengths of the protein sequences, sequence identity, sequence similarity. Here, sequence length, identity and similarity are given for the entire sequence, not for the domain used to classify the protein sequences. Identities and similarities were computed using the needle command of the EMBOSS package (version 6.6.0) and took into account the full-length sequence, not just the domain portion. Note that the variability of sequence length is due to the fact that sequences might be portions of a protein in the family.

| Protein Family | # seqs | seq length | seq identity | seq similarity |
| --- | --- | --- | --- | --- |
| Cryptochrome/<br>Photolyase (CPF) | 397 | 586± 198aa | 20.96 | 30.56 |
| WW domain | 349 | 30 ± 2aa | 35.19 | 50.83 |
| Glycoside hydrolase<br>family 30 (GH30) | 1803 | 524 ± 235aa | 23.62 | 35.03 |
| HAD/β-PGM/<br>Phosphatase-like | 391 | 261 ± 101aa | 25.47 | 37.79 |
| B12-binding<br>domain containing | 273 | 549±83 | 28.33 | 40.95 |
| Methylthiotransferase | 400 | 463±65 | 29.58 | 46.64 |
| SPASM/twitch<br>domain containing | 128 | 366±36 | 38.02 | 51.96 |

**Table S2:**
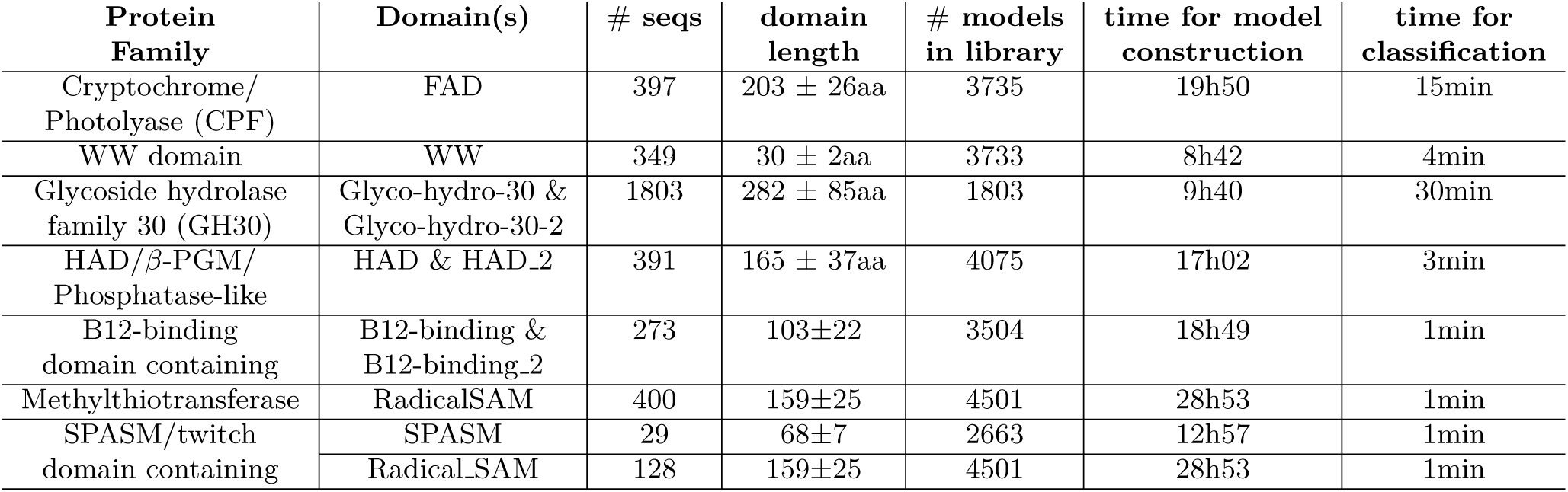
Computational time and other features. To be added to the list of features in Figure I: protein family name, domain(s) used for classification, number of sequences before filtering, domain length, number of models constructed for ProfileView analysis, time for model construction and time for classification.

**Table S3:** Validation of ProfileView on functionally characterised CPF sequences and on functionally characterised positions in CPF sequences. For each representative motif of a CPF functional class, we report the number of CPF sequences with a characterised function, the number of sequences belonging to the associated ProfileView subtree, the number of experimentally characterised positions listed in **Supplemental file** “CPF mutants used for validation.xlsx” and the number of these positions that belong to the associated motif. For transcriptional regulators, we distinguish the two ProfileView motifs (#1 and #2 in the text) and report the corresponding characterized positions in two different lines of the table. The total number of experimentally characterised sequences to be classified is reported in the line “Overall”. The sequence identified as FN corresponds to the one left white in the ProfileView tree of Fig S1.

| CPF functional class | # Experimentally characterized seqs | # Sequences in the ProfileView subtrees |  | # Experimental positions | # Positions in the ProfileView motif |
| --- | --- | --- | --- | --- | --- |
|  |  | TP | FN |  |  |
| Animal PR CRY | 4 | 4 | 0 | 15 | 13 |
| (6-4) PL | 12 | 11 | 1 | 19 | 16 |
| Transcriptional Regulator (motifs #1 and #2) | 18 | 18 | 0 | 16 | 7 |
|  |  |  |  | 16 | 5 |
| Plant PR CRY | 8 | 8 | 0 | 47 | 35 |
| Class I & III CPD PL | 15 | 15 | 0 | 33 | 26 |
| Class II CPD PL | 5 | 5 | 0 | 22 | 18 |
| CRY Pro | 2 | 2 | 0 | 11 | 10 |
| CRY DASH | 8 | 8 | 0 | 12 | 9 |
| Overall | 72 | 71 | 1 |  |  |

**Table S4:**
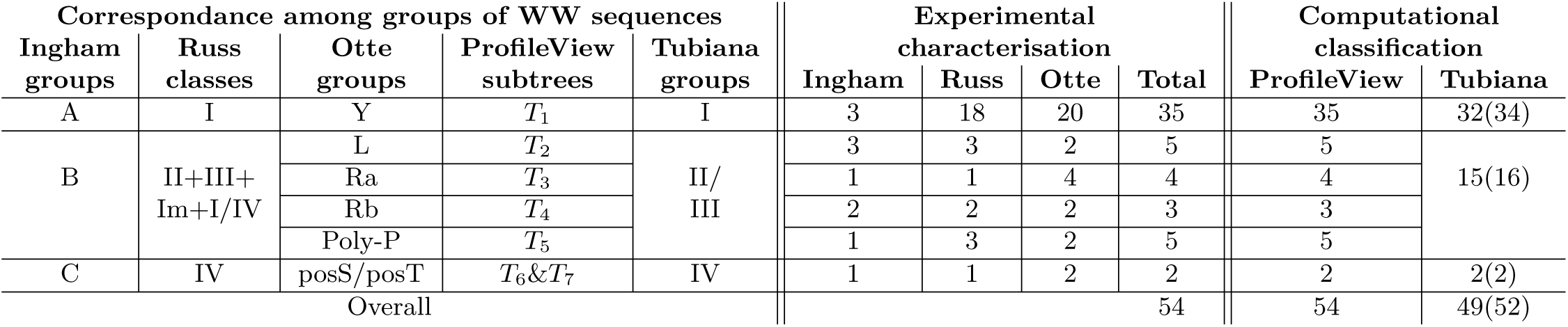
Validation of ProfileView on 54 functionally characterised sequences of the WW domain family. ProfileView classification is based on 7 distinct subtrees corresponding to the six Otte’s groups (subtrees *T*_6_ and *T*_7_ are associated to posS/posT-group in Otte), the three Ingham’s groups, and the three Russ’s classes, as indicated. This correspondance is reported in the three outer circles of the ProfileView tree in **Fig. S11**, where the names of the ProfileView subtrees are also indicated. The total number of functionally characterised sequences belonging to classes or groups in the three experimental studies is reported together with the total number of sequences characterised by the three experiments for each group (central columns). Note that some of the sequences have been experimentally characterised by more than one experiment. The right-most columns report ProfileView and Tubiana’s classifications. For Tubiana, 52 sequences among the 54 ones have been considered for classification as reported in parenthesis. Overall, Tubiana wrongly classifies 3 sequences.

**Table S5:** CAZy classification versus ProfileView classification of the 695 sequences in the GH30 family. The GH30 sequences are counted as TP whenever they make the ProfileView subtrees for the corresponding CAZy subfamily and FN whenever they wrongly appear in a subtree of another subfamily. See **Fig. S15**.

| CAZy subfamilies | # of CAZy<br>classified seq. | # of ProfileView<br>classified seq. |  |
| --- | --- | --- | --- |
|  |  | TP | FN |
| GH30-1 | 151 | 151 | 0 |
| GH30-2 | 48 | 48 | 0 |
| GH30-3 | 244 | 243 | 1 |
| GH30-4 | 60 | 60 | 0 |
| GH30-5 | 83 | 83 | 0 |
| GH30-6 | 4 | 4 | 0 |
| GH30-7 | 29 | 29 | 0 |
| GH30-8 | 63 | 63 | 0 |
| GH30-9 | 13 | 13 | 0 |
| Overall | 695 | 694 | 1 |

**Table S6:** CAZy characterisation based on EC numbers versus ProfileView classification of the 34 sequences in the GH30 family. The GH30 sequences are counted as TP whenever they belong to the ProfileView subtrees corresponding to the CAZy subfamilies GH30-1,…, GH30-9 (GH30-1: 3.2.1.45 and 3.2.1.21+3.2.1.37; GH30-2: 3.2.1.37; GH30-3: 3.2.1.75; GH30-4: 3.2.1.38; GH30-5: 3.2.1.164; GH30-6:3.2.1.136; GH30-7: 3.2.1.*; GH30-8: 3.2.1.8, 3.2.1.136, 3.2.1.8+3.2.1.136; GH30-9: 3.2.1.31) and FN when-ever they wrongly appear in a subtree of another class. We define FN only for classes of more than 3 sequences. See **Fig. S15**.

| EC numbers in CAZy<br>functional annotation | # of CAZy<br>characterised seq. | # of ProfileView<br>classified seq. |  |
| --- | --- | --- | --- |
|  |  | TP | FN |
| 3.2.1.8 | 10 | 10 | 0 |
| 3.2.1.8+3.2.1.136 | 2 | 2 | 0 |
| 3.2.1.136 | 2 | 2 | 0 |
| 3.2.1.75 | 7 | 7 | 0 |
| 3.2.1.21+3.2.1.37 | 1 | 1 | 0 |
| 3.2.1.* | 3 | 3 | 0 |
| 3.2.1.45 | 2 | 2 | 0 |
| 3.2.1.37 | 2 | 2 | 0 |
| 3.2.1.31 | 1 | 1 | 0 |
| 3.2.1.38 | 2 | 2 | 0 |
| 3.2.1.164 | 2 | 2 | 0 |
| Overall | 34 | 34 | 0 |

**Table S7:** SFLD characterisation versus ProfileView classification of the 259 HAD/*β*-PGM/Phosphatase-like sequences of the Haloacid Dehydrogenase family. The sequences are counted as TP whenever they make the ProfileView subtrees for the functional class and FP whenever they wrongly appear in a subtree of another class. We define FN only for classes of more than 3 sequences. The SFLD classes correspond to distinct ProfileView subtrees, hence the FN column contains only 0s. See **Fig. S16**.

| HAD functional classes | # of SFLD characterised seq. | # of ProfileView classified seq. |  |
| --- | --- | --- | --- |
|  |  | TP | FN |
| β-phosphoglucomutase | 43 | 43 | 0 |
| phosphonoacetaldehyde hydrolase | 36 | 36 | 0 |
| Phosphoglycolate phosphatase | 59 | 59 | 0 |
| 2-haloacid dehalogenase | 113 | 113 | 0 |
| 2-deoxyglucose-6-phosphatase | 6 | 6 | 0 |
| glycerol-3-phosphate phosphatase | 2 | 2 | 0 |
| Overall | 259 | 259 | 0 |

**Table S8:** SFLD characterisation versus ProfileView classification of the 258 B12-binding domain containing sequences of the Radical SAM family. The sequences are counted as TP whenever they make the ProfileView subtrees for the functional class and FN whenever they wrongly appear in a subtree of another class. We define FN only for classes of more than 3 sequences. See **Fig. S17**.

| B12-binding domain containing functional classes | # of SFLD characterised seq. | # of ProfileView classified seq. |  |
| --- | --- | --- | --- |
|  |  | TP | FN |
| anaerobic magnesium-protoporphyrin-IX monomethyl ester cyclase Q bacteriocin maturation | 81 | 80 | 1 |
| bacteriocin maturation | 78 | 78 | 0 |
| hopanetetrol cyclitol ether synthase | 93 | 89 | 4 |
| ladderane biosynthesis | 1 | 1 | 0 |
| valine cyclopropanase | 1 | 1 | 0 |
| OxsB-like | 1 | 1 | 0 |
| paromamine deoxygenase | 3 | 3 | 0 |
| Overall | 258 | 253 | 5 |

**Table S9:** SFLD characterisation versus ProfileView classification of the 393 Methyltiotransferase sequences of the Radical SAM family. The sequences are counted as TP whenever they make the ProfileView subtrees for the functional class and FN whenever they wrongly appear in a subtree of another class. We define FN only for classes of more than 3 sequences. See **Fig. S18**.

| Methylthiotransferase functional classes | # of SFLD characterised seq. | # of ProfileView classified seq. |  |
| --- | --- | --- | --- |
|  |  | TP | FN |
| (dimethylallyl)adenosine tRNA methylthiotransferase (MiaB-like) | 99 | 94 | 5 |
| ribosomal protein S12 methylthiotransferase (RimO-like) | 98 | 91 | 7 |
| threonylcarbamoyl-adenosine tRNA methylthiotransferase | 99 | 99 | 0 |
| CDKSRAP1 | 97 | 96 | 1 |
| Overall | 393 | 380 | 13 |

**Table S10:** SFLD characterisation versus ProfileView classification of the 29 SPASM/twitch domain containing subgroup of Radical SAM sequences based on the SPASM domain. The sequences are counted as TP whenever they make the ProfileView subtrees for the functional class and FN whenever they wrongly appear in a subtree of another class. We define FN only for classes of more than 3 sequences. See **Fig. S19**.

| SPASM/twitch domain containing functional classes | # of SFLD characterised seq. | # of ProfileView classified seq. |  |
| --- | --- | --- | --- |
|  |  | TP | FN |
| coenzyme PQQ synthesis protein E (PqqE-like) | 1 | 1 | 0 |
| Kxxx.W cyclic peptide maturase (StrB-like) | 14 | 14 | 0 |
| adenosyl-hopene transferase | 8 | 8 | 0 |
| UDP-N-acetyl-tunicamine-uracil synthase (TunB-like) | 4 | 4 | 0 |
| cytosylglucuronate decarboxylase | 0 | 0 | 0 |
| neomycin C-like epimerase | 1 | 1 | 0 |
| pcfB-like | 1 | 1 | 0 |
| Overall | 29 | 29 | 0 |

**Table S11:**
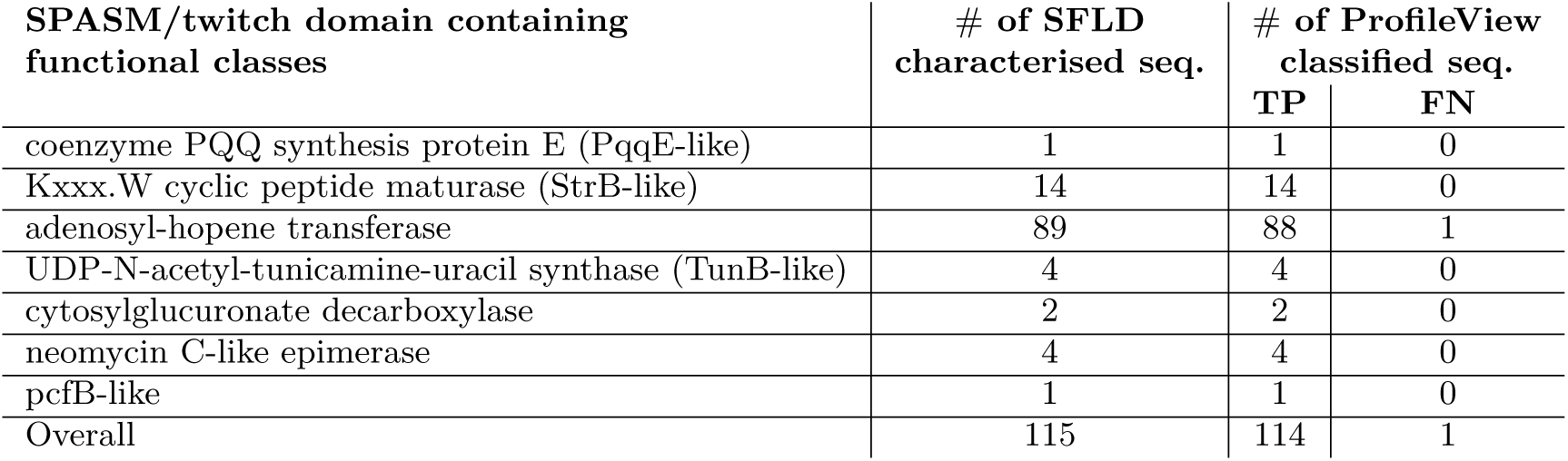
SFLD characterisation versus ProfileView classification of the 115 SPASM/twitch domain containing subgroup of Radical SAM sequences based on the Radical SAM domain. The sequences are counted as TP whenever they make the ProfileView subtrees for the functional class and FN whenever they wrongly appear in a subtree of another class. We define FN only for classes of more than 3 sequences. See **Fig. S20**.

